# Inflammasome activation in infected macrophages drives COVID-19 pathology

**DOI:** 10.1101/2021.09.27.461948

**Authors:** Esen Sefik, Rihao Qu, Caroline Junqueira, Eleanna Kaffe, Haris Mirza, Jun Zhao, J. Richard Brewer, Ailin Han, Holly R. Steach, Benjamin Israelow, Holly N. Blackburn, Sofia Velazquez, Y. Grace Chen, Stephanie Halene, Akiko Iwasaki, Eric Meffre, Michel Nussenzweig, Judy Lieberman, Craig B. Wilen, Yuval Kluger, Richard A. Flavell

## Abstract

Severe COVID-19 is characterized by persistent lung inflammation, inflammatory cytokine production, viral RNA, and sustained interferon (IFN) response all of which are recapitulated and required for pathology in the SARS-CoV-2 infected MISTRG6-hACE2 humanized mouse model of COVID-19 with a human immune system^1-20^. Blocking either viral replication with Remdesivir^21-23^ or the downstream IFN stimulated cascade with anti-IFNAR2 *in vivo* in the chronic stages of disease attenuated the overactive immune-inflammatory response, especially inflammatory macrophages. Here, we show SARS-CoV-2 infection and replication in lung-resident human macrophages is a critical driver of disease. In response to infection mediated by CD16 and ACE2 receptors, human macrophages activate inflammasomes, release IL-1 and IL-18 and undergo pyroptosis thereby contributing to the hyperinflammatory state of the lungs. Inflammasome activation and its accompanying inflammatory response is necessary for lung inflammation, as inhibition of the NLRP3 inflammasome pathway reverses chronic lung pathology. Remarkably, this same blockade of inflammasome activation leads to the release of infectious virus by the infected macrophages. Thus, inflammasomes oppose host infection by SARS-CoV-2 by production of inflammatory cytokines and suicide by pyroptosis to prevent a productive viral cycle.

## Introduction

Acute SARS-CoV-2 infection resolves in most patients but becomes chronic and sometimes deadly in about 10-20%^1-18,20,24-27^. Two hallmarks of severe COVID-19 are a sustained interferon (IFN) response and viral RNA persisting for months^1-13,20,27,28^. This chronicity is recapitulated in SARS-CoV-2 infected MISTRG6-hACE2 humanized mice^19^. Copious Interleukin (IL)-1β, IL-18 and lactate dehydrogenase (LDH) correlates with COVID-19 severity in patients, suggesting a role for inflammasome activation and pyroptosis in pathology^5-7,14-18,29^. Here, we show that human lung macrophages are infected by SARS-CoV-2. Replicating SARS-CoV-2 in these human macrophages activates inflammasomes and initiates an inflammatory cascade with a unique transcriptome, results in pyroptosis, and contributes to the downstream type-I-IFN response. Blocking either viral replication, the downstream IFN response or inflammasome activation *in vivo* during the chronic phase of the disease attenuates many aspects of the overactive immune-inflammatory response, especially the inflammatory macrophage response, and disease.

## Results

### Targeting viral replication and downstream interferon signaling ameliorates chronic COVID-19

Chronic interferon is associated with disease severity and impaired recovery in influenza infection^30^. To test whether a viral RNA-dependent type-I IFN response was a driver of chronic disease, we treated SARS-CoV-2 infected MISTRG6-hACE2 mice with Remdesivir^21-23^ and/or anti-IFNAR2 antibody (Fig. 1a) to inhibit viral replication and the IFN-response downstream of chronic infection, respectively. As control, we used dexamethasone, which reverses many aspects of immunopathology in infected MISTRG6-hACE2 mice^19^ and in patients^31^. Although Remdesivir and anti-IFNAR2 alone were partially therapeutic, combined therapy achieved more rapid weight recovery and suppression of the immune inflammatory response, especially macrophages, as effectively as dexamethasone (Fig. 1b-c, Extended data figure 1a-e), suggesting a combinatorial effect of Remdesivir and anti-IFNAR2 in chronic infection.

**Figure 1.**
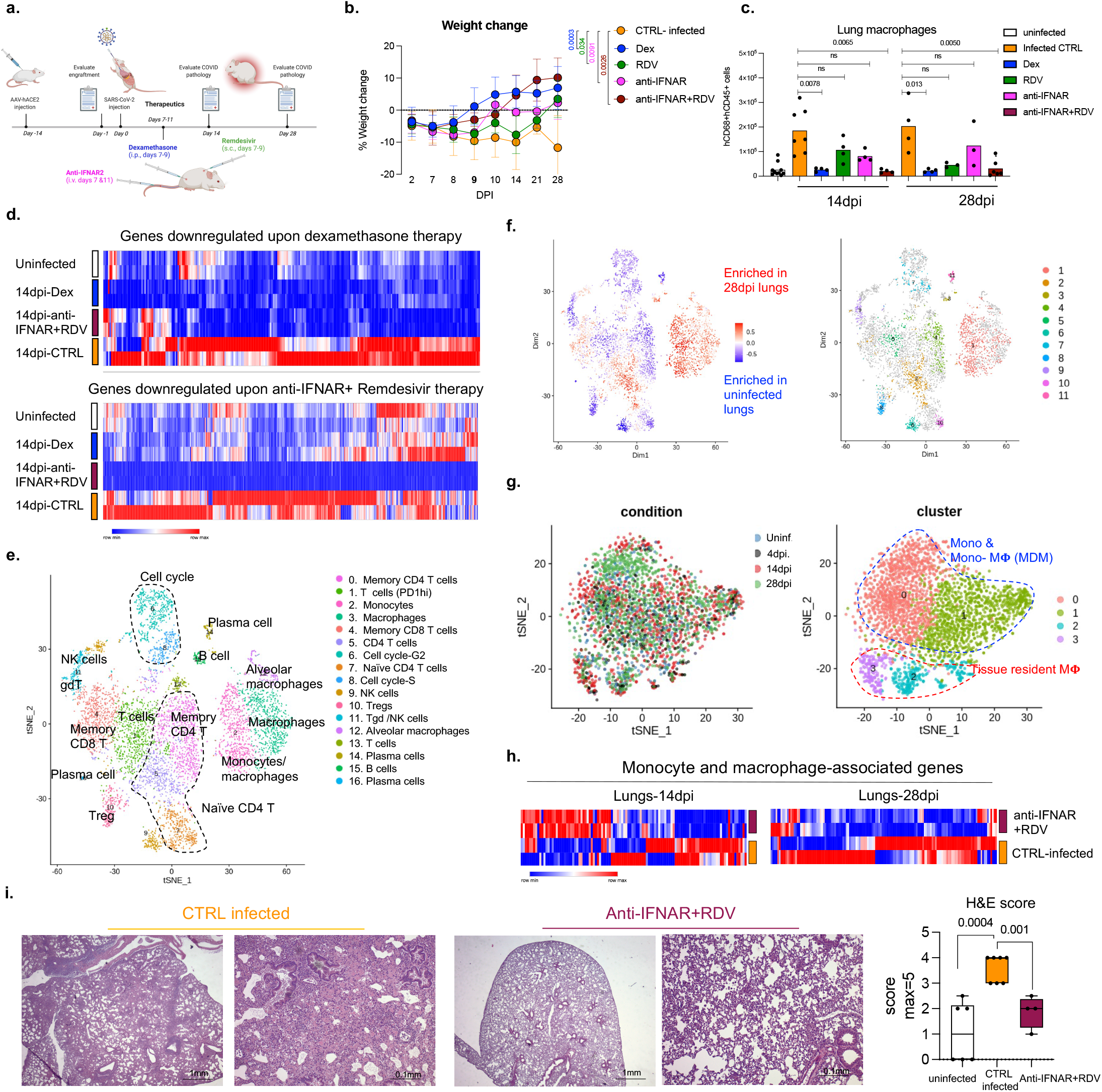
Targeting viral replication and downstream interferon signaling ameliorates chronic COVID-19. a. Therapy Schematic: Remdesivir (RDV), anti-IFNAR2 or dexamethasone (Dex). SARS-CoV-2 infected MISTRG6-hACE2 mice were treated with Dex and RDV at 7,8,9 dpi, with anti-IFNAR2 at 7, 11dpi and analyzed at 14 or 28dpi. b. Weight changes post-infection plotted as percent change compared with pre-infection weight. 28dpi: CTRL-infected n=5, Dex, anti-IFNAR2+RDV n=4, RDV, anti-IFNAR2 n=3 mice examined over at least 2 experiments. Means with SD. Unpaired, two-tailed t-test. c. Human macrophages in lungs. Uninfected: n=10; 14dpi: CTRL-infected n=7 Dex, RDV, anti-IFNAR, anti-IFNAR+RDV n=4; 28dpi: CTRL-infected, Dex n=4, RDV, anti-IFNAR n=3, anti-IFNAR+RDV n=6 mice examined over 3 experiments. Means with datapoints. Unpaired, two-tailed t-test. d. Heatmap of genes suppressed by dexamethasone or anti-IFNAR2/Remdesivir combined therapy in lungs of uninfected or infected MISTRG6-hACE2 mice (Log2, Foldchange >1; P-adj<0.05). Differential expression analysis performed with DESeq2. Statistical significance deemed using Wald test. P-adj with the Bonferroni correction. Normalized counts of duplicates visualized as min-max transformed values, calculated by subtracting row mean and diving by SD for each gene. Rows (genes) clustered by hierarchical clustering (one-minus Pearson). e. t-distributed stochastic neighbor embedding (*t*-SNE) plot of human immune cell clusters from uninfected or SARS-CoV-2 infected lungs (28dpi). Pooled duplicates. Marker genes for each cluster identified with Wilcoxon-test (Fig. S1F). Uninfected=3,655, 28dpi=3,776 cells analyzed. f. t-SNE plots highlighting differentially abundant (DA) human immune cell populations identified by DA-seq^61^. Top: Distribution of DA-populations. Red: enrichment at 28dpi, Blue: enrichment in uninfected lungs. Bottom: t-SNE plot showing 11 DA-clusters. g. *t*-SNE plots of human monocyte/macrophage clusters from 4dpi, 14dpi and 28dpi and uninfected lungs. Left: dpi, right: clusters. Cells from different conditions were combined using the integration method^62^ as described in Methods. Marker genes identified with Wilcoxon test (Seurat, Fig. S1G, H). Adjusted P-values with the Bonferroni correction. Uninfected: 438 cells, 4dpi: 336 cells, 14dpi: 793 cells, 28dpi: 1368 cells analyzed. h. Heatmap visualizing response to the combined therapy based on monocyte and macrophage associated differentially regulated genes during infection. Normalized expression of duplicates plotted as min-max of transformed values, calculated by subtracting row mean and diving by SD for each gene. Rows clustered by hierarchical clustering (one-minus Pearson). i. Representative H&E staining, box, and whisker plot of histopathological scores. Uninfected n=6, CTRL-infected n=7, anti-IFNAR2+RDV n=4 mice examined over 3 independent experiments. Whiskers: smallest (minimum) to the largest value (maximum). Box: 25^th^-75^th^ percentiles. Center line: median. Unpaired, two-tailed, t-test. Data associated with dexamethasone used here as a control have been reported^19^. Duplicates are biologically independent.

We assessed the impact of therapeutics on the lung transcriptome. Both dexamethasone and the combined therapy reversed overactive immune transcripts to uninfected animal levels (Fig. 1d, S1B,C, Table S1). The reduced transcripts were enriched for chemokine and cytokine networks (*CXCL10, CXCL8, CCL2*), inflammatory (*TLR7, NLRP3, CASP1*) and anti-viral (*MPO, OAS1, OAS2*) response, and interferon stimulated genes (ISGs) (*IFITM3, IFITM2, IRF7*) (Table S1, Fig. S1D,E), emphasizing the central role of IFN signaling and inflammatory cytokine-chemokines in chronic COVID-19. Comparison of single-cell transcriptomes of human immune cells from infected mice with their uninfected counterparts (Fig. 1e,-g) showed tissue-resident macrophages such as alveolar macrophages (AMs) activated at the peak of infection, followed by an inflammatory response with infiltrating monocytes and monocyte-derived macrophages (Fig. 1g, S1G-I, Table S2). As macrophages differentiated, they maintained their inflammatory signature and activated status throughout infection (Fig. S1G-I, Table S2). All macrophage subsets were enriched for ISGs at all timepoints (Fig. S1H). These ISGs were suppressed upon anti-IFNAR2/Remdesivir combination therapy (Fig. 1h, S1J, Table S3). Yet, key anti-viral responses such as *IFNG* primarily produced by cytotoxic T cells were spared (Fig. S1K), highlighting the selective effects of combined anti-IFNAR2/Remdesivir therapy on chronic COVID-19 pathology. Consistent with the fibrosis, seen both in patients^32-36^ and humanized mice^19^, alveolar self-renewal and differentiation programs were inhibited, resulting in the accumulation of pre-alveolar type 1 transitional cell state (PATS) program in pneumocytes^7,37-39^ that was reversed in infected MISTRG6-hACE2 mice by anti-IFNAR2/Remdesivir combination therapy, restoring self-renewal and differentiation programs (Fig. S1L). Overall, reducing chronic inflammation enhanced lung tissue recovery, and prevented transition to fibrosis seen in humanized mice^19^ (and humans^32-36^) (Fig. 1i, Extended data Fig.1f).

### SARS-CoV-2 replicates in human macrophages

To determine the cellular source of persistent viral RNA and replication, we measured genomic (gRNA) and subgenomic viral RNA (sgRNA)^40^ in lung tissue or in sorted lung epithelial cells or human immune cells from infected MISTRG6-hACE2 mice (Extended Data Fig. 2a-d). Surprisingly, epithelial cells and human immune cells had similar levels of viral RNA (Extended Data Fig. 2d). Although gRNA was abundant, we could not discern sgRNA in either cell type. We tracked infected cells in MISTRG6-hACE2 mice using a reporter strain of virus, SARS-CoV-2-mNG^41^, which encodes the fluorescent protein mNG in infected cells. By this assay, most epithelial cells in bronchioalveolar lavage (BAL) but only few total lung epithelial cells were infected with SARS-CoV-2 (Extended Data Fig. 2e). Strikingly, human macrophages were strongly mNG positive throughout disease (Fig. 2a, Extended Data Fig. 2f,g). No mouse immune cells expressed mNG (Fig. 2a, Extended Data Fig. 2f). To address whether the SARS-CoV-2 viral RNA replicates in these cells or is acquired by phagocytosis, we measured the mNG signal in human macrophages from infected MISTRG6 mice untransduced with hACE2. In these mice, epithelial cells were not infected or infected poorly with SARS-CoV-2^19,42^ (Extended Data Fig. 2h). These mice had, however, similar levels of mNG+ human macrophages as AAV-hACE2 mice, suggesting viral uptake by macrophages is independent of infected epithelial cells (Extended Data Fig. 2i). To determine whether SARS-CoV-2 replicates in human macrophages, we quantified gRNA and sgRNA^40^ in mNG+ vs mNG-epithelial or human immune cells at 4dpi or 14dpi (Fig S2A). Only mNG+, not mNG-, epithelial and immune cells had sgRNA (Fig. 2b). Second, we stained for dsRNA, diagnostic of viral replication (Fig. 2c). As expected, mNG and dsRNA were detected/colocalized in human macrophages (Fig. 2c, S3B). Third, we detected viral RdRp in human macrophages, which colocalized with a viral Spike protein supporting specificity (Fig. 2d, S3C-F). Viral RdRp and Spike were also present in human macrophages of human autopsy lungs with SARS-CoV-2 pneumonia (Extended Data Fig. 3). Thus, the mouse model observations reflected human disease. Remdesivir reduced the mNG signal and viral titers by the same amount in infected MISTRG6-hACE2 mice (Fig. 2e, Extended Data Fig. 4a). Thus, SARS CoV-2 appeared to replicate in human immune cells.

**Figure 2.**
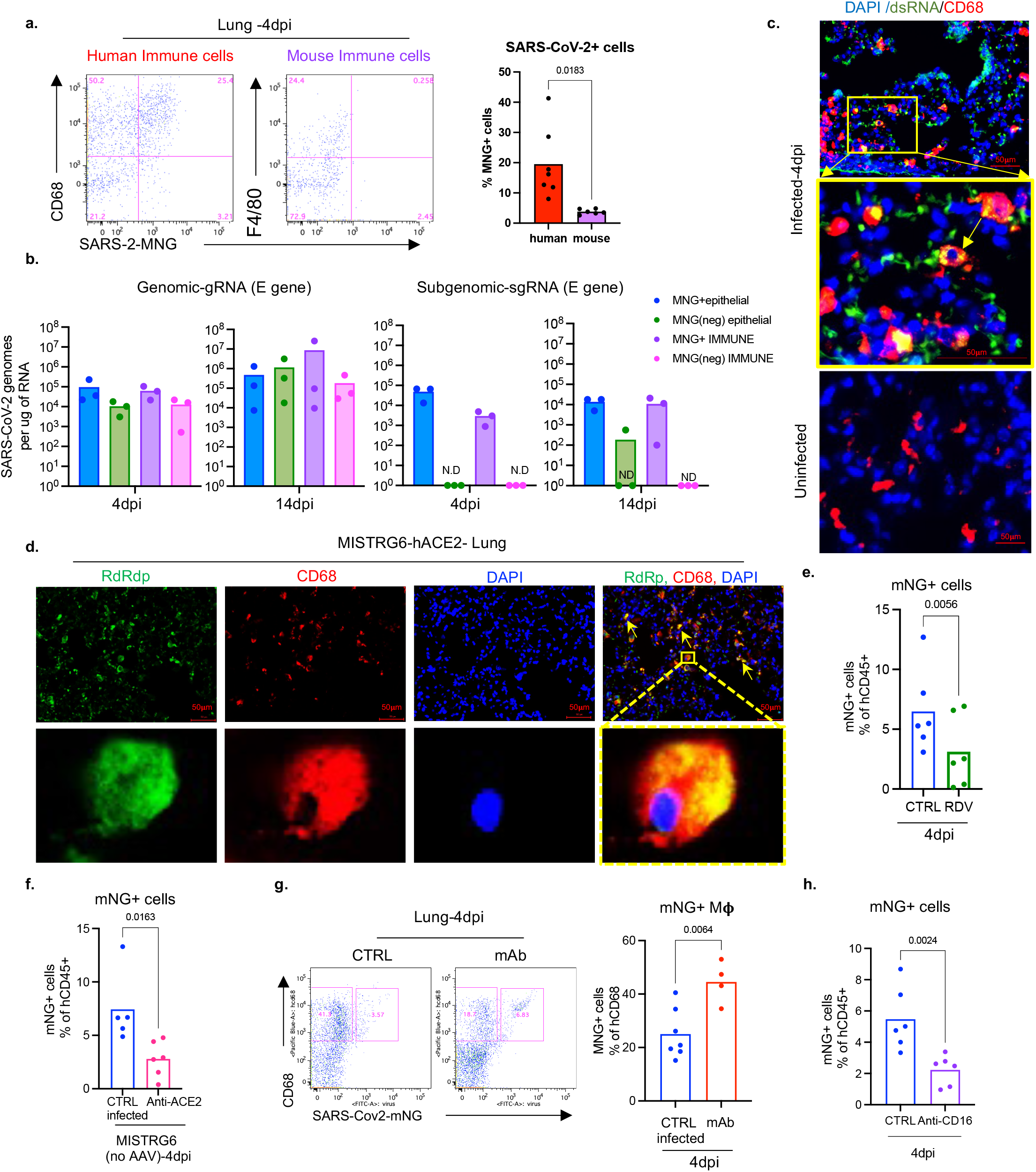
SARS-CoV-2 replicates in human macrophages. a. Representative flow cytometry plots and frequencies of mNG+ human (CD68+) or mouse (F4/80+) lung macrophages in SARS-CoV-2-mNG infected MISTRG6-hACE2 mice. Human n=7, mouse n=6 mice over at least 3 experiments. Unpaired, two-tailed t-test. b. Quantification of gRNA and sgRNA (E-gene)^40,63^ in sorted mNG+ or mNG-epithelial cells or human immune cells. N=3 mice over 2 experiments. Means with datapoints. c. Representative fluorescent microscopy images of dsRNA (rJ2), CD68 and DAPI staining in fixed lung tissues from SARS-CoV-2 infected MISTRG6-hACE2 mice. Representative of n=5 mice examined over 3 experiments. Yellow rectangle: higher magnification view of the selected area. Yellow arrow: colocalization of CD68 with dsRNA. Pseudo-colors were assigned. d. Representative fluorescent microscopy images of RdRp, CD68 and DAPI staining in fixed lung tissues from SARS-CoV-2 infected MISTRG6-hACE2 mice. Representative of n=5 mice examined over 3 experiments. Yellow arrows: colocalization of human CD68 with dsRNA. Yellow rectangle: higher magnification view of the selected area. Pseudo-colors were assigned. e. Frequencies of mNG+ human immune cells in Remdesivir treated (1-3dpi) or control MISTRG6-hACE2 mice infected with SARS-CoV-2-mNG. N=6 mice examined over 3 experiments. Means with datapoints. Paired t-test, two-tailed. f. Frequencies of mNG+ human immune cells upon ACE2 blockade (1-3dpi) in MISTRG6 (no AAV) mice infected with SARS-CoV-2-mNG. CTRL-infected n=5, anti-ACE2 treated n=6 mice examined over 2 experiments. Means with datapoints. Paired t-test, two-tailed. g. Representative flow cytometry plots and frequencies of mNG+ macrophages in infected MISTRG6-hACE2 mice treated with monoclonal antibodies (mAb)^19,45,64^ at 35hpi. CTRL-infected n=7, treated n=4 mice examined over 2 experiments. Means with datapoints. Unpaired, two-tailed t-test. h. Frequencies of mNG+ human immune cells in MISTRG6-hACE2 mice after CD16 blockade (Abcam, 2dpi). N=6 mice examined over 3 experiments. Means with datapoints shown. Paired t-test, two-tailed.

### SARS-CoV-2 infection is mediated by ACE2 and CD16

The ACE2 receptor utilized by SARS-CoV-2 to infect lung epithelium can be expressed in macrophages^43^. We measured ACE2 expression by flow cytometry and immunofluorescence staining in mouse epithelial cells and human lung macrophages (Extended Data Fig. 4b-f). Human lung macrophages from both MISTRG6 and MISTRG6-hACE2 mice, but only epithelial cells from MISTRG6-hACE2 mice expressed human ACE2 (Extended Data Fig. 4b-e). Interestingly, ACE2 expression was higher in both infected (mNG+) human macrophages and epithelial cells. We treated SARS-CoV-2 infected MISTRG6 mice with a blocking antibody against human ACE2. In these mice only, SARS-CoV-2 infects epithelial cells poorly^19,42^ as the mice did not receive AAV-ACE2 and only human macrophages express human ACE2 (Extended Data Fig. 2h). ACE2 blockade significantly diminished infected human macrophages (Fig 2f), suggesting ACE2 can mediate viral entry in human lung macrophages.

Antibodies can also mediate viral uptake by macrophages (e.g., Dengue virus^44^). To test the role of antibody-mediated viral entry to macrophages, we treated infected mice with monoclonal antibodies (mAb)^45^ against SARS-CoV-2-Spike protein early (35hpi) when effects of endogenous antibodies are minimal or late (7dpi) (Extended Data Fig. 4g). Indeed, mAb treatments increased infected lung macrophages (Fig. 2g, Extended Data Fig. 4h). Immune cells express a wide range of surface Fcγ receptors (FcγRs) which interact with the Fc moiety of antibodies. These interactions lead to multiple protective or pathological effector functions^44,46^. COVID-19 severity correlates with high serum IgG levels and specific IgG-Fc structures and interactions^47-49^. One such Fc-interaction is mediated by CD16, expressed at high levels in mNG+ macrophages. We treated mice early (2dpi, low antibody levels) as proof of concept, or late (7dpi and 11dpi, high antibody levels) as a possible therapeutic with anti-CD16 antibody. Anti-viral antibody levels in lung tissue were sufficient to mediate viral uptake and positively correlated with mNG levels at 4dpi (Extended Data Fig. 4g,i). With dosing optimized, CD16 blockade which did not alter distribution of macrophages, however resulted in significantly fewer infected human macrophages at both timepoints (Fig. 2h and Extended Data Fig. 4j).

To elucidate whether viral replication products are the result of bona-fide infection, we cultured bone-marrow derived macrophages (BMDM) with SARS-CoV-2 *in vitro*. Indeed, SARS-CoV-2 were taken up by BMDM and replicated in these cells as measured by mNG signal (Extended data Fig. 5a) and high levels of sgRNA (Extended data Fig. 5b). This was true for multiple types of macrophages (Extended data Fig. 5c). As *in vivo, in vitro* macrophage infection enhanced by antibodies (convalescent plasma or mAbs) was reduced by CD16, ACE2, or RdRp blockade (Extended data Fig. 5d, e). SgRNA levels in these macrophages were also reduced by these treatments (Extended data Fig. 5b), further supporting a role for both ACE2 and CD16 in viral uptake and RdRp in viral replication. SARS-CoV-2 infection in human macrophages was not productive or produced very little as indicated by undetectable infectious virus, titered in culture, from sorted immune cells from infected mice at 4dpi and *in vitro* infected macrophages at 48hpi (Extended data Fig. 5f-h).

### Infected macrophages have a unique transcriptional signature

We next determined the consequences of infection of human macrophages by SARS-CoV-2. Infected macrophages preferentially produced CXCL10, a chemokine which recruits many types of immune cells (Fig. 3a), but not TNF. Like mNG positivity itself, CXCL10 production by human macrophages was also enhanced by antibodies and inhibited by Remdesivir, also reflected in serum levels and *in vitro* (Fig. 3b,c, S3A-C). Thus, we used CXCL10 as a proxy for SARS-CoV-2-infected macrophages and determined a unique transcriptional signature enriched for genes encoded by tissue-resident macrophages, in particular AMs^50^ (*APOC1, MRC1, ALOX5AP, FABP5, INHBA*), chemokines of interstitial macrophages (*CCL18, CCL3, CCL7, CCL8, CCL20, CXCL8*), inflammatory cytokines (*IL1A, IL18, IL27*), complement genes (*C1QA, C1QB*) and ISGs (*ISG20, IFI27*) (Fig. 3d, S3D-G, Table S4). Further flow cytometric characterization of mNG+ cells also confirmed enrichment for CD16+ AMs, which produced more CXCL10 (Fig. 3e, S3H). In line with our findings, CD14^hi^CD16^hi^ cells and AMs enriched with viral RNA in autopsy lungs of COVID-19 patients^7,20^ also had distinct transcriptomes which were largely recapitulated in what we construe as CXCL10-associated genes (CXCL11, CCL18, CCL8, ISG15, CD83). Interestingly, this strong network of CXCL10 specific gene signature was no longer restricted to AMs later in infection as different macrophage subsets continuously differentatiate, evident in high *IL7R* expression by developing lung macrophages^50^ (CXCL10+ and AM) at all timepoints (Fig. 1g, 3d, S3I).

**Figure 3.**
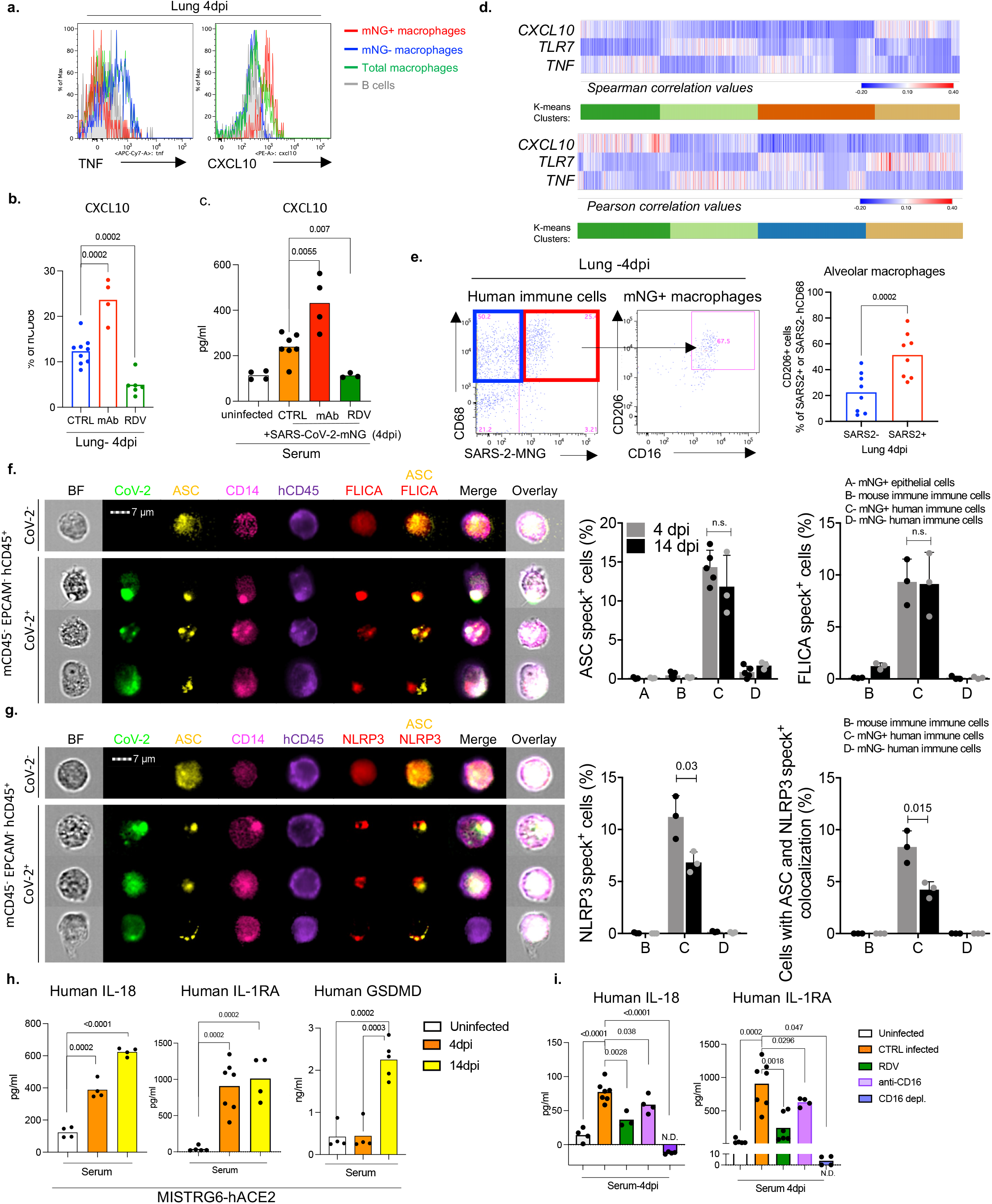
SARS-CoV-2 infection of human macrophages activates inflammasomes and pyroptosis. a. CXCL10+ or TNF+ human macrophages. Representative of n=6 mice over 3 experiments. b. CXCL10+ lung macrophage frequencies upon mAbs or Remdesivir therapy. CTRL-infected n=9, mAb n=4, RDV n=6 mice over at least 2 experiments. Means with datapoints. Unpaired, two-tailed-t-test. c. Serum CXCL10 levels upon mAb or Remdesivir therapy. Means with datapoints. Uninfected, mAb n=4; CTRL-infected n=7, RDV n=3 mice examined over 2 experiments. Unpaired, two-tailed-t-test. d. Correlation (Pearson and Spearman) of each gene with *CXCL10, TNF* or *TLR7* in human lung monocytes and macrophages. K-means clustering. P-values: T-distribution with length(x)-2 degrees of freedom or algorithm AS 89 with exact = TRUE. Two-tailed. e. Representative flow cytometry plots: Alveolar macrophage frequency within mNG+ or mNG-macrophages. N=8 mice examined over 4 experiments. f. ASC speck visualization/quantification and colocalization with active caspase-1 (FLICA) in mNG+ or mNG-human immune cells from MISTRG6-hACE2 mouse lung. Cells sorted based on Fig.S2A. 1000-cells analyzed/per condition. ASC+specks: 4dpi n=3(A), 5(B-D); 14dpi n=3 mice, FLICA: n=3 mice examined over at least 2 experiments. Means with datapoints, SD. Unpaired, two-tailed-t-test. g. ASC speck visualization/quantification and colocalization with NLRP3 oligomerization in sorted mNG+ or mNG-human lung immune cells. 1000-cells analyzed/per condition. N=3 mice over 2 experiments. Means with datapoints, SD. Unpaired, two-tailed-t-test. h. Serum IL-18, IL-1RA and GSDMD levels. IL-18: n =4 mice examined over 2 experiments. IL-1RA: Uninfected n=5, 4dpi n=7, 14dpi n=4 mice examined over 3 experiments. GSDMD: uninfected, 4dpi n=4, 14dpi n=5 mice over 3 experiments. Means with datapoints. Unpaired, two-tailed t-test i. Serum IL-18 and IL-1RA levels in mice treated with CD16-blocking (Abcam) or depleting (ThermoFisher), antibodies or Remdesivir. IL-18: uninfected, CD16-blocking, CD16-depletion n=4, CTRL-infected n=7, RDV n=3; IL-1RA: uninfected n=5, CTRL-infected n=7, RDV n=6, CD16-blocking, CD16-depletion n=4 mice examined over at least 2 experiments. Means with datapoints. Unpaired, two-tailed t-test.

### SARS-CoV-2 infection activates inflammasomes

Morphological analysis of sorted mNG+ cells revealed the appearance of membrane bubbles, a characteristic of pyroptosis and prompted us to investigate inflammasome activation as part of the inflammatory cascade initiated by infection. Inflammasomes are dynamic multiprotein complexes in which specific NOD-like receptors (NLRs) and adaptor molecules are assembled to activate caspases, the central effector proteins. We sorted mNG+ and mNG-human immune cells, mNG+ epithelial cells, and mouse immune cells (Fig.S2A) and assayed for sensors, adaptors, and effectors of the inflammasome pathway. First, focusing on adaptor molecule apoptosis-associated speck-like protein containing a CARD (ASC) as the common adaptor molecule with a pivotal role in inflammasome assembly and activation, we found that infected (mNG+) human cells exclusively showed significant inflammasome activation, quantified by ASC speck formation (Fig.3f, Extended Data Fig. 6a,b). ASC specks co-localized with both NLRP3 and active caspase-1 (visualized by fluorochrome-labeled inhibitor of caspases assay (FLICA)) (Fig 3f, g, Extended Data Fig. 6a-d). Inflammasome activation in infected human macrophages was sustained during disease (4-14dpi; Fig 3f,g, Extended Data Fig. 6c).

Once inflammasome complexes are formed, active caspase-1 cleaves and proteolytically activates the pro-inflammatory IL1-family cytokines, IL-1β and IL-18, typically elevated and characteristic of severe COVID-19 in patients. IL-18 levels in blood and lungs were significantly elevated in SARS-CoV-2 infected mice and correlated well with proportions of infected (mNG+) macrophages (Fig. 3h, Extended Data Fig. 6e). Although IL-1β levels in serum in vivo were not detectable, we measured IL-1RA. This specific receptor antagonist, induced by IL-1β, served as a proxy of IL-1β and it paralleled enhanced IL-18 levels and correlated with mNG+ cells (Fig 3h, Extended Data Fig. 6f).

Finally, we assayed for pyroptosis by detecting LDH and gasdermin D (GSDMD) in serum. GSDMD, a substrate of active caspase-1 and pore-forming executer of pyroptosis, and LDH, released by pyroptosis, were particularly enriched in serum of infected mice at late timepoints (14dpi-Fig3h, Extended Data Fig. 6g), further supporting continuous inflammasome activation during infection. In addition, infected lung macrophages showed higher incorporation of a small fixable dye (Zombie Aqua) that enters dying cells with a compromised cell membrane, consistent with the pore-forming function of GSDMD and pyroptosis (Extended Data Fig. 6h).

All aspects of inflammasome activation were also recapitulated in vitro when BMDM were infected in vitro with SARS-CoV-2. Active caspase-1 in infected BMDM was dependent on viral replication was inhibited by Remdesivir (Extended Data Fig. 7a). High levels of inflammasome products IL-18, IL-1β, and IL-1RA (in response to IL-1β) and two measures of pyroptosis, GSDMD and LDH, were detected at high levels in supernatants of infected BMDM were also inhibited by remdesivir (Extended data Fig. 7b-f). In vitro infected cells also had higher incorporation of Zombie-Aqua consistent with pyroptosis (Extended Data Fig. 7g).

### Viral entry or replication impacts the inflammatory profile of macrophages

To determine the role of viral infection on the inflammatory macrophage response, we first blocked viral entry and replication in vivo and measured inflammatory cytokines and chemokines. Blocking viral entry (CD16 or ACE2 blockade) or inhibiting viral replication (Remdesivir) all reduced IL-18, IL-1RA, and CXCL10 levels, paralleling mNG levels (Fig 3i, Extended Data Fig. 8a-e). Depletion of CD16+ cells in vivo (Extended Data Fig. 8f,g) resulted in complete loss of IL-18 and IL-1RA in serum consistent with the concept that viral replication and inflammasome activation occurred mainly in myeloid cells (Fig 4i). On the other hand, mAbs which promoted viral infection of human macrophages (Fig. 3i) enhanced systemic IL-18, IL-1RA, and CXCL10 (Extended Data Fig. 10H-K) particularly in early disease. Nonetheless, despite changes in levels of the inflammatory cytokines and chemokines, neither mAbs nor CD16 blockade impacted lung pathology potentially because of the conflicting role of these antibodies on viral titers vs. inflammation (Extended Data Fig. 8l, m). In line with the in vivo studies, IL-18, IL-1β, IL-1RA, and CXCL10 levels were also reduced in supernatants of in vitro infected BMDM upon CD16, ACE2 or RdRp inhibition, again paralleling the reduced viral replication inferred from mNG levels (Fig. S2C, Extended Data Fig. 5e, 8n-q).

**Figure 4.**
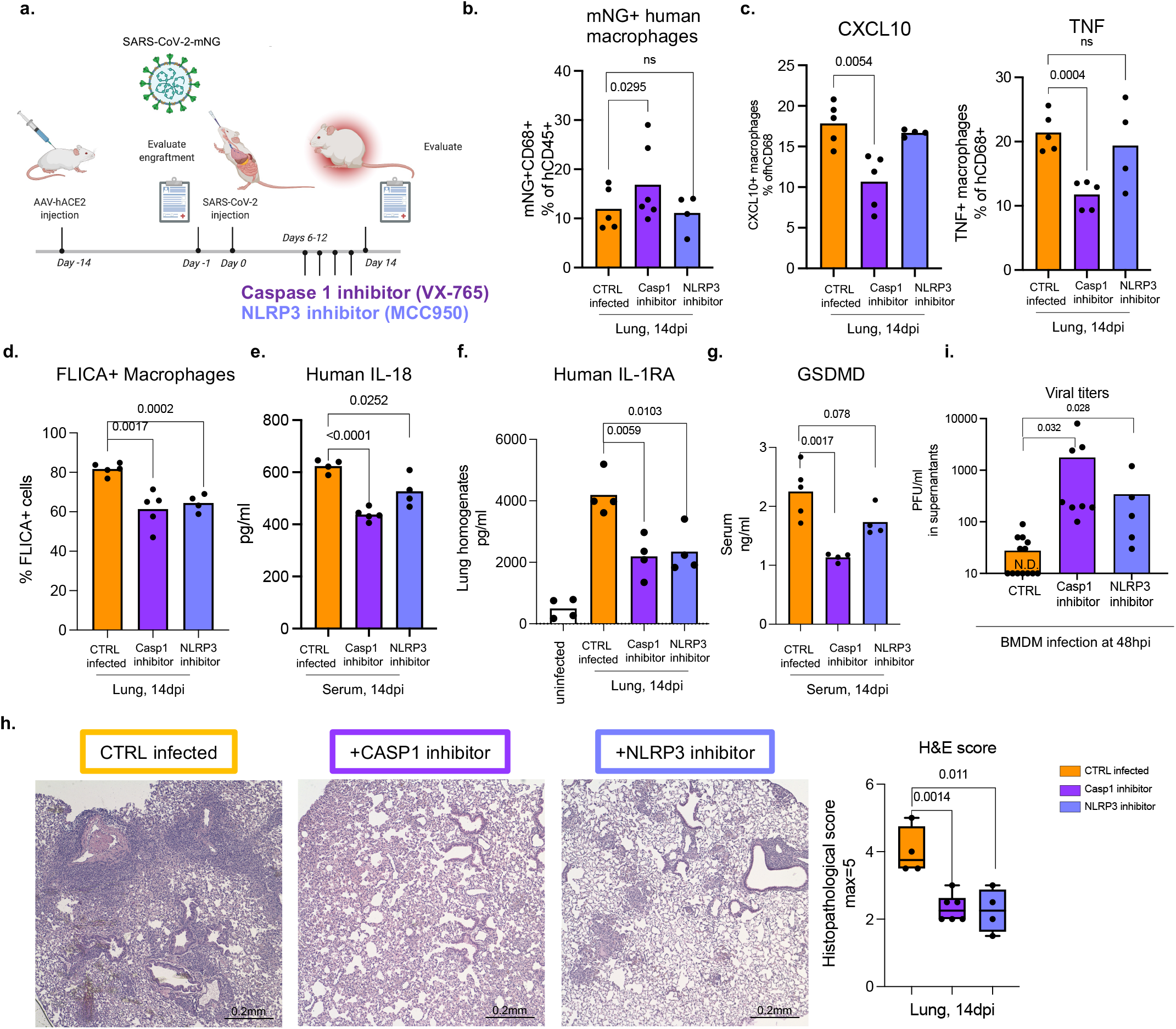
Inflammasome inhibition ameliorates inflammation and disease in infected MISTRG6-hACE2 mice. a. Schematic of inflammasome inhibition *in vivo*. SARS-CoV-2 infected MISTRG6-hACE2 mice treated with caspase-1 or NLRP3 inhibitors, 6-12dpi. b. Frequencies of mNG+ human immune cells upon inflammasome inhibition. CTRL-infected n=5, caspase-1 inhibitor n=6, NLRP3 inhibitor n=4 mice examined over at least 2 experiments. Means with datapoints. Paired, two-tailed t-test. c. Frequencies of CXCL10+ or TNF+ human lung macrophages upon inflammasome inhibition. CTRL-infected n=5, caspase-1 inhibitor n=5, NLRP3 inhibitor n=4 mice examined over at least 2 experiments. Means with datapoints. Unpaired, two-tailed t-test. d. Quantification of active caspase-1 in mNG+ human macrophages upon inflammasome inhibition. CTRL-infected n=5, Casp1-inhibitor n=5, NLRP3-inhibitor n=4 mice examined over at least 2 experiments. Means with datapoints. Unpaired, two-tailed t-test. e. Serum human IL-18 levels upon inflammasome inhibition. CTRL, NLRP3: n=4, Casp1 n=5 mice examined over 2 experiments. Means with datapoints. Unpaired, two-tailed t-test. P<0.0001=1.00114×10^−5^. f. Human IL-1RA levels in lung homogenates upon inflammasome inhibition. N=4 mice examined over 2 experiments. Means with datapoints. Unpaired, two-tailed t-test. g. Serum GSDMD levels upon inflammasome inhibition. CTRL-infected n=5, Casp1, NLRP3 inhibitors n=4 mice examined over 2 experiments. Means with datapoints. Unpaired, two-tailed t-test. h. Box and whisker plot of histopathological scores upon inflammasome inhibition. CTRL-infected n=4, Casp1-inhibitor n=6, NLRP3-inhibitor n=4 independent mice over at least 2 experiments. Whiskers: smallest (minimum) to the largest value (maximum). Box: 25^th^-75^th^ percentiles. Center line: median. Unpaired, two-tailed t-test. i. Viral titers from supernatants of BMDM infected with SARS-CoV-2-mNG *in vitro* and treated with Casp1 or NLRP3 inhibitors. CTRL-infected: n=13, Casp1-inhibitor-: n=8, NLRP3 inhibitor: n=5 independent datapoints collected over 3 experiments. Means with datapoints. Unpaired two-tailed t-test.

### Inflammasome inhibition ameliorates inflammation and disease

Finally, to assess the causal role of NLRP3 and caspase-1 activation in inflammasome mediated inflammation and disease, we treated mice with caspase-1 and NLRP3 inhibitors (Fig.4a). As expected, the proportion of infected cells did not diminish, but the inflammatory profile of these cells and other lung macrophages was drastically attenuated (Fig. 4b). In inhibitor treated mice, mNG+ cells produced less CXCL10 which was also reflected in reduced serum levels (Fig. 4c, Extended Data Fig. 9b-c). Lung macrophages (mNG-) also produced less TNF (Fig. 4b, Extended Data Fig. 9a,b). Overall, inhibitor treated mice had lower levels of caspase-1 activation, IL-18, IL-1RA and GSDMD levels (Fig. 4d-g). The cumulative decrease in proinflammatory cytokines and chemokines upon inflammasome inhibition reversed the immune-pathological state of the lung, measured by scoring of lung histopathology (Fig. 4h). Inflammasome inhibition reduced immune cell infiltration and enhanced tissue recovery to homeostasis in lungs despite persistently high levels of mNG+ human immune cells in lungs.

Caspase-1 and NLRP3 inhibitors blocked inflammasomes *in vitro* but did not impact macrophage infection, measured as mNG+ macrophage frequency, and reduced the inflammatory response to infection (Extended Data Fig. 9d). All parameters of inflammasome activation: active caspase-1, IL-1β, IL-18, GSDMD and LDH were significantly reduced upon caspase-1 and NLRP3 inhibition *in vitro* (Extended Data Fig. 9e-i). Consistent with decreased pyroptosis, inflammasome blockade significantly reduced Zombie Aqua positive cells (Extended Data Fig. 9j). As seen *in vivo*, in*-vitro* infected BM macrophages produced less CXCL10 and IL-1RA (Extended Data Fig. 9k,l).

Finally, we tested whether inflammasome activation translated to any changes in infectious virus levels. We therefore first measured viral titers in lungs of caspase-1 inhibitor treated mice. Indeed, caspase-1 treated mice had higher viral load at 14dpi *in vivo* (Extended Data Fig. 9m). Given that, reduced inflammatory response could result in deficient viral clearance, we infected macrophages in vitro and treated them with caspase-1 or NLRP3 inhibitors to test the direct effect of inflammasome activation on infectious virus. Analysis of supernatants of these cultures showed that inhibitor treated cells produce substantially higher amounts of virus than the uninhibited controls (Fig. 4i, Extended Data Fig. 9n). Thus, the activation of inflammasomes in infected macrophages plays two protective functions-attenuates virus production and signals infection to the immune system by releasing inflammatory cues to recruit and activate more immune cells at the site of infection.

Overall, these findings suggest that infection of macrophages by SARS-CoV-2 activates inflammasomes and drives pyroptosis. Pyroptosis interrupts the viral replication cycle and prevents viral amplification; in parallel it releases immune cell activators and recruiters. Viral RNA/PAMPs and proinflammatory cytokines released from these cells likely shape the hyperinflammatory macrophage response sustained by infiltrating monocytes and MDMs and drive immunopathology.

## Discussion

The MISTRG6 COVID-19 model faithfully reflects many of the chronic immunoinflammatory features of the human disease such as chronic viral RNA, IFN-response, and inflammatory state in macrophages^19^. Overall, our mechanistic study of this model defines a cascade of events, which initiates with infection of lung macrophages generating replicative intermediates and products including RdRp, dsRNA, sgRNA. SARS-CoV-2 replication activates an inflammatory program with activation of inflammasomes, production, and release of inflammatory cytokines and chemokines, and pyroptosis. We established all steps of inflammasome activation by visualizing ASC oligomerization, colocalization with active caspase-1 and NLRP3, maturation of inflammasome-mediated cytokines IL-1β and IL-18, and pyroptosis assayed by GSDMD and LDH release. Inhibitors of both caspase-1 and NLRP3 blocked the downstream aspects of inflammasome activation and the inflammatory cascade both *in vivo* and *in vitro*. More importantly, targeting inflammasome mediated hyperinflammation or combined targeting of viral replication and the downstream interferon response in the chronic phase of the disease prevented immunopathology associated with chronic SARS-CoV-2 infection *in vivo*.

Unlike epithelial cells, infected macrophages produce little virus. Strikingly however, inhibition of the inflammasome pathway led to a substantial increase in infectious virus produced by infected macrophages; though the degree these macrophages contribute, if at all, to high titers of virus production is unclear. Remarkably, inflammasome activation denies the virus the opportunity to replicate productively in these sentinel immune cells, and instead broadcasts inflammatory signals which inform the immune system of the infectious menace. While this is potentially beneficial, excessive inflammation that occurs through this mechanism coupled with the dysregulated interferon response may be the key factor which leads to the excessive inflammation that typifies chronic COVID-19^2,5,51-54^. Indeed, attenuation of the inflammasome *in vivo* blocks the inflammatory infiltrates in the lungs of infected mice *in vivo*. We speculate that, by contrast, an early interferon response, as may occur in the majority of patients who rapidly clear infection, and in the acute mouse models of infection where human immune cells that can be infected are not present, leads to viral elimination before this inflammatory chain reaction can occur.

Viral RNA and particles can be detected by a variety of innate immune sensors. Inflammasome sensor NLRP3 is both upregulated and activated by replicating SARS-CoV-2. NLRP3 inflammasome can directly sense viral replication/RNA or can rely on other viral RNA sensors like MDA5 or RIG-I^55-57^. Loss of IL-18/IL-1β production upon Remdesivir treatment in our studies strongly suggest viral replication is involved. Recent reports have also identified a possible role for NLRP3 driven inflammasome activation in SARS-CoV-2 infected myeloid cells in post-mortem tissue samples and PBMC^58^. Although many candidates have been proposed (lytic cell death upon infection, N protein^59^, ORF3A^60^), the exact mechanism of NLRP3 activation is still poorly understood^29^. Activation of other NLRs may also contribute to the process, as inhibition of caspase-1 was stronger than NLRP3 alone. Finally, there may be other mechanisms that enhance SARS-CoV-2 infection or the downstream inflammatory response in human macrophages that are unexplored here.

A role for inflammasome driven hyperinflammation in COVID-19 pathophysiology in patients is now recognized ^5-7,14-18^. Targeting inflammasome pathways in patients may provide alternative therapeutic options for resolving chronicity in COVID-19. However, the increased virus production seen upon inflammasome blockade could pose a significant risk to the benefit of wholesale inhibition of the pathway. The findings from our study and its implications provide alternative therapeutic avenues to be explored in the clinic and may guide novel therapeutic developments and prompt clinical trials to investigate combinatorial therapies that target viral RNA, inflammasome activation or its products and sustained IFN response.

## Data Availability

All data that support the findings of this study are available within the paper, its supplementary Information files, and source files. The data supporting this publication and presented as part of the supplementary information is available at Figshare.com under the project entitled, “Viral replication in human macrophages enhances an inflammatory cascade and interferon driven chronic COVID-19 in humanized mice” (https://doi.org/10.6084/m9.figshare.19401335). All 10x Genomics single cell RNA sequencing and bulk RNA sequencing data that support the findings of this study are deposited in the Gene Expression Omnibus (GEO) repository with accession codes GSE186794 and GSE199272.

## MAIN TEXT STATEMENTS

## Acknowledgements

The generation of the original MISTRG6 model was supported by the Bill and Melinda Gates Foundation. We thank G. Yancopoulos, D. Valenzuela, A. Murphy, and W. Auerbach at Regeneron Pharmaceuticals who generated, in collaboration with our groups, the individual knock-in alleles combined in MISTRG. We thank M. Chiorazzi, I. Odell and W. Philbrick and all the other members of the Flavell lab for discussions and comments; J. Alderman, B. Cadugan, and E. Hughes-Picard for administrative assistance; B. Cadugan and E. Eynon for extensive manuscript editing; P. Ranney, C. Hughes for mouse colony management; D. Urbanos for human CD34+ cell isolation; L. Devine and E. Manet for help with cell sorting; R. Filler for help with viral stocks and cell lines. E. Sefik is a HHMI Fellow of the Damon Runyon Cancer Research Foundation (DRG-2316-18). This work was funded by the Howard Hughes Medical Institute (RAF, MCN, and AI). This study was also supported in part by awards from National Institute of Health grants, R35 GM142687 (YGC), R01AI157488 (AI), F30CA239444 (ES), 2T32AI007517 (BI), R01 AI061093(EM), AI118855(EM), CA016359 (EM), K08 AI128043 (CBW), U01 CA260507 (SH), R01 GM131642 and GM135928 (RAF, YK), Burroughs Wellcome Fund (CBW), Patterson Foundation (CBW), Fast Grant from Emergent Ventures at the Mercatus Center (AI, CBW), Mathers Foundation (AI, CBW, EM), the Ludwig Family Foundation (AI, CBW), Harvard Lemann Brazil Research Fund (JL), and the Rita Allen Foundation (YGC).

## Author contributions

E.S conceived the project, performed experiments, analyzed the data, and wrote the manuscript. R.Q and J.Z. performed bioinformatics analysis. C. J. performed imaging flow cytometry experiments for characterization of the NLPR3 inflammasome. E.K. prepared samples for histopathological assessment and performed all immunofluorescence staining. H. M. performed histopathological assessment of lung pathology, quantification of immunofluorescence staining and offered essential conceptual insight in interpreting lung pathology. B.I. helped establish the model in Biosafety Level 3. M.N. provided monoclonal antibodies used in the study. H.N.B helped with tissue preparation and immunofluorescence staining. S.V. provided help with IL-1 quantification protocols. Y.G.C. provided protocols and insight on dsRNA staining. J.R.B., A.H., H.S., S.H., A. I., E.M., M.N., J.L., C. W., Y.K. offered vital conceptual insight, contributed to the overall interpretation of this work, and aided in writing of the manuscript. R.A.F. co-conceived and supervised the project, helped interpret the work and supervised writing of the manuscript.

## Extended Data Figures

**Extended data figure 1. Combined anti-IFNAR2+RDV therapy attenuates the hyperactive immune/inflammatory response and prevents transition to fibrosis (matched to figure 1)**.

a. Human immune cells (numbers) in BAL (14dpi) or lungs (14 and 28dpi) of SARS-Cov-2 infected MISTRG6-hACE2 mice treated with dexamethasone (Dex), Remdesivir (RDV), anti-IFNAR2 or a combined therapy of Remdesivir (RDV) and anti-IFNAR2. 14dpi, BAL: CTRL-infected n=5, Dex; RDV anti-IFNAR2 n=3; anti-IFNAR2+RDV n=4 biologically independent mice examined over 2 independent experiments. 14dpi, Lung: CTRL-infected n=7; Dex, RDV, anti-IFNAR2, anti-IFNAR2+RDV n=4 biologically independent mice examined over 3 independent experiments. 28dpi, lung: CTRL infected n=5, Dex n=4, RDV n=3, anti-IFNAR2 n=3, anti-IFNAR2+RDV n=6 biologically independent mice examined over 3 independent experiments. Means with individual datapoints plotted. Unpaired, two-tailed t-test. P<0.0001=8.19×10^−5^.

b. Representative flow cytometry plots and frequencies of alveolar macrophages (AMs) (middle: hCD206^hi^hCD86^+^hCD68^+^) or inflammatory macrophages (bottom: hCD206^lo/-^hCD86^+^hCD68^+^) in 14dpi or 28dpi lungs of treated or untreated MISTRG6-hACE2 mice. 14dpi: CTRL infected n=5, Dex n=4, RDV n=4, anti-IFNAR2 n=4, anti-IFNAR2+RDV n=4 biologically independent mice examined over at least 2 independent experiments. 28dpi: CTRL infected n=5, Dex n=4, RDV n=3, anti-IFNAR2 n=3, anti-IFNAR2+RDV n=6 biologically independent mice examined over 3 independent experiments. Means with individual datapoints. Unpaired, two-tailed t-test. P<0.0001=4.67×10^−5^.

c. Frequencies (left) and numbers (right) of lung pDCs at 14dpi. CTRL-infected n=6, Dex n=4, RDV, anti-IFNAR2 n=3, anti-IFNAR2+RDV n=6 mice examined over at least 2 experiments. Means with datapoints. Unpaired, two-tailed t-test. P<0.0001=7.29×10^−5^.

d. *IFNA* transcript levels measured by qPCR in treated or control untreated MISTRG6-hACE2 mice infected with SARS-CoV-2: Uninfected n=5; CTRL infected: 4dpi n=8, 14dpi n =9, 28dpi n=: 6; Dex 14dpi n=4, Dex 28dpi= 4; RDV 14 and 28dpi n=3, anti-IFNAR2 14 and 28dpi n=3, anti-IFNAR2+ Remdesivir 14 and 28dpi n=4 biologically independent mice examined over at least 2 independent experiments. Normalized to *HPRT1*. Violin plots with individual datapoints. Unpaired, two-tailed t test.

e. Representative histograms and frequencies of HLA-DR^+^ activated T cells in treated or control mice. 14dpi: CTRL-infected n=5, Dex, RDV, anti-IFNAR2, anti-IFNAR2+RDV n=4; 28dpi: CTRL infected, Dex, anti-IFNAR2+RDV n=4, RDV, anti-IFNAR2 n=3 biologically independent mice examined over 3 independent experiments. Means with datapoints. Unpaired, two-tailed t-test.

f. Representative images of Trichrome staining and box and whisker plot (min to max, with all datapoints) of the trichrome scoring of MISTRG6-hACE2 mice treated with a combined therapy of Remdesivir and anti-IFNAR2 or not (CTRL infected). The whiskers go down to the smallest value (minimum) and up to the largest value (maximum). The box extends from the 25th to 75th percentiles. The median is shown as a line in the center of the box. N=4 biologically independent mice examined over 2 independent experiments. Unpaired, two-tailed t-test.

Some of the data associated with dexamethasone therapy used here as a control have been reported^19^.

**Extended data figure 2. Cellular source of persistent SARS-CoV-2 viral RNA and sustained viral replication in lungs (matched to figure 2)**.

a. Quantification of genomic (gRNA) and subgenomic (sgRNA) viral RNA (E-gene) in whole homogenized lung tissue at 4, 14 and 28dpi. 4dpi: n=7, 14dpi n=5, 28dpi n=4 biologically independent mice examined over 3 independent experiments. Means with all datapoints and SD.

b. Quantification of genomic (gRNA) and subgenomic (sgRNA) viral RNA (E-gene) in whole homogenized lung tissue at 14dpi in mice treated with combined therapy of Remdesivir and anti-IFNAR2. CTRL: n=4, anti-IFNAR2+RDV: n=4 biologically independent mice examined over 2 independent experiments. N.D.= not detected.

c. Quantification of genomic (gRNA) and subgenomic (sgRNA) viral RNA (E-gene) in whole homogenized lung tissue at 28dpi in mice treated with Remdesivir, anti-IFNAR2 or combined therapy of Remdesivir and anti-IFNAR2. N=3 biologically independent mice representative of 2 independent experiments. N.D.= not detected.

d. Representative gating strategy for sorting human immune cells (human CD45+) or epithelial cells (mouse EPCAM+) from lungs of mice infected with SARS-CoV-2 and quantification of viral RNA (E and N genes) in these sorted cells. N gene: 4dpi n=3, 14dpi n=6(epithelial), n=5 (immune), 28dpi n=4 (epithelial) n=3 (immune) biologically independent mice analyzed over 3 independent experiments. E gene: 4dpi n=3, 14dpi n=7 (epithelial), n=6 (immune), 28dpi n=4 (epithelial) n=3 (immune) biologically independent mice analyzed over 3 independent experiments.

e. mNG signal in epithelial (EPCAM+) cells from lungs and BAL of mice infected with reporter SARS-CoV-2-mNG or control wild type SARS-CoV-2/WA1. mNG is expressed in infected cells following viral replication. Representative of n=4 biologically independent mice examined over 2 independent experiments.

f. Representative histograms of mNG expression in human or mouse lung macrophages isolated from BAL of infected MISTRG6-hACE2 mice at 4dpi. Representative of n=3 biologically independent mice examined over 2 independent experiments.

g. Frequencies of mNG+ cells within human lung immune cells (hCD45^+^) of SARS-CoV-2-mNG infected MISTRG6-ACE2 mice at 4dpi and 14dpi. 4dpi n=4, 14dpi n=6 biologically independent mice examined over at least 2 experiments. Unpaired, two-tailed t-test. P value=0.066.

h. Viral titers measured as PFU using Vero ACE2+ TMPRSS2+ cells that over express ACE2 from lung homogenates of MISTRG6 mice transduced with AAV-hACE2 (+AAV) or not (-AAV) and infected with SARS-CoV-2. MISTRG6-hACE2 (+AAV): 2dpi n=2, 4dpi n=5, 7dpi n=2, 14dpi n=6 MISTRG6(-AAV): 2dpi n=4, 4dpi n=10 and 7, 14 dpi n=2, biologically independent mice representative of at least 2 independent experiments. Viral titers using standard Vero E6 cells do not have any detectable titers (previously reported^19^) in MISTRG6 mice without AAV. Some of the MISTRG6-hACE2 data presented here have been previously reported as part of the characterization of the model^19^.

i. Frequencies of mNG+ cells within human macrophages (human CD68+) isolated from lungs of infected MISTRG6 mice transduced with AAV-hACE2 (AAV+) or not (AAV-). MISTRG6 mice with and without AAV-hACE2 were reconstituted with human progenitor cells from the same donor. AAV+ n=6, AAV-n=5 biologically independent mice examined over 3 independent experiments.

**Extended data figure 3. Viral RNA dependent RNA polymerase** (**RdRp) and Spike in human macrophages of human autopsy lungs with SARS-CoV-2 pneumonia (matched to figure 2)**.

Representative fluorescent microscopy images and quantification of colocalization of Spike (S), RNA dependent RNA polymerase (RdRp), human CD68 and DAPI staining in fixed human autopsy lungs with SARS-CoV-2 pneumonia or non-SARS-CoV-2 pneumonia. Quantification was performed based on representative high-power images (40x) in areas showing diffuse alveolar damage. Top panels: Representative of RdRp staining with human CD68; middle panels: Representative of Spike staining with CD68; bottom panels: RdRp and Spike staining in SARS-CoV-2-infected autopsy lungs. Yellow rectangle provides a higher magnification view of the selected area. Pseudo-colors are assigned for visualization. SARS-CoV-2 pneumonia n=4, non-SARS-CoV-2 pneumonia n=3 biologically independent specimens.

**Extended data figure 4. Human lung macrophage infection was enhanced by antibodies and reduced by CD16, ACE2, or RdRp blockade (matched to figure 2)**.

a. Viral titers measured by PFU in lung homogenates of Remdesivir (RDV) treated or control untreated MISTRG6-hACE2 mice infected with SARS-CoV-2-mNG. Infected MISTRG6-hACE2 mice were treated with Remdesivir twice daily starting at 1dpi for 3 days. CTRL infected n=6, RDV treated n=6 biologically independent mice examined over 3 independent experiments.

b. Representative histograms and mean florescent intensity (MFI) for ACE2 expression in mNG+ or mNG-epithelial cells from MISTRG6-hACE2 mice or total epithelial cells from MISTRG6 (AAV-) mice infected with SARS-CoV-2-mNG. AAV+ N=10, AAV-N=6 biologically independent mice examined over at least 3 independent experiments. Paired, two-tailed t-test.

c. Representative histograms for ACE2 expression in mNG+ or mNG-human macrophages, human B cells (CD19+) or mouse immune cells isolated from MISTRG6-hACE2 mice infected with SARS-CoV-2-mNG. Representative of N=10 for epithelial cells, N=7 for human macrophages biologically independent mice examined over at least 3 independent experiments.

d. Mean florescent intensity (MFI) of ACE2 expression in mNG+ or mNG-human macrophages or mouse epithelial cells isolated from MISTRG6-hACE2 mice infected with SARS-CoV-2-mNG. Epithelial cells n=10, human macrophages n=7 biologically independent mice examined over at least 3 independent experiments. Paired, two-tailed t-test.

e. Mean florescent intensity (MFI) of ACE2 expression in mNG+ or mNG-human macrophages isolated from MISTRG6 (AAV-) mice infected with SARS-CoV-2-mNG. Epithelial cells are virtually not infected with SARS-CoV-2-mNG in MISTRG6 mice without transduced hACE2. N=8 biologically independent mice examined over at least 3 independent experiments. Paired, two-tailed t-test.

f. Representative fluorescent microscopy images showing colocalization of human ACE2 and human CD68 cells in SARS-CoV-2 infected MISTRG6-hACE2 mice. Representative of 3 independent mice over 2 independent experiments.

g. Anti-Spike (RBD) IgG levels measured by ELISA in serum or lung homogenates of SARS-CoV-2 infected (4 and 14dpi) or uninfected MISTRG6-hACE2 mice treated therapeutically with mAbs (treated at 35hpi or 7dpi) or not. Lung homogenates: Uninfected n=5, 4dpi n=8, 14dpi n=8, 4dpi+mAB n=2, 14dpi+mAB n=2 biologically independent mice representative of at least 2 experiments. Serum: Uninfected n=3, 4dpi n=3, 14dpi n=3, 4dpi+mAB n=3, 14dpi+mAB n=2 biologically independent mice representative of at least 2 experiments. Unpaired, two-tailed t-test.

h. Frequencies of mNG signal in human immune cells in infected mice (14dpi) treated therapeutically with monoclonal antibodies (mAb) at 7dpi. MISTRG6-hACE2 mice received a mixed cocktail of monoclonal antibodies clone 135 (m135) and clone 144(m144) at 20mg/kg at 7dpi or left untreated (CTRL-infected). Monoclonal recombinant antibodies (mAbs) used in this study were cloned from the convalescent patients and had high neutralizing activity against SARS-CoV-2 *in vitro* and *in vivo* in mouse-adapted SARS-CoV-2 infection^45,64^ CTRL infected n=5, mAb treated n=4 biologically independent mice examined over 2 independent experiments. Means with datapoints and SD. Paired, two-tailed t-test.

i. Two-way plot showing anti-Spike IgG levels and corresponding mNG+ human immune cell proportions (within total human immune cells) in lungs of infected MISTRG6-hACE2 mice at 4dpi. Pearson’s correlation value =0.70. N=8 biologically independent mice examined over 4 independent experiments.

j. Frequencies of mNG+ human macrophages in human immune cells in infected mice treated with anti-CD16 antibody (Abcam-clone SP175) at 7dpi and 11 dpi and analyzed at 14dpi. MISTRG6-hACE2 mice were infected with SARS-CoV-2-mNG. n=6 biologically independent mice examined over 3 independent experiments. Unpaired, two-tailed t-test.

**Extended data figure 5. SARS-CoV-2-mNG can infect human macrophages in vitro (matched to figure 2)**.

a. Representative histograms and frequencies of mNG+ cells in bone marrow derived macrophages (BMDM) cultured (or not) with SARS-CoV-2-mNG. Cells were treated with pooled plasma from healthy controls or convalescent COVID-19 patients. mNG+ macrophages were pre-gated on live (live-dead marker negative) cells at 48hpi. Convalescent plasma samples from the top 30 neutralizers in a cohort of 148 individuals were pooled to create a mixture with an NT50 titer of 1597 against HIV-1 pseudotyped with SARS-CoV-2 S protein^45^. Healthy plasma was collected from healthy volunteers and pooled prior to COVID-19 pandemic. Concentration of plasma used 5ul plasma/ml. Uninfected n=3, infected+ healthy plasma n=7, infected+ COVID plasma n=10 independent samples cultured and analyzed over at least 3 experiments. Means with datapoints. Unpaired t-test. P<0.0001=1.57×10^−5^.

b. Quantification of genomic (gRNA) and subgenomic (sgRNA) viral RNA (E gene) in infected BMDM at 48hpi. Cells were treated with plasma from healthy controls or convalescent COVID-19 patients. Healthy plasma: n=4, COVID plasma n=6, RDV: n=6, anti-CD16+anti-ACE2 n=4 independent samples analyzed over at least 2 independent experiments. Means with datapoints. Mann-Whitney, two-tailed, t-test.

c. Representative histograms and frequencies of mNG+ cells in BMDM and lung macrophages cultured with SARS-CoV-2 in presence of plasma of convalescent COVID-19 patients. mNG+ macrophages were pre-gated on live (live-dead stain negative) cells at 48hpi. BMDM N=6, Lung macrophages N=4 independent samples analyzed over 2 independent experiments. Unpaired, two-tailed t-test.

d. Frequencies of mNG+ cells in BMDM cultured with SARS-CoV-2 or not in presence of healthy patient plasma, COVID plasma, monoclonal antibodies (clones 135 and 144) or no antibodies. COVID plasma n=5, mAb n=4, healthy plasma n=4, no Ab n=5 independent samples analyzed over 2 independent experiments. Means with datapoints and SD. The same monoclonal antibody cocktail used was used in vivo (Fig. 3). Unpaired, two-tailed t-test.

e. Representative histograms and frequencies of mNG+ cells in bone marrow derived macrophages cultured with SARS-CoV-2-mNG (or not) in presence or absence of COVID plasma. Cultures were treated with Remdesivir, anti-human CD16 antibody and/or anti-human ACE2 antibody. Healthy plasma n=5, COVID plasma n=10, RDV n=5, anti-CD16 n=6, anti-ACE2 n=4, anti-CD16+ACE2 n=4 independent samples analyzed over at least 2 independent experiments. Means with datapoints. Unpaired two-tailed t-test. P values<0.0001: anti-CD16 vs COVID plasma= 1.98×10^−5^, RDV vs. COVID plasma= 5.24×10^−6^.

f. Viral titers measured as PFUs from supernatants of human or mouse BMDM infected with SARS-CoV-2 mNG in vitro (without COVID plasma). Infectious virus from supernatants of infected macrophage cultures collected at 24hpi, 48hpi and 72 hpi was plaqued using Vero ACE2+TMPRSS2+ cells. Plaques were resolved at 48hpi. Representative images of plaques were presented as reference. Supernatant collected from Vero E6 cell cultures were provided as reference. Human: 24hpi n=9, 48 hpi n=13, 72hpi n=4. Mouse: 24hpi n=6 independent samples analyzed over at least 2 independent experiments.

g. Viral titers measured as PFUs and representative plaque from supernatants of BMDM infected with SARS-CoV-2 mNG in vitro and treated with Remdesivir (RDV) or a combination of anti-CD16 and anti-ACE2 antibodies. Cultures were not supplemented with COVID plasma. Infectious virus from supernatants of infected macrophage cultures collected at 24hpi was plaqued using Vero ACE2+TMPRSS2+ cells. Plaques were resolved at 48hpi. CTRL n=9, RDV n=4, anti-CD16 and anti-ACE2 n=4 independent samples representative of 2 independent experiments.

h. Viral titers measured as PFUs using Vero ACE2+ TMPRSS2+ cells using supernatants containing concentrations of Remdesivir (1μm) or anti-ACE2 (1μg/ml) and anti-CD16 antibodies diluted to (1:10) allow quantification of PFUs at 24hpi from macrophage cultures. Supernatants were applied on Vero ACE2+ TMPRSS2+ cells which were then infected with a matched inoculum of SARS-CoV-2 mNG (10^3 PFU quantified in Vero E6 cells) to test carry over effect in plaque quantification. Plaques were resolved at 48hpi. Untreated N=9, RDV N=6, anti-ACE2+anti-CD16 n=4 independent datapoints collected over 3 independent experiments. Means with all datapoints. Unpaired, two-tailed, t-test.

**Extended data figure 6. SARS-CoV-2 infection of human macrophages activates inflammasomes and leads to death by pyroptosis (matched to figure 3)**.

a. Representative images of single stained cells for ASC specks, NLRP3, CD14, human CD45, mouse CD45 and mouse EPCAM. Cells from SARS-CoV-2 infected humanized mice were sorted based on: human immune cells (hCD45^+^); mouse immune cells (mCD45^+^) or epithelial mouse cells (EPCAM^+^). Sorted cells were stained with single antibodies against ASC, CD14 or NLRP3. Left panel shows human immune cells and right panel shows mouse immune cell (mCD45^+^) and mouse epithelial cell (EPCAM^+^). Representative of n=5 independent mouse examined over 3 independent experiments.

b. Visualization of ASC specks as a measure of inflammasome activation in mNG+ (SARS-Cov2+) or mNG-(SARS-Cov2-) human immune cells at 4dpi. Human immune cells were sorted from SARS-CoV-2 infected humanized mice were sorted based on expression of human CD45 and mNG and lack of mouse CD45 and EPCAM expression. Representative of n=5 biologically independent mice examined over 3 independent experiments.

c. Left: Representative flow cytometry plot displaying SARS-CoV-2-mNG and Casp1-FLICA staining of CD11b^+^ human immune cells. Right: quantification of FLICA^+^ cells (%) as a measure of active caspase-1 in infected (mNG+) and uninfected (mNG-) human lung macrophages (CD11b^+^hCD45^+^) at 4dpi and 14dpi. 4dpi: n=3 biologically independent mice examined over 2 independent experiments, 14dpi n=5 biologically independent mice examined over 3 independent experiments. Lung cells were incubated with FLICA-Casp1 substrate for 30 minutes. Means with individual datapoints plotted. Paired, two-tailed t-test. P<0.0001=4.29×10^−9^.

d. Quantification of Casp1-FLICA staining as a measure of active caspase-1 in infected (mNG+) human or total mouse CD11b+ cells at 4dpi. Mouse cells: mCD45+CD11b+hCD45-. Human cells: mCD45-CD11b+hCD45+mNG+. N=6 biologically independent mice examined over 3 independent experiments. Means with individual datapoints plotted. Paired, two-tailed t-test. P<0.0001=1.79×10^−9^.

e. Human IL-18 (measured by ELISA) in lungs and serum and corresponding mNG levels (measured as percent within human immune cells by flow cytometry) in lungs of infected MISTRG6-hCE2 mice at 4dpi. Lung: Pearson’s correlation value=0.69. N=8 biologically independent mice examined over 3 independent experiments. Serum: Pearson’s correlation value=0.75. n=7 biologically independent mice examined over 3 independent experiments. Unpaired, two-tailed t-test.

f. Human IL-1RA (measured by ELISA) in lungs and serum and corresponding mNG levels (measured as percent within human immune cells by flow cytometry) in lungs of infected MISTRG6-hCE2 mice at 4dpi. Lung: Pearson’s correlation value=0.82 n=8 biologically independent mice examined over 3 independent experiments. Serum: Pearson’s correlation value=0.46 n=8 biologically independent mice examined over 3 independent experiments. Unpaired t-test, two-tailed.

g. LHD levels measured as absorbance at OD 490nm in serum of uninfected or infected MISTRG6-hACE2 mice at 4dpi and 14dpi. Fresh serum was assayed for LDH. Uninfected n=3, 4dpi n=4, 14dpi n=4 biologically independent mice examined over 2 independent experiments. Means with individual datapoints. Unpaired, two-tailed t-test.

h. Zombie Aqua incorporation in infected (mNG^+^) or uninfected (mNG^-^) CD16^+^CD11b^+^ or CD16^-^CD11b^+^ human myeloid cells. Frequencies of Zombie+ cells were measured in Annexin V-cells. N=5 biologically independent mice examined over 2 independent experiments. Means with individual datapoints. Paired, two-tailed t-test.

**Extended data figure 7. SARS-CoV-2 infection of human macrophages activates inflammasomes in vitro (matched to figure 3)**.

a. Representative histograms and quantification of Casp1-FLICA staining as a measure of active caspase-1 in bone marrow derived macrophages (BMDM) infected with SARS-CoV-2 in vitro or not for 48 hours. BMDM cultures were either supplemented with healthy or COVID plasma or monoclonal antibodies for the duration of the infection. Cultures were treated with Remdesivir, anti-ACE2 and anti-CD16 to block viral replication or viral entry. Coloring on the histograms matches the bar graph legend. Uninfected n=4; healthy plasma CTRL infected n=9, anti-ACE2 n=4, RDV n=4; mAb n= 4; COVID plasma CTRL infected n=12, anti-ACE2+anti-CD16 n=6, RDV n=5 independent datapoints collected over at least 2 independent experiments. Means with all datapoints. Unpaired, two-tailed t-test. P-values< 0.0001: COVID plasma vs. RDV: 5.85×10^−6^.

b. Representative histograms and quantification of IL-1β in supernatants of BMDM infected with (or not) SARS-CoV-2 in vitro for 48 or 72 hours. Uninfected n=7, 48hpi n=10, 72hpi n=5 independent datapoints collected over 3 independent experiments. Means with SD and individual datapoints. Unpaired, two-tailed t-test. P-values< 0.0001: uninfected vs 72hpi= 4.96×10^−7^, 48hpi vs 72hpi=1.17 ×10^−8^.

c. Human IL-18 levels at 48hpi in supernatants of BMDM infected or not with SARS-CoV-2 in vitro. BMDM cultures were supplemented with COVID plasma for the duration of the infection. Uninfected n=4, 48hpi n=7 independent datapoints collected over at least 2 independent experiments. Unpaired, two-tailed t-test. Means with individual datapoints. Unpaired, two-tailed t-test.

d. Human IL-1RA levels at 48hpi in supernatants of BMDM infected with SARS-CoV-2 in vitro or not. Uninfected n=3, infected n=7 independent datapoints collected over at least 2 independent experiments. Unpaired, two-tailed t-test. Means with individual datapoints. Unpaired, two-tailed t-test.

e. Gasdermin D (GSDMD) levels in supernatants of BMDM infected with (or not) SARS-CoV-2 *in vitro* for 48 hours. BMDM cultures were supplemented with COVID plasma for the duration of the infection. Cultures were treated with Remdesivir to block viral replication. Uninfected N=3, CTRL infected n=10, RDV n=3 independent datapoints collected over at least 2 independent experiments. Means with individual datapoints. Unpaired, two-tailed t-test.

f. LHD levels measured by absorbance at OD 490nm in supernatants of infected or uninfected. Infected BMDM were treated with caspase-1 inhibitor or not. Uninfected n=6, Infected n=11 independent datapoints collected over 3 independent experiments. Means with individual datapoints. Unpaired, two-tailed t-test. P-value= 7.38×10^−6^.

g. Zombie Aqua incorporation in uninfected or SARS-CoV-2-mNG infected BMDM. Frequencies of Zombie+ cells within Annexin V-population at 48hpi are presented. Uninfected n=4, infected n=7 independent datapoints collected over 3 independent experiments. Means with individual datapoints. Unpaired, two-tailed t-test.

**Extended data figure 8. Promoting or blocking viral entry or replication in human macrophages in vivo and in vitro impacts inflammatory profile of macrophages (matched to figure 3)**.

a. Human IL-18 levels measured in lung homogenates of infected (4dpi) MISTRG6-hACE2 mice treated (or not) with CD16 blocking antibody (Abcam). CTRL-infected N=7, ant-CD16 n=5 biologically independent mice examined over 3 independent experiments. Means with datapoints. Unpaired, two-tailed t-test.

b. Human IL-1RA levels measured in lung homogenates of infected (4dpi) MISTRG6-hACE2 mice treated (or not) with CD16 blocking antibody (Abcam). N=4 biologically independent mice examined over 2 independent experiments. Means with all datapoints. Unpaired, two-tailed t-test.

c. Human CXCL10 levels measured in serum of infected (4dpi) MISTRG6-hACE2 mice treated (or not) with CD16 blocking antibody (Abcam). CTRL infected N=7, anti-CD16 n=4 biologically independent mice examined over at least 2 experiments. Mean with individual values. Unpaired, two-tailed t-test.

d. Human IL-18 levels measured in serum of infected (14dpi) MISTRG6-hACE2 mice treated (or not) with CD16 blocking antibody (Abcam). Mice were treated with anti-CD16 blocking antibody at 7dpi and 11dpi. CTRL infected N=4, anti-CD16 n=4 biologically independent mice examined over 2 independent experiments. Means with individual datapoints. Unpaired, two-tailed t-test.

e. Human IL-18 levels measured in serum of infected (4dpi) MISTRG6 mice treated (or not) with anti-ACE2 antibody (Abcam). Mice were treated with anti-ACE2 antibody at 1,2,3 dpi. Uninfected n=4, CTRL infected n=5, anti-ACE2 n=4 biologically independent mice examined over 2 independent experiments. Means with individual datapoints. Unpaired, two-tailed t-test. P<0.0001= uninfected vs CTRL-infected=1.43×10^−5^, CTRL-infected vs anti-ACE2=6.95×10^−5^.

f. Representative flow cytometry plots of CD14 staining on total human immune cells (hCD45+) as a proxy for myeloid cells in infected MISTRG6-hACE2 mice (4dpi) treated (or not) with a depleting antibody against CD16 (ThermoFisher, clone 3G8). MISTRG6-hACE2 mice were infected with SARS-CoV-2-mNG. Representative of n=4 biologically independent mice examined over 2 independent experiments.

g. Frequencies of mNG+ cells in infected MISTRG6-hACE2 mice at 4dpi treated (or not) with a depleting antibody against CD16 (ThermoFisher, clone 3G8). N=4 biologically independent mice examined over 2 independent experiments. Means with datapoints. Unpaired, two-tailed t-test.

h. Human IL-18 levels in serum of infected or uninfected MISTRG6-hACE2 mice that were therapeutically treated with monoclonal antibodies (mAb) at 36hpi or not. Sera from infected mice were analyzed at 4dpi. N=4 biologically independent mice examined over 2 independent experiments. Means with datapoints. Unpaired, two-tailed t-test.

i. Human IL-1RA levels in serum of infected (4dpi) or uninfected MISTRG6-hACE2 mice that were therapeutically treated with mAb at 36hpi or not. Uninfected and mAB treated N=4, CTRL infected n=3 (matched to mAb treatment) biologically independent mice examined over 2 independent experiments. Means with datapoints. Paired, two-tailed t-test.

j. Human IL-18 levels measured in serum of infected (14dpi) MISTRG6-hACE2 mice treated (or not) with mAb (clone 135+ clone 144) at 7dpi and analyzed at 14dpi. N=3 biologically independent mice examined over 2 independent experiments. Mean with individual datapoints. Unpaired, two-tailed t-test.

k. Frequencies of CXCL10+ macrophages within total human macrophages (hCD45+hCD68+) in lungs of infected MISTRG6-hACE2 mice treated (or not) with mAb (clone 135, clone 144) at 7dpi and analyzed at 14dpi. Mean with individual values. Unpaired, two-tailed t-test. CTRL-infected N=5, mAb N=4 biologically independent mice examined over 2 independent experiments.

l. Box and whisker plot (min to max, with all datapoints) of the histopathological scoring of the H&E staining of infected MISTRG6-hACE2 lungs at 4dpi. Mice were either treated with monoclonal antibodies at 35hpi or anti-CD16 at 2dpi. CTRL infected N=9, mAb treated n=6, anti-CD16 treated n=3 biologically independent mice examined over at least 2 independent experiments. The whiskers go down to the smallest value (minimum) and up to the largest value (maximum). The box extends from the 25th to 75th percentiles. The median is shown as a line in the center of the box. Unpaired t-test, not significant.

m. Box and whisker plot (min to max, with all datapoints) of the histopathological scoring of the H&E staining of infected MISTRG6-hACE2 lungs at 14dpi. Mice were either treated with monoclonal antibodies at 7dpi or anti-CD16 at 7 and 11dpi. CTRL infected N=4, mAb treated n=4, anti-CD16 treated n=4 biologically independent mice examined over 2 independent experiments. The whiskers go down to the smallest value (minimum) and up to the largest value (maximum). The box extends from the 25th to 75th percentiles. The median is shown as a line in the center of the box. Unpaired, two-tailed t-test, not significant.

n. Human IL-18 levels in supernatants of SARS-CoV-2 infected BMDM treated with anti-CD16 and anti-ACE2 antibodies to block viral entry or with Remdesivir to block viral replication. CTRL infected n=8, anti-CD16+anti-ACE2 n=5, RDV n=5 independent datapoints over 3 independent experiments. Means with all datapoints. Unpaired, two-tailed t-test. P<0.0001= 5.0×10^−5^.

o. Human IL-1β levels in supernatants of SARS-CoV-2 infected BMDM treated with anti-CD16 and anti-ACE2 antibodies to block viral entry or with Remdesivir to block viral replication. N=4 independent datapoints over 2 independent experiments. Means with all datapoints. Unpaired, two-tailed, t-test.

p. Human IL-1RA in supernatants of SARS-CoV-2 infected BMDM treated with anti-CD16 and anti-ACE2 antibodies to block viral entry or with Remdesivir to block viral replication. CTRL infected n=7, anti-CD16+anti-ACE2 N=3, RDV n=3 independent datapoints over 2 independent experiments. Means with all datapoints. Unpaired, two-tailed, t-test.

q. Human CXCL10 levels in supernatants of BMDM infected *in vitro* in presence or absence of anti-CD16 and anti-ACE2 antibodies to block viral entry. CTRL infected N=12, anti-ACE2+ anti-CD16 treated n=6 independent datapoints over 2 independent experiments. Means with all datapoints. Unpaired, two-tailed, t-test.

**Extended data figure 9. Blockade of inflammasome activation leads to reduced cytokine production in vitro (matched to figure 4)**.

a. Representative flow cytometry plots of CXCL10 and TNF staining in total macrophages and histograms of CXCL10 expression in infected (mNG^+^) and uninfected (mNG^-^) macrophages from lungs of SARS-CoV-2-mNG infected MISTRG6-hACE2 mice treated with caspase-1 (Casp1) or NLRP3 inhibitors in vivo. Mice were treated on days 6,8,10,12 post-infection and analyzed at 14dpi. Representative of n=5 biologically independent mice.

b. Mean florescent intensity (MFI) of CXCL10 expression in human macrophages isolated from infected MISTRG6-hACE2 mice treated with Casp1 inhibitor or left untreated. N=3 biologically independent mice. Representative of 3 independent experiments. Means with all datapoints and SD. Unpaired, two-tailed t-test.

c. CXCL10 levels in serum of SARS-CoV-2-mNG infected MISTRG6-hACE2 mice (14dpi) treated with Casp1 or NLRP3 inhibitors. n=4 biologically independent mice examined over 2 independent experiments. Means with all datapoints and SD. Unpaired, two-tailed t-test.

d. Frequencies of mNG^+^ bone marrow-derived macrophages (BMDM) infected with SARS-CoV-2-mNG in vitro. BMDM were treated with caspase-1 (Casp1) or NLRP3 inhibitors or left untreated and analyzed at 48hpi. CTRL infected n=22, Casp1 inhibitor n=17, NLRP3 inhibitor n=6 independent datapoints collected over at least 3-experiments. Means with all datapoints and SD. Unpaired, two-tailed t-test.

e. Frequencies of FLICA^+^ BMDM infected with SARS-CoV-2 in vitro for 48 hours. BMDM were treated with Casp1 or NLRP3 inhibitors or left untreated. CTRL infected n=13, Casp1 inhibitor n=12, NLRP3 inhibitor n=6 independent datapoints collected over at least 3 independent experiments. Means with all datapoints. Unpaired, two-tailed t-test. P<0.0001=3.33×10^−5^.

f. Human IL-18 levels in supernatants of SARS-CoV-2-mNG infected BMDM treated with caspase-1 (Casp1) inhibitor or left untreated. CTRL infected n=8, Casp1 inhibitor-treated n=6 independent datapoints collected over 2 independent experiments. Means with all datapoints and SD. Unpaired, two-tailed t-test.

g. Representative histograms and quantification of IL-1β in supernatants of BMDM infected with SARS-CoV-2 in vitro. Cultures were treated with caspase-1 (Casp1) inhibitor. over at least 2 independent experiments. Uninfected n=7; 48hpi CTRL infected n=10, Casp1 inhibitor n=9, NLRP3 inhibitor n=3; 72hpi CTRL infected n=5 Casp1 inhibitor n=3, NLRP3 inhibitor n=3 independent datapoints collected over at least 2 experiments. Means with all datapoints and SD. Unpaired, two-tailed t-test. P<0.0001=4.96×10^−7^.

h. Human Gasdermin D (GSDMD) levels at 48hpi in supernatants of SARS-CoV-2-mNG infected BMDM treated with Casp1 inhibitor or left untreated. CTRL infected n=10, Casp1 inhibitor-treated n=6 independent datapoints collected over at least 3 independent experiments. Means with all datapoints. Unpaired, two-tailed t-test.

i. LDH levels measured as absorbance at OD 490nm in supernatants of uninfected or SARS-CoV-2-mNG infected BMDM treated with Casp1 or NLRP3 inhibitor or left untreated in vitro. Uninfected n=6, CTRL infected (48hpi) n=11, Casp1 inhibitor-treated(48hpi) n=9, NLRP3 inhibitor-treated (48hpi) n=5 independent datapoints collected over 2 independent experiments. Means with all datapoints and SD. Unpaired, two-tailed t-test. P<0.0001=7.38×10^−6^.

j. Zombie Aqua incorporation in SARS-CoV-2-mNG infected BMDM treated with Casp1 or NLRP3 inhibitor or left untreated (CTRL infected). Frequencies of Zombie+ cells within Annexin V-population at 48hpi are reported. Uninfected n=4, CTRL infected n=7, Casp1 inhibitor n=4, NLRP3 inhibitor n=3 over 2 experiments. Means with all datapoints. Unpaired, two-tailed t-test.

k. Human CXCL10 levels in supernatants of infected human BMDM treated with Casp1 inhibitor or NLRP3 inhibitor or left untreated. Supernatants were collected at 48hpi. Uninfected n=5, CTRL infected n=12, Casp1 inhibitor n=5, NLRP3 n=4 independent datapoints over at least 2 independent experiments. Means with all datapoints and SD. Unpaired, two-tailed t-test.

l. Human IL-1RA levels in supernatants of SARS-CoV-2-mNG infected human BMDM treated with Casp1 inhibitor or not. BMDM were treated with Casp1 inhibitor. Supernatants were collected at 48hpi. CTRL infected n=7, Casp1 inhibitor-treated n=6, NLRP3 inhibitor-treated n=4 independent datapoints collected over at least 2 independent experiments. Means with all datapoints. Unpaired, two-tailed t-test.

m. Viral titers measured as PFU in lung homogenates of MISTRG6-hACE2 mice infected with SARS-CoV-2 and treated with caspase-1 inhibitor in vivo. Infected MISTRG6-hACE2 mice were treated with Caspase 1 inhibitor on days 6,8,10,12 post-infection and analyzed at 14dpi. Lung homogenates were plaqued using Vero ACE2+TMPRSS2+ cells. of CTRL infected: n=7, Casp1 inhibitor-treated: n=6 biologically independent mice examined over 3 independent experiments. Box and whisker plot (min to max, with all datapoints) The whiskers go down to the smallest value (minimum) and up to the largest value (maximum). The box extends from the 25th to 75th percentiles. The median is shown as a line in the center of the box. Ratio paired, two-tailed t-test.

n. Representative images of plaque assays used to quantify infectious virus in supernatants of BMDM infected with SARS-CoV-2-mNG and treated with caspase-1 or NLRP3 inhibitors. Supernatants of infected macrophage cultures was collected at 48hpi and was plaqued using Vero ACE2^+^TMPRSS2^+^ cells. Plaques were resolved at 48hpi. Representative of CTRL infected: n=13, Casp1 inhibitor-treated: n=8, NLRP3 inhibitor-treated: n=5 independent datapoints collected over 3 independent experiments.

