## Supplemental Data for "Inflammasome activation in infected macrophages drives COVID-19 pathology"

**Author information:**

1. Department of Immunobiology, Yale University School of Medicine, New Haven, CT, USA.
2. Department of Pathology, Yale University School of Medicine, New Haven, CT, USA.
3. Program in Cellular and Molecular Medicine, Boston Children's Hospital, Boston, MA, USA.
4. Department of Pediatrics, Harvard Medical School, Boston, MA, USA.
5. Instituto René Rachou, Fundação Oswaldo Cruz, Belo Horizonte, Minas Gerais, Brazil.
6. Section of Hematology, Yale Cancer Center and Department of Internal Medicine, Yale University School of Medicine, New Haven, CT.
7. Laboratory of Molecular Immunology, The Rockefeller University, New York, NY, USA.
8. Department of Laboratory Medicine, Yale University School of Medicine, New Haven, CT, USA.
9. Howard Hughes Medical Institute, Yale University School of Medicine, New Haven, CT, USA.
10. Department of Surgery, Yale University School of Medicine

### Supplementary Discussion

##### Infection of human macrophages and inflammasome activation

The MISTRG6 model of COVID-19 faithfully reflects many of the chronic immunoinflammatory features of the human disease and provides an opportunity to dissect the mechanisms of late immunopathogenesis in this disease[19]. As in severe human disease, COVID-19 in MISTRG6-hACE2 mice presents with persistent viral RNA, chronic IFN response accompanied with a chronic inflammatory state in macrophages that is initiated by infection of human macrophages and maintained by subsequent inflammasome activation[19]. These events may eventually contribute to the development of persistent pulmonary immunopathology and fibrosis, which is supported by histopathological and transcriptional analysis of lungs late in infection. Overall, our mechanistic study of this model defines a cascade of events, which, initiates with lung epithelial infection and is followed with infection of tissue-resident macrophages in an ACE2 and CD16 mediated manner. SARS-CoV-2 replicates in these macrophages generating replicative intermediates which include dsRNA, subgenomic viral RNA, viral RNA polymerase (RdRp), and expression of a virally encoded fluorescent reporter gene (mNG), all of which is inhibited by Remdesivir, an inhibitor of viral replication. SARS-CoV-2 replication and replicative intermediates activate an inflammatory program which involves activation of inflammasomes, production, and release of inflammatory cytokines and chemokines, and finally pyroptosis. We established inflammasome activation by visualizing ASC speck formation, which colocalized with active caspase-1 and NLRP3; this led to maturation of inflammasome mediated cytokines IL-1β and IL-18, and results in pyroptosis assayed by gasdermin D (GSDMD) and LDH release. Inflammasome activation and downstream effectors in these infected macrophages are caspase-1 and NLRP3 dependent, as inhibitors of both caspase-1 and at NLRP3 block all downstream aspects of inflammasome activation and the inflammatory cascade both *in vivo* and *in vitro*. More importantly, targeting inflammasome mediated hyperinflammation prevented immunopathology associated with chronic SARS-CoV-2 infection *in vivo*.

#### Mechanisms of viral uptake

Consistent with the enhancing role for antiviral antibodies in macrophage infection, COVID-19 severity in patients was correlated with early, high levels of afucosylated IgG which enhanced the inflammatory response by monocytes and macrophages through Fc-mediated interactions with CD32 and CD16[47-49]. We observe a similar role for CD16 and antibodies in humanized mice infected with SARS-CoV-2-mNG. The frequency of infected macrophages which express high levels of CD16 correlated with the levels of anti-Spike antibodies in the lung tissue, particularly at 4dpi. mNG positivity in these cells also correlated with a strong proinflammatory cytokine signature as measured by elevated levels of IL-18, IL-1RA, and CXCL10, all of which contribute to severe disease in humans. CD16 blockade *in vivo* and *in vitro* prevented viral uptake and blocked this subsequent inflammatory response as measured by reduced levels of CXCL10, IL-18, and IL-1RA.

The ACE2 receptor that is utilized by SARS-CoV-2 to infect lung epithelium is also expressed preferentially by infected human macrophages *in vivo*[43]. Notably, $CD14^{hi}CD16^{hi}$ cells and alveolar macrophages which had measurable viral RNA cells in patient samples did not appear to co-express the traditional viral entry factors, ACE2 and TMPRSS2 as measured by the relatively insensitive method of single cell RNA sequencing (scRNAseq)[7]. This may however be a technical limitation as we similarly could not detect measurable ACE2 transcript in alveolar macrophages or CXCL10+ macrophages (which is a proxy for mNG+ infected macrophages) by scRNAseq at any time point during infection. Yet, ACE2 protein clearly colocalized with CD68, a marker of human macrophages and correlated with viral replication quantified by mNG in these cells. More importantly blocking ACE2 prevented viral uptake by macrophages. Interestingly in our system, *ACE2* expression was inducible (Extended data figure 4a), and its levels, correlated well with normalized viral RNA levels measured in the same samples (Extended data figure 4b). To determine factors that could regulate ACE2 expression, we identified genes that correlate with *ACE2*. Interestingly, the top 100 genes that correlate with *ACE2* expression (r>0.6) were enriched for interferon responsive genes (Extended data figure 4c), further highlighting the importance of the interferon pathway in COVID19 patients (Extended data figure 4d).

Infection of macrophages in our system is therefore dependent on both ACE2 receptor as well as antibody-mediated uptake by CD16. Given the prevalence of antibodies increases as the disease progresses, it is likely that the latter mechanism plays a more important role later in infection. However, there may be other mechanisms that enhance SARS-CoV-2 infection or the downstream inflammatory response in human macrophages that are not explored in this study. SARS-CoV-2-mNG lacks-Orf7a which can enhance proinflammatory cytokine production in monocytes via its interaction with CD14[41,65]. It is not clear whether CD14 expression in macrophages could mediate viral entry or enhance inflammatory cytokine production[65]. It has been noted that low molecular weight immune complexes formed prior to the specific humoral response, combined with the afucosylated state of IgGs, can further enhance the CD16-mediated activation of monocytes and macrophages, mimicking a state similar to systemic lupus erythematosus (SLE) disease. Given that we observe SLE-like features in our lung transcriptome[19,66], it is possible that blocking CD16-mediated macrophage infection and activation may impact this SLE-like state observed late in our disease model.

**Infected human macrophages initiate and maintain an inflammatory cascade that impacts disease outcome**

In our mouse model, monocytes, and macrophages in SARS-CoV-2 infected MISTRG6-hACE2 are central to disease pathology and are the main source of inflammatory cytokines IL-1β, IL-1RA, IL-18, TNF-α, IL6 and inflammatory chemokines like CXCL10. It is likely that this is also true in human disease[7,15,20,67]. Of these cytokines and chemokines elevated in COVID-19 patients, IL-1β, IL-1RA, IL-18 and CXCL10 also correlated with disease severity[2-7,14,15,20]. Infected macrophages in MISTRG6-hACE2 mice (and *in vitro* infected BMDM) were the main producers of these cytokines correlating with disease severity and had a unique transcriptional signature revealed by association with *CXCL10* correlating transcripts in our transcriptional datasets. In humans, SARS-CoV-2 viral RNA was detected in mononuclear phagocytes characterized by scRNA-seq analysis of autopsied lungs of COVID-19 patients[7,20] although whether this results from viral replication in these cells or phagocytosis could not be distinguished. In line with our findings, $CD14^{hi}CD16^{hi}$ cells and alveolar macrophages in autopsied lungs of COVID-19 patients were particularly enriched with viral RNA[7,20]. We also found clear evidence for the presence of viral components Including viral RdRp in macrophages and epithelial cells of infected human lungs. Several recent studies of human macrophages and other myeloid cells also suggest that SARS-CoV can infect these cells[68-70]. However, in our humanized mouse system it is clear that the majority of RNA found associated with host cells may be the result of phagocytic or other non-replicative uptake mechanisms. It was only by using SARS-CoV-2-mNG virus (see Fig. 2, 3) that we were able to distinguish these two processes at which point we could clearly distinguish infection from mere uptake of viral debris, which in fact is prevalent. Some SARS-CoV-2+ myeloid cells in humans also had distinct transcriptomes which were largely recapitulated in what we construe as *CXCL10* associated genes (*CXCL11, CCL18, CCL8, ISG15, CD83;* Fig. 3) from MISTRG6-hACE2, with the exception of TNF which was co-expressed by these same CXCL10+ cells. This in fact is complementary to our findings where the CXCL10-associated gene signature and its inverse relationship with TNF weakens late in infection (28dpi, Fig. S3I), a time point that corresponds to autopsied lungs of severe COVID-19, suggesting a convergent inflammatory state in macrophages as inflammation progresses. Blocking inflammasome and pyroptosis by inhibition of the inflammatory cascade by caspase-1 attenuated this convergent inflammatory state and lung pathology. The effects of caspase-1 inhibition extended beyond infected macrophages to infiltrating macrophages that are not infected with virus and resulted in reduced levels of TNF. Nonetheless, this inhibition yielded substantially increased virus production. It should be noted that it is not clear to what degree these macrophages contribute, if at all, to high titers of virus production compared with the permissive epithelial cells- although fluorescent levels of mNG virus RNA was similar in the two cell types.

Although viral uptake and the subsequent antiviral immune response, such as CXCL10, IL-18, IL-1RA production, is enhanced in presence of monoclonal antibodies, the outcome of this enhancement does not appear to be pathological when given early or late. This is in line with extensive clinical findings that show patients given convalescent plasma or monoclonal antibodies responded well to therapy and did not present with disease enhancement[71]. Several lines of evidence also suggest that FcRs are essential for antibody mediated protection and therapy[72,73]. A possible explanation for this conflicting role of antiviral antibodies is potentially explainable by the fact that the antibodies enhance infection of macrophages and thus inflammation but at the same time they neutralize virus and thereby attenuate disease leading to a net null effect consistent with the enhancing role of antiviral Abs on macrophage infection.

**Viral sensing by NLRP3 in infected human macrophages**

Viral RNA and particles can be detected by a variety of innate immune sensors. Among these, myeloid cell expressed inflammasomes including NLRP3, and NLRP1 can be activated by RNA viruses[55,56,74]. Human lung monocytes and macrophages in infected MISTRG6-hACE2 mice express low levels of inflammasome sensors, NLRP3, and NLRP1. Of these NLRP3 is both upregulated and activated by replicating SARS-CoV-2 in these macrophages. NLRP3 can be activated by a diverse, promiscuous set of stimuli but initially requires a priming event, which results in the transcriptional induction of NLRP3 and triggers post-translation modifications. In line with this priming step, NLRP3 transcript expression in lung tissues of infected MISTRG6-hACE2 mice was upregulated in response to infection, but interestingly was inhibited by combined therapy of anti-IFNAR and Remdesivir or by dexamethasone, all of which reduce both levels of replicating virus and inflammatory ligand. Activation of NLRP3 by some RNA viruses relies on viral replication and direct sensing of viral RNA or via other viral RNA sensors as MDA5 and RIG-I. Viral replication in the context of an early inefficient IFN response (which SARS-CoV-2 is thought to accomplish[51,53,75]), is likely a stimulus for NLRP3 activation in human macrophages. Loss of IL-18 and IL-1β production upon inhibition of viral replication in our studies strongly suggest viral replication is involved. Recent reports have also identified a possible role for NLRP3 driven inflammasome activation in infected monocytes and macrophages in post-mortem tissue samples and peripheral blood mononuclear cells (PBMC) of COVID-19 patients. Although there have been many candidates for NLRP3 activation ligands (lytic cell death upon infection, N protein[59], Orf3a[60]), the exact mechanism of NLRP3 activation is still poorly understood, especially given the diverse set of stimuli that can activate NLRP3. Nonetheless activation of other NLRs may contribute to the process, as inhibition of caspase-1 gave in general stronger inhibition of responses than inhibition of NLRP3.

##### Infiltrating macrophages and the essential role of viral RNA-dependent type I IFN response in disease pathology

Infection causes human macrophages to preferentially produce CXCL10 which likely attract blood monocytes to the lung where they differentiate to inflammatory macrophages. These monocytes and monocyte-derived macrophages (MDM) eventually outnumber tissue-resident macrophages; they express higher levels of TLRs and may play a central role in viral RNA detection, possibly also released by pyroptosis of infected macrophages, and the ensuing inflammatory and IFN response. IFN production is critical for the antiviral response of the early phase of disease, as also evidenced in our model by drastically higher viral loads and precipitous decline in health when the antiviral response is disabled too early by dexamethasone treatment at the peak of infection[19]. However, this same response when persistent can be pathogenic. We found that targeting either chronic viral replication or the late IFN response therapeutically *in vivo* attenuates many aspects of the overactive immune-inflammatory response, especially the inflammatory macrophage response.

##### Implications of our findings

Inhibition of viral replication, viral uptake and inflammasome activation in infected macrophages reduced lung hyperinflammation with high levels of inflammasome-induced cytokine IL-18, IL-1RA, and CXCL10 in infected MISTRG6-hACE2 mice. Patients with severe COVID-19 also have higher levels of IL-18, IL-1β, IL1-RA and CXCL10 [2-6,14,34,76]. Inhibition of both caspase-1 and NLRP3 resolved lung immunopathology associated with chronic disease in MISTRG6-hACE2 mice. Given that multiple reports in patient samples also identify a role for inflammasome driven hyperinflammation in pathophysiology of COVID-19, targeting inflammasome sensors or downstream effector molecules in patients may provide alternative therapeutic options for resolving chronicity in COVID-19. However, the increased virus production seen upon inflammasome blockade could pose a significant risk to the benefit of wholesale inhibition of the pathway. The combination of Remdesivir and anti-IFNAR2 antibodies could be an effective therapy for chronic COVID-19 which spares the antiviral T cell response unlike dexamethasone. More generally, the findings from our study and its implications provide alternative therapeutic avenues to be explored in the clinic and may guide novel therapeutic developments and prompt clinical trials to investigate combinatorial therapies that target viral RNA, inflammasome activation or its products and sustained IFN response.

#### Supplementary Materials and Methods

##### Mice

MISTRG6 was generated by the R. Flavell laboratory by combining mice generated by this lab, the laboratory of Markus Manz and Regeneron Pharmaceuticals based on the *Rag2-/- IL2rg-/-*129xBalb/c background supplemented with genes for human M-CSF, IL-3, SIRPα, thrombopoietin, GM-CSF and IL-6 knocked into their respective mouse loci[77,78]. MISTRG6 mice are deposited in Jackson Laboratories and made available to academic, non-profit, and governmental institutions under a Yale-

Regeneron material transfer agreement (already approved and agreed to by all parties). Instructions on obtaining the material transfer agreement for this mouse strain will be available along with strain information and upon request. CD1 strain of mice acquired from Charles River Laboratories were used for cross-fostering of MISTRG6 pups upon birth to stabilize healthy microbiota. All mice were maintained under specific pathogen free conditions in our animal facilities (either Biosafety Level 1, 2 or 3) under our Animal Studies Committee-approved protocol. Unconstituted MISTRG6 mice were maintained with cycling treatment with enrofloxacin in the drinking water (Baytril, 0.27 mg/ml). All animal experimentations were performed in compliance with Yale Institutional Animal Care and Use Committee protocols. For SARS-CoV-2–infected mice, all procedures were performed in a Biosafety Level 3 (BSL-3) facility with approval from the Yale Institutional Animal Care and Use Committee and Yale Environmental Health and Safety.

**Transplantation of human CD34+ hematopoietic progenitor cells into mice**

Fetal liver samples were cut in small fragments, treated for 45 min at 37 °C with collagenase D (Roche, 200 μg/ml), and prepared into a cell suspension. Human CD34+ cells were purified by performing density gradient centrifugation (Lymphocyte Separation Medium, MP Biomedicals), followed by positive immunomagnetic selection with EasySep™ Human CD34 Positive Selection Kit (StemCell). For intra-hepatic engraftment, newborn 1–3-day-old pups were injected with 20,000 fetal liver CD34+ cells in 20 μl of PBS into the liver with a 22-gauge needle (Hamilton Company). All use of human materials was approved by the Yale University Human Investigation Committee.

**AAV-hACE2 administration**

AAV9 encoding hACE2[19,79] was purchased from Vector Biolabs (AAV9-CMV-hACE2). Animals were anaesthetized using isoflurane. The rostral neck was shaved and disinfected. A 5-mm incision was made, and the trachea was visualized. Using a 32-G insulin syringe, a 50-μl injection dose of $10^{11}$ genomic copies per milliliter of AAV-CMV-hACE2 was injected into the trachea. The incision was closed with 4–0 Vicryl suture and/or 3M Vetbond tissue adhesive. Following administration of analgesic animals were placed in a heated cage until full recovery. Mice were then moved to BSL-3 facilities for acclimation.

***In vivo* SARS-CoV-2 infection**

SARS-CoV-2 isolate USA-WA1/2020 was obtained from BEI reagent repository. SARS-CoV-2 mNG was obtained from Dr. P.Y. Shi (UTMB)[41]. All infection experiments were performed in a BSL-3 facility, licensed by the State of Connecticut and Yale University. Mice were anesthetized using 20% vol/vol isoflurane diluted in propylene glycol. Using a pipette, 50 µl of SARS-CoV-2-WA1 or SARS-CoV-2-mNG (1-3x$10^6$ PFU) was delivered intranasally.

**Therapeutics**

SARS-CoV-2 infected MISTRG6-hACE2 were treated intraperitoneally daily with dexamethasone at 10mg/kg for 3 days starting at 7dpi. Mice were treated subcutaneously with Remdesivir at 25mg/kg dosing as has been previously described[22] for 3 consecutive days starting at 7dpi (Fig. 1) or 1dpi (Fig. 2- for human macrophage infection studies mice were treated twice, daily). Mice were treated with anti-IFNAR2 antibody at 1.5mg/kg dosing on days 7 and 11 post infection.

Infected MISTRG6-hACE2 mice were treated with two different clones of anti-human CD16 antibodies. For CD16 blockade experiments, mice were treated with anti-CD16 (Abcam, clone SP175) antibody early and late. For early CD16 blockade studies mice were treated with anti-CD16 antibody at 2dpi with a single dose (20µg per mouse) and euthanized at 4dpi. For late CD16 blockade studies mice were treated with anti-CD16 antibody at 7dpi and 11dpi and euthanized at 14dpi. For depletion experiments mice were treated with anti-CD16 (ThermoFisher, clone 3G8) antibody with a daily dose of 20µg for 3 days starting 1dpi. Rabbit IgG, monoclonal [EPR25A] Isotype Control (ab172730) and Mouse IgG1 kappa Isotype Control (P3.6.2.8.1) were used.

Infected MISTRG6 (without AAV-hACE2) mice were treated with monoclonal antibody against human ACE2 (clone MM0073-11A31, Abcam-ab89111) for 3 days i.p. with a daily dose of 20µg starting at 1dpi. In these mice only, epithelial cells were not infected or infected poorly with SARS-CoV-2 with undetectable titers using standard plaque assays[19] presumably due to differences between mouse and human ACE2 that limit viral entry and replication[42]. Mouse IgG2 Isotype was used as control.

Infected MISTRG6-hACE2 mice received a mixed cocktail of monoclonal antibodies clone 135 (m135) and clone 144 (m144) at 20mg/kg at 35hpi or 7dpi. Monoclonal recombinant antibodies (mAbs) used in this study were cloned from the convalescent patients (whose plasma was used for in vitro studies infecting BMDM) and had high neutralizing activity against SARS-CoV-2 *in vitro* and *in vivo* in mouse adapted SARS-CoV-2 infection and ancestral stain of SARS-CoV-2/WA1[45,64,66].

For NLRP3 inhibitor experiments, Infected MISTRG6-hACE2 mice were treated with MCC950 (R&D Systems) at a dose of 8 mg/kg intraperitoneally on days 6, 8, 10,12- post infection and euthanized on day 14[80-82]. For caspase-1 inhibitor experiments, infected MISTRG6-hACE2 mice treated with VX-765

(Invivogen) at a dose of 8 mg/kg on days 6, 8, 10,12 post infection and euthanized on day 14. Control infected mice were treated with PBS[82].

**Viral titers**

Mice were euthanized in 100% isoflurane. Approximately half of the right lung lobe was placed in a bead homogenizer tube with 1 ml of DMEM+2% FBS. After homogenization, 300 µl of this mixture was placed in 1mL Trizol (Invitrogen) for RNA extraction and analysis. Remaining volume of lung homogenates was cleared of debris by centrifugation (3,900 g for 10 min). Infectious titers of SARS-CoV-2 were determined by plaque assay in Vero E6 (standard) or Vero ACE2+TMPRSS2+ (sensitive) cells in DMEM 4% FBS, and 0.6% Avicel RC-581[83]. Plaques were resolved at 48 h after infection by fixing in 10% formaldehyde for 1 hour followed by staining for 1 hour in 0.5% crystal violet in 20% ethanol. Plates were rinsed in water to visualize plaques. Multiple dilutions of lung homogenates were used to quantify Infectious titers (minimum number of plaques that can be quantified= 10 per ml of lung homogenate or ml of supernatant). Viral titers from supernatants of bone-marrow derived macrophage cultures were determined by plaque assay in Vero ACE2+TMPRSS2+ (sensitive) cells following the same protocols described for lung homogenates. VERO C1008 (Vero 76, clone E6, Vero E6) were obtained from ATCC. Vero ACE2+ TMPRSS2+ cells were obtained from B. Graham (NIAID). None of the cell lines were authenticated or tested for mycoplasma contamination.

**Viral RNA analysis**

RNA was extracted with the RNeasy mini kit (Qiagen) per the manufacturer's protocol. SARS-CoV-2 RNA levels were quantified using the Luna Universal Probe Onestep RT-qPCR kit (New England Biolabs) and US CDC real-time RT-PCR primer/probe sets for 2019-nCoV_N1. For each sample, 1 µg of RNA was used. Subgenomic viral RNA was quantified using primer and probe sets targeting E gene as has been previously described[40,63]. The primer-probe sequences were as follows: E_Sarbeco_F primer: ACAGGTACGTTAATAGTTAATAGCGT (400 nM per reaction). E_Sarbeco probe _P1: FAM-ACACTAGCCATCCTTACTGCGCTTCG-BBQ (200nM per reaction); E_Sarbeco_R primer ATATTGCAGCAGTACGCACACA (400 nM per reaction); E leader specific primer sgLead-F: CGATCTCTTGTAGATCTGTTCTC (400 nM per reaction).

**Histology and immunofluorescence**

Yale pathology kindly provided assistance with embedding, sectioning of lung tissue. A pulmonary pathologist reviewed the slides blinded and identified immune cell infiltration and other related pathologies. Paraffin embedded lung tissue (fixed in 4% paraformaldehyde for no more than 24 hours) sections were deparaffinized in xylene and rehydrated. After antigen retrieval with 10 mM Sodium Citrate pH 6 and permeabilization with 0.1% Triton-X for 10 min the slides were blocked with 5% BSA in PBS with 0.05% Tween 20 for an hour. Then the samples were stained with primary antibodies against SARS-CoV-2-dsRNA; SARS-CoV2-RNA-dependent RNA Polymerase, SARS-CoV-2-Spike, human CD68, human ACE2 their isotype controls diluted in 1%BSA overnight at 2-8 °C. The next day, the samples were washed and incubated with fluorescent secondary antibodies. After washes, samples were treated with TrueBlack lipofuscin autofluorescence quencher for 30 seconds and mounted on DAPI mounting media (Sigma). Images were acquired using Keyence BZ-X800 Fluorescence Microscope or Nikon ECLIPSE Ti Series Confocal Microscope. Pseudo-colors were assigned for visualization.

**Isolation of cells and flow cytometry**

All mice were analyzed at approximately 9-14 weeks of age. Single cell suspensions were prepared from blood, spleen, bronchioalveolar lavage (BAL) and lung. Mice were euthanized with 100% isoflurane. BAL was performed using standard methods with a 22G Catheter (BD). Blood was collected either retro-orbitally or via cardiac puncture following euthanasia. BAL was performed using standard methods with a 22G Catheter (BD)[84]. Lungs were harvested, minced, and incubated in a digestion cocktail containing 1 mg/ml collagenase D (Sigma) and 30 µg/ml DNase I (Sigma-Aldrich) in RPMI at 37°C for 20 min with gentle shaking. Tissue was then filtered through a 70 or 100-µm filter. Cells were treated with ammonium- chloride-potassium buffer and resuspended in PBS with 1% FBS. Mononuclear cells were incubated at 4C with human (BD) and mouse (BioxCell, BE0307) Fc block for 10 min. After washing, primary antibody staining was performed at 4C for 20 min. After washing with PBS, cells were fixed using 4% paraformaldehyde. For intracellular staining, cells were washed with BD permeabilization buffer and stained in the same buffer for 45 min at room temperature. Samples were analyzed on an LSRII flow cytometer (BD Biosciences). Data were analyzed using FlowJo software.

For cell sorting experiments, single cell suspensions from digested lungs were stained with antibodies against human CD45, mouse CD45, mouse EPCAM and sorted using BD FACS Aria II which is contained in a Baker BioProtect IV Biological Safety Cabinet. Cell viability was assessed with DAPI when applicable.

For imaging flow cytometry, cells from SARS-CoV-2 infected humanized mice were sorted based on: human immune cells (hCD45+); mouse immune cells (mCD45+) or epithelial mouse cells (EPCAM+). A- mNG+ epithelial cells (SARS-CoV-2-mNG+ mCD45(PE)- EPCAM(APC)+ hCD45(PB)-; B-total mouse immune cells (mCD45(PE)+ EPCAM(APC)- hCD45(PB-); C- mNG+ human immune cells (SARS-CoV-2-mNG+ mCD45(PE)- EPCAM(APC)- hCD45(PB)+); D- mNG- human immune cells (SARS-CoV-2-mNG- mCD45(PE)-EPCAM(APC)- hCD45(PB)+). These sorted cells (epithelial or immune cells) were fixed in 4% paraformaldehyde (PFA) for at least 30 minutes. Fixed sorted cells (epithelial or immune cells) were permeabilized, stained with unconjugated primary antibodies for ASC (1:200, rabbit), NLRP3 (1:200, goat), then stained with secondary antibodies (donkey anti-rabbit or goat conjugated with AlexaFluor 546 or 647, at 1:1000). Cells data were acquired using an ImageStream X MKII (Amnis) with 63X magnification and analyzed using Ideas software (Amnis). ASC, NLRP3 specks were gated and quantified based on fluorophore intensity/max pixels. For FLICA-Caspase1 colocalization, macrophages were pretreated with FLICA prior to sorting.

***In vitro* infection with SARS-CoV-2**

Using aseptic techniques under sterile conditions, bone marrow cells were isolated from femurs of reconstituted MISTRG6 mice. For differentiation into macrophages, bone marrow cells were incubated in media supplemented with 10% FBS, 1% penicillin/streptomycin and recombinant human M-CSF (50ng/ml), GM-CSF (50ng/ml) and IL-4 (20ng/ml) at $1x10^6$ per ml concentration for 6 days in 5% CO2 incubator at 37°C. Media supplemented with 10% FBS was replenished with new media every 3–4 days. Prior to infection, cells were monitored for granularity, elongated morphology, and stronger adherence to the plate. Human macrophages were then cultured with SARS-CoV-2-mNG in presence or absence of COVID patient plasma, healthy plasma, monoclonal antibodies (mix of clones 135 and 144, described as therapeutics), Remdesivir, anti-CD16 antibody, anti-ACE2 antibody, control isotype antibody, caspase-1 inhibitor (VX-765[85]) or NLRP3 inhibitor (MCC950).

*Ex vivo* lung macrophage cultures: To enrich for human macrophages and monocytes, lung cells from uninfected MISTRG6 mice were sorted based on CD11B and human CD45 expression. These cells were then incubated with GM-CSF and IL-4 for 48 hours to mature macrophages. Non-adherent cells were aspirated prior to culturing with SARS-CoV-2.

Bone-marrow derived macrophages (BMDM) *in vitro* or lung macrophages *ex vivo* were cultured with a viral inoculum at 10^4 PFU of SARS-CoV-2-mNG (~ MOI=0.1). These macrophage cultures were then incubated at 37 C, 5% CO2 for 24, 48 and 72 hours at which time cells were harvested. Cells were dissociated from culture plate with 10 mM EDTA or Accutase (ThermoFisher) cell dissociation reagent (10-20 minutes). For studies pertaining to the mechanism of viral entry, viral replication and inflammasome activation, infected macrophages were treated with Remdesivir (10uM), anti-CD16 (Abcam clone, 10μg/ml) and anti-ACE2 (10 μg/ml), caspase-1 inhibitor (VX765, 20μM) and NLRP3 inhibitor (MCC950, 20μg/ml) in culture. Cells were stained when applicable and fixed for 30 min with 4% PFA. Convalescent plasma samples from the top 30 neutralizers in a cohort of 148 individuals were pooled to create a mixture with an NT50 titer of 1597 against HIV-1 pseudotyped with SARS-CoV-2 S protein[45]. We used this pooled serum at a concentration of 5μl-plasma/ml for *in vitro* experiments and refer to it as COVID patient plasma. Healthy plasma was collected from healthy volunteers and pooled prior to COVID-19 pandemic and used at a concentration of 5μl-plasma/ml. Monoclonal antibodies (a mix of clones 135 and 144) were used at 4 μg per ml concentration.

**Zombie Aqua and Annexin V staining**

Single cell suspension from in vitro cultures or enzymatically dissociated lungs were washed and stained for viability with Zombie Aqua ((Biolegend- 423101) in PBS (1:400) for 15 min at 4C. Without washing the cells, cell surface antibody cocktail was added, and cells were incubated for another 15 minutes. Cells were then washed with PBS twice and resuspended in Annexin V binding buffer. Cells were stained with Annexin V PE (1:400) in binding buffer for 15 min at 4C. Cells were then washed with Annexin V buffer and fixed in 4% PFA.

**FLICA assay**

Single cell suspension from in vitro cultures or enzymatically dissociated lungs were resuspended in RPMI 10% FBS with FLICA substrate (BioRad-FLICA 660 caspase-1 kit- ICT9122) and cultured for 1h (for microscopy) or 30 min (for flow cytometry) at $37^0$C. Cells were then washed twice with PBS and stained with Zombie Aqua and Annexin V as described. Cells were then fixed with 1x Fixative (provided in BioRad-FLICA caspase-1 kit) for at least 1 hour not exceeding 16 hours. Cells were kept at $4^o$C until further staining and analysis. FLICA 660 caspase-1 kit uses a target sequence (YVAD) sandwiched between a far-red fluorescent 660 dye (excitation max 660nm, emission max 685nm).

**LDH measurement**

LDH levels were measured from freshly collected supernatant of infected cells (BMDM) or freshly collected serum using CyQUANT LDH Cytotoxicity Assay (ThermoFisher- C20300) following manufacturer’s instructions under BSL3 conditions.

#### Human Samples

For this study we have acquired six control uninfected, and two SARS-CoV-2 infected deidentified lung (4 different cuts) specimens as paraffin embedded tissues from autopsies of individuals admitted to Yale New Haven Hospital. Lungs were fixed in 10% Formalin (Table S5).

##### Supplementary Table 5. Patient specimens used for immunofluorescent (IF) staining.

| Case | Age | Gender | Medication | Death (dps) | Co-morbidities | Cause of death | Histopathological findings |
|---|---|---|---|---|---|---|---|
| COVID sample#1 | 54 | male | Dexamethas one, Remdesivir | 20 | Obese | Acute fibrinous organizing pneumonia | Diffuse alveolar damage, overlapping features of exudative and proliferative phase |
| COVID sample#2 | 37 | male | | 35 | alcoholic liver disease and micro nodular cirrhosis. | Complications of COVID-19 | Diffuse alveolar damage, overlapping features of exudative and proliferative phase |
| COVID sample#3 | 77 | female | Dexamethas one, Remdesivir | 32 | Diabetes, Chronic Kidney disease, COPD Obesity | Complications of COVID-19 | Diffuse alveolar damage, overlapping features of exudative and proliferative phase |
| COVID sample#4 | 74 | male | | 25 | Diabetes, Dementia, Coronary artery disease Hyperlipidemia, | Complications of COVID-19 | Diffuse alveolar damage, predominantly proliferative phase |
| Non-COVID control #1 | 74 | male | n/a | n/a | Chronic obstructive pulmonary disease | Chronic obstructive pulmonary disease and pneumonia | Pneumonia |
| Non-COVID control #2 | 67 | male | n/a | n/a | Myelodysplastic syndrome | Pneumonia | Diffuse fibrinous and organizing pneumonia |
| Non-COVID control # 3 | 50 | male | n/a | n/a | Chronic kidney disease, sarcoidosis, chronic pancreatitis | Cryptococcal pneumonia | Cryptococcal pneumonia |

**Cytokine, chemokine, and IgG quantification**

Human IL-18 (Sigma or RND), human CXCL10 (RND), human IL-1RA(Abcam), Human Gasdermin D (MyBioSource) were quantified from supernatants of BMDM infected (or not) with SARS-CoV-2-mNG or from serum or lung homogenates of SARS-CoV-2-mNG infected (or not) MISTRG6 or MISTRG6hACE2 mice following manufacturer's instructions. Human IL-1β was quantified from supernatants of BMDM infected with SARS-CoV-2-mNG using cytometric bead array for human IL-1B (BD) following manufacturer's instructions. Human anti-Spike-RBD IgG (Biolegend) was quantified from sera and lung homogenates of infected or uninfected MISTRG6-hACE2 mice.

**Antibodies**

*Flow cytometry:*

All antibodies used in flow cytometry were obtained from Biolegend, unless otherwise specified.

Antibodies against the following antigens were used for characterization or isolation of cells by flow cytometry:

*Mouse antigens:*

CD45(Clone: 30-F11, Cat #103130), CD45(Clone: 30-F11, Cat #103108), CD45(Clone, 30-F11, Cat #103147), CD326(Clone: G8.8, Cat #118218), F4/80 (Clone: BM8, Cat #123117).

Human antigens: CD45(Clone: HI30, Cat #304044), CD45(Clone: HI30, Cat #304029), CD3(Clone: UCHT1, Cat #300408), CD14(Clone: HCD14, Cat #325620),CD16(Clone: 3G8, Cat #302030), CD16(Clone: 3G8, Cat #302006), CD19(Clone: HIB19, Cat #302218), CD19(Clone: HIB19, Cat #302226), CD33(Clone: WM53, Cat #983902), CD20(Clone: 2H7, Cat #302313), CD20(Clone: 2H7, Cat #302322), CD206(Clone: 15-2, Cat #321106), CD206(Clone: 15-2, Cat #321109), CD86(Clone: BU63, Cat #374210), CD123(Clone: 6H6, Cat #306006), CD11B(Clone: M1/70, Cat #101242), CD11C(Clone: 3.9, Cat #301608), HLA-DR(Clone: LN3, Cat #327014), HLA-DR(Clone: LN3, Cat #327020), HLA-DR(Clone: LN3, Cat #327005), CD183(Clone: G025H7, Cat #353720), CD335-NKp46(Clone: 9E2, Cat #331916), CD4(Clone: OKT4, Cat #317440), CD8(Clone: SK1, Cat #344718), CD8(Clone: SK1, Cat #344748), CD68(Clone: YI/82A, Cat #333828).

*Immunofluorescence:*

Anti-dsRNA antibody (Clone: rJ2,) was purchased from Sigma (Cat# MABE1134) or Antibodies online (Cat# Ab01299-23.0). Polyclonal SARS-CoV-2 RNA-dependent RNA Polymerase antibody was purchased from CellSignaling (Cat # 67988S). Monoclonal SARS-CoV-2 RNA-dependent RNA Polymerase antibody was purchased from Kerafest (Cat# ESG004). Anti-Spike (Spike 1) antibody (clone: 1A9, Cat# GTX632604) was obtained from GeneTex. Anti-Spike (Spike 2) antibody (clone: T01Khu, Cat# 703958) was obtained from ThermoFisher.

*Image Flow Cytometry:*
Mouse anti-human PE-Cy7 CD16(Clone 3G8) was purchased from Biolegend (Cat# 302016). Rabbit anti-human ASC(Polyclonal) was purchased from Santa Cruz (Cat# sc-22514-R). Goat anti-human NLRP3(Polyclonal) was purchased from Abcam(Cat# ab4207). Donkey anti-Rabbit IgG (H+L) Highly Cross-Adsorbed Secondary Antibody(Polyclonal) was purchased from ThermoFisher (Cat# A-31573). Donkey anti-Rabbit IgG (H+L) Cross-Adsorbed Secondary Antibody (Polyclonal) was purchased from ThermoFisher (Cat# A-10040). Donkey anti-Goat IgG (H+L) Cross-Adsorbed Secondary Antibody(Polyclonal) was purchased from ThermoFisher(Cat# A-21447).

*Therapeutic antibodies:*
Monoclonal antibody against human CD16 used in blocking experiments were purchased from Abcam (SP175). Monoclonal antibody against human ACE2 was purchased from Abcam. Anti-CD16 antibody used in depletion experiments was purchased from ThermoFisher (3G8). Monoclonal antibodies (clones 135 and 144) were acquired from M. Nussenzweig as has been previously described[45]. Anti-IFNAR2 antibody was purchased from PBL Assay science (Cat #21385-1).

**Gene expression**
RNA was extracted with the RNeasy mini kit (Qiagen) per the manufacturer's protocol High-Capacity cDNA Reverse Transcription Kit was used to make cDNA. Quantitative reverse transcription PCR (qRT-PCR) was performed using an SYBR FAST universal qPCR kit (KAPA Biosystems). Predesigned KiCqStart primers for *DDX58, IL6, IFITM3, IRF7, IFIH1, IFNA6, IFNG* and *HPRT1* were purchased from Sigma.

**Bulk whole tissue RNA-sequencing**
RNA isolated from homogenized lung tissue, also used for viral RNA analysis, was prepared for whole tissue transcriptome analysis using low input (14dpi) or conventional (28dpi) bulk RNA sequencing. Libraries were made with the help of the Yale Center for Genomic Analysis. Briefly, libraries were prepared with an Illumina rRNA depletion kit and sequenced on a NovaSeq. Raw sequencing reads were aligned to the human-mouse combined genome with STAR[86], annotated and counted with HTSeq[87], normalized using DESeq2[88] and graphed using the Broad Institute Morpheus web tool. Differential expression analysis was also performed with DESeq2. For IFN-stimulated gene identification, http://www.interferome.org was used with parameters -*In Vivo*, -Mus musculus or Homo sapiens -fold change up 2 and down 2.

**Single Cell RNA Sequencing 10X Genomics**

Sorted human lung immune cells (hCD45+ in uninfected, 14dpi and 28dpi) were stained with TotalSeq (TotalSeq™-B0251 anti-human Hashtag 1 Antibody: GTCAACTCTTTAGCG; TotalSeq™-B0252 anti-human Hashtag 2 Antibody: TGATGGCCTATTGGG) antibodies (Biolegend) prior to processing for droplet based scRNA-seq. 10X Chromium GEM technology. Single cell transcriptomes and associated protocols of 4dpi lungs (total lung cells as opposed to sorted human immune cells analyzed) were previously described[19]. Duplicates from each condition/time point were pooled in equal numbers to ensure 10000 cells were encapsulated into droplets using 10X Chromium GEM technology. Libraries were prepared in house using Chromium Next GEM Single Cell 3' Reagent Kits v3.1 (10X Genomics). scRNA-seq libraries were sequenced using Nova-Seq. Raw sequencing reads were processed with Cell Ranger 3.1.0 using a human-mouse combined reference to generate a gene-cell count matrix. To distinguish human and mouse cells, we counted the number of human genes (nHuman) and mouse genes (nMouse) with nonzero expression in each cell, and selected cells with nHuman > 20 * nMouse as human cells. The count matrix of human cells and human genes was used in the downstream analysis with Seurat 3.2[62]. Specifically, this matrix was filtered to remove low quality cells, retaining cells with > 200 and < 5,000 detected genes and < 20% mitochondrial transcripts. We then log normalized each entry of the matrix by computing log (CPM/100 + 1), where CPM stands for counts per million. To visualize the cell subpopulations in two dimensions, we applied principal component analysis followed by t-SNE, a nonlinear dimensionality reduction method, to the log-transformed data. Graph-based clustering was then used to generate clusters that were overlaid on the t-SNE coordinates to investigate cell subpopulations. Marker genes for each cluster of cells were identified using the Wilcoxon test (two-tailed) with Seurat. For the adjusted P values the Bonferroni correction was used. In this analysis, uninfected: 438 cells, 4dpi: 336 cells, 14dpi: 793 cells, 28dpi: 1368 cells were included.

To identify differentially abundant (DA) subpopulations not restricted to clusters, we applied DA-seq[61], a targeted, multiscale approach that quantifies a local DA measure for each cell for comprehensive and accurate comparisons of transcriptomic distributions of cells. DA measure defined by DA-seq. shows how much a cell's neighborhood is enriched by the cells from either uninfected or infected lungs. DA-seq analysis where on our data revealed that T cells, monocytes and macrophages were responsible for most of the chronic infection driven changes. Red coloring signify enrichment at 28dpi lungs and blue coloring mark enrichment in uninfected lungs

To combine cells from different DPIs (uninfected, 4dpi, 14dpi, 28dpi), we applied the integration method[62] in Seurat to remove batch effects. We then performed principal component analysis and retained top 30 PCs as the input to tSNE, a nonlinear dimensionality reduction method, to embed the data onto 2-dimensional space for visualization. Graph-based clustering with a resolution of 0.8 was then used to generate clusters that were overlaid on the t-SNE coordinates to investigate cell subpopulations. Marker genes for each cluster of cells were identified using the Wilcoxon test (two-tailed) with Seurat (For the adjusted P values the Bonferroni correction was used). After cell type identification, we separated out macrophage populations from all DPIs, and applied the same procedures as described above to re-preprocess and visualize the data. Clusters were redefined based on a resolution of 0.3.

**Statistics and Reproducibility**

Unpaired or paired t-test (always two-tailed) was used to determine statical significance for changes in immune cell frequencies and numbers while comparing infected mice to uninfected control mice or treated mice to untreated mice. To determine whether the viral RNA quantification is statistically significant across treatment groups or timepoints, Mann-Whitney, two-tailed test was used. Wilcoxon test (two-tailed) or ratio paired t-test (two-tailed) was used to determine whether the viral titer quantification of the untreated condition is significantly different from the treated groups. For Pearson's test, significance was based on the t-test. The test statistic is based on Pearson's product-moment correlation coefficient cor(x, y) and follows a t distribution with length(x)-2 degrees of freedom. For Spearman's test, p-values are computed using algorithm AS 89 with exact = TRUE. All micrographs presented in the study were representative of at least 3 animals or specimens. Each experiment was repeated independently at least two times. All attempts yielded similar results. In *in vivo* studies, each dot represents a biologically independent mouse.

#### Supplementary Figures

**Figure S1, A-L: Anti-IFNAR2 and Remdesivir therapy reverses infection induced transcriptional changes (matched to figure 1).**

**Figure S2, A-F. Viral replication products are detected in human macrophages (matched to figure 3).**

**Figure S3, A-I. SARS-CoV-2 infection of human macrophages activates inflammasomes and leads to a unique inflammatory transcriptional signature *in vivo* (matched to figure 4).**

**Figure S4, A-D. *ACE2* expression is inducible and highly correlates with genes that are associated with type I interferon signaling.**

The data supporting this publication and presented as part of the supplementary information here is available at Figshare.com under the project entitled: "Viral replication in human macrophages enhances an inflammatory cascade and interferon driven chronic COVID-19 in humanized mice," (https://doi.org/10.6084/m9.figshare.19401335).

Figure S1

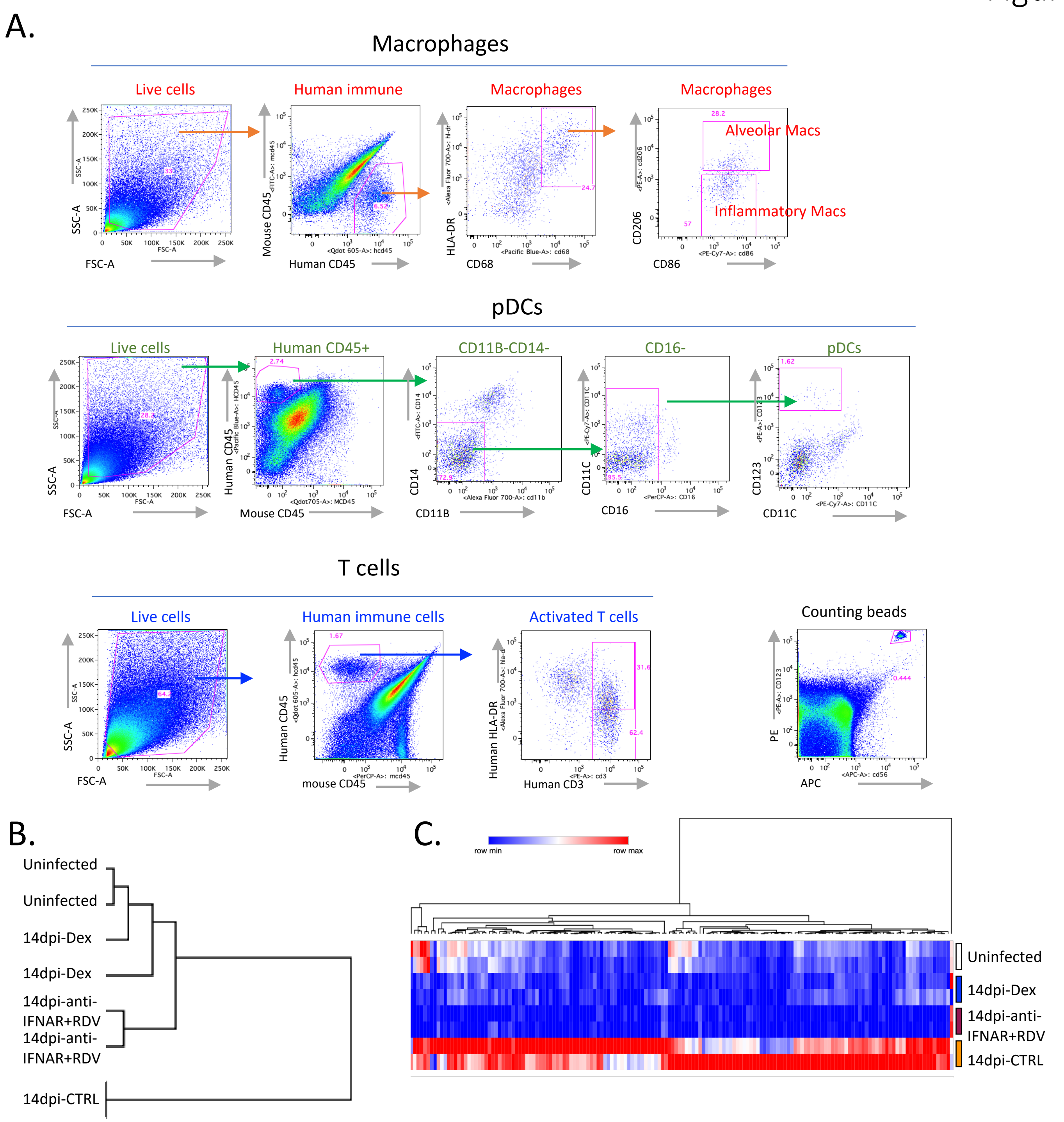

**Figure S1: Anti-IFNAR2 and Remdesivir therapy reverses infection induced transcriptional changes (matched to figure 1).**

A. Representative gating strategy of human immune cells in the lungs of SARS-CoV-2 infected MISTRG6-hACE2 mice. Cells isolated from lungs or bronchioalveolar lavage (BAL) were stained with antibodies against human CD45, HLA-DR, CD68, CD16, CD14, CD206, CD86, CD11B, CD11C, CD123, CD3, and mouse CD45. Cell numbers were calculated using counting beads.

B. Similarity comparison of uninfected, infected, and therapeutically manipulated lungs based on dexamethasone suppressed genes. Pearson correlation. Duplicates analyzed for each condition.

C. Genes suppressed by both dexamethasone and combined therapy of Remdesivir and anti-IFNAR2 (Log2, Foldchange <-1, P adj<0.05). P adj: For the adjusted P values the Bonferroni correction was used. Duplicates analyzed for each condition. See Table S1 for a full list of genes and their normalized expression.

D.

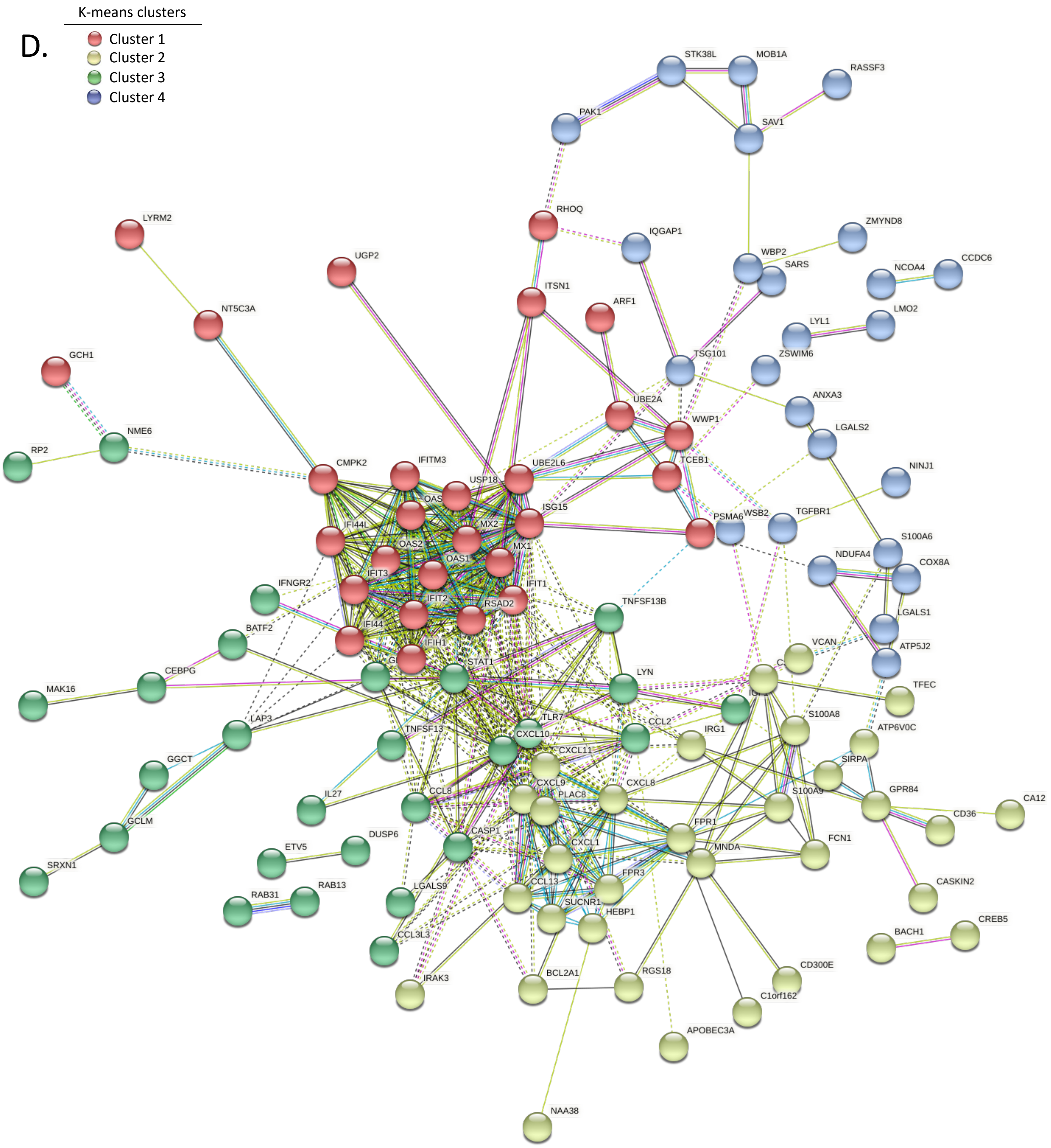

**Figure S1: Anti-IFNAR2 and Remdesivir therapy reverses infection induced transcriptional changes (matched to figure 1).**
D. Network analysis (STRING v11.0) of genes suppressed by both dexamethasone and combined therapy of Remdesivir and anti-IFNAR2 (as shown in S2B). Duplicates analyzed for each condition. K-means clustering (n=4).

E.

| Name | p-value | Overlap | |
|---|---|---|---|
| Role of Hypercytokinemia/hyperchemokinemia in the Pathogenesis of Influenza | 5.55E-16 | 16.3 % | 14/86 |
| Interferon Signaling | 8.29E-11 | 22.2 % | 8/36 |
| Granulocyte Adhesion and Diapedesis | 5.44E-09 | 6.3 % | 12/189 |
| Role of IL-17F in Allergic Inflammatory Airway Diseases | 5.18E-07 | 13.3 % | 6/45 |
| Role of Pattern Recognition Receptors in Recognition of Bacteria and Viruses | 1.05E-06 | 5.8 % | 9/156 |

F.

Cluster identifying markers (Fig. 2b)

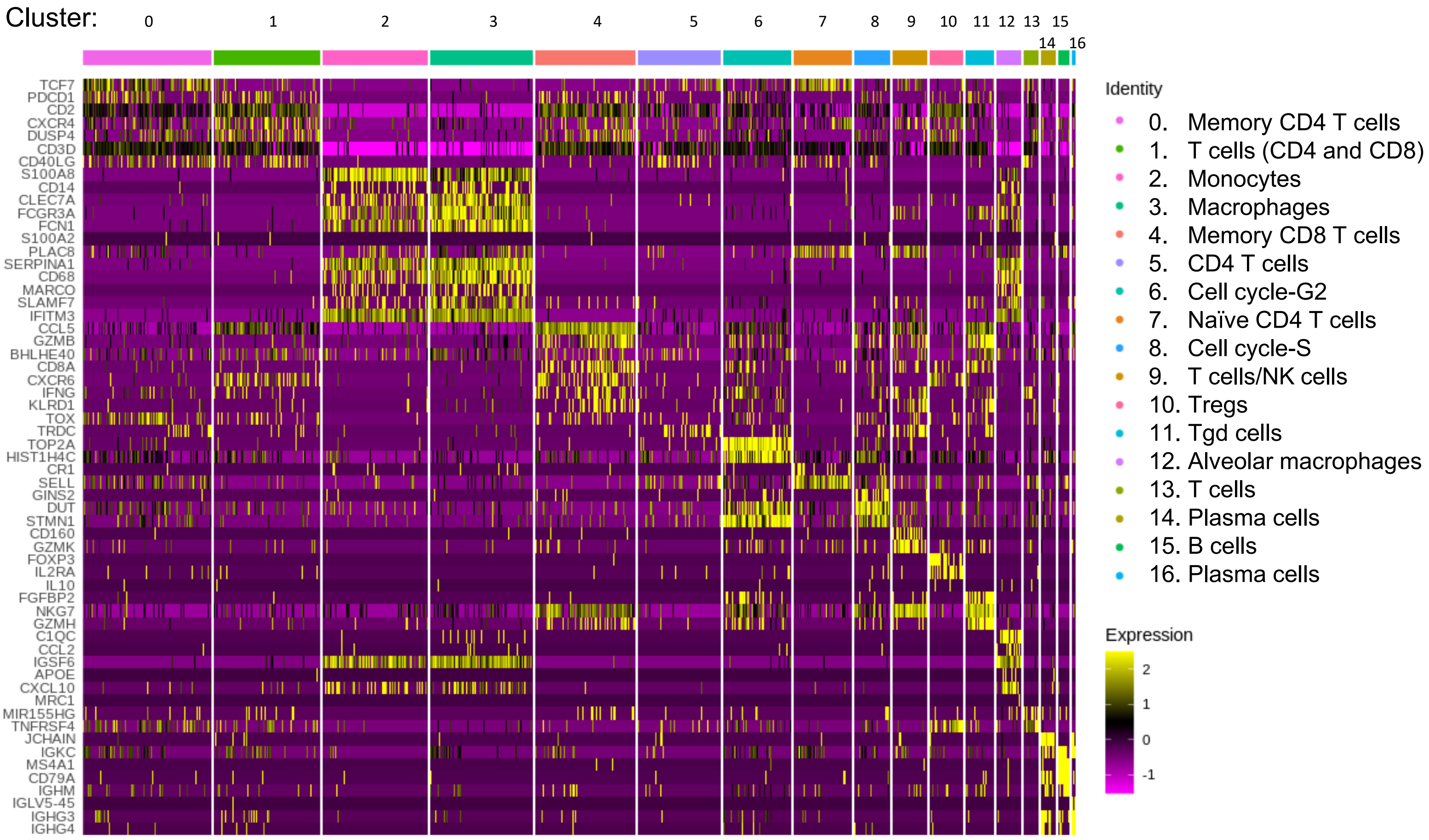

**Figure S1: Anti-IFNAR2 and Remdesivir therapy reverses infection induced transcriptional changes (matched to figure 1).**
E. Pathway (Ingenuity) analysis of genes suppressed by both dexamethasone and combined therapy of Remdesivir and anti-IFNAR2 (as shown n in S2B). Duplicates analyzed for each condition. Fisher's Exact Test was used to determine statistical significance in the overlap between the dataset genes and the genes suppressed by therapy.
F. Cluster identifying genes comparing human immune cells from infected (28dpi) or uninfected lungs for 17 clusters shown in Fig 1d. Marker genes for each cluster of cells were identified using the Wilcoxon test with Seurat. Pooled duplicates analyzed for each condition.

G

Cluster specific markers (Fig. 1g)

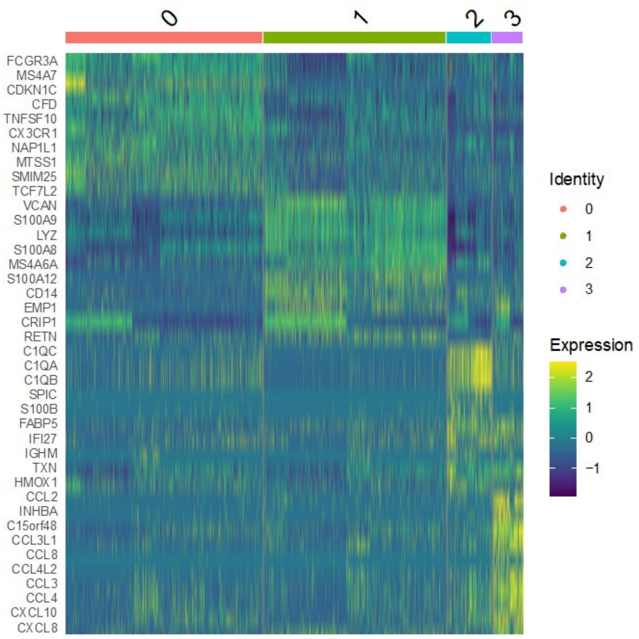

H.

DPI-specific markers (Fig. 1g)

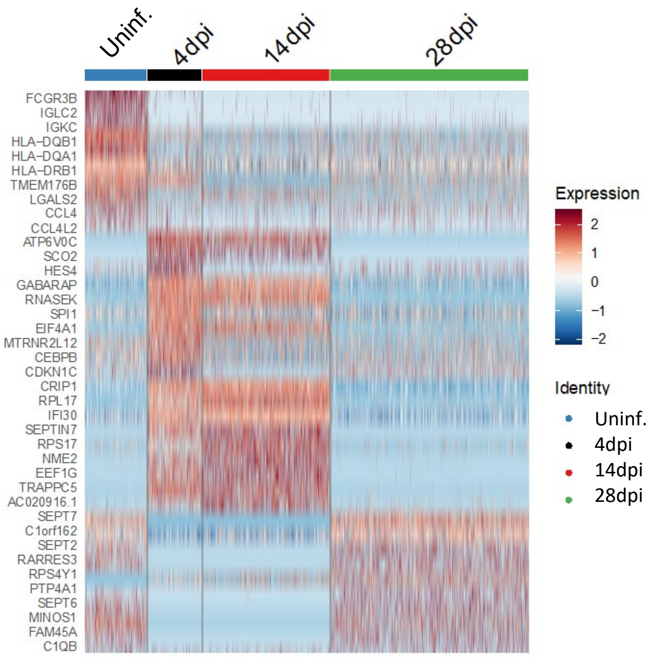

**Figure S1: Anti-IFNAR2 and Remdesivir therapy reverses infection induced transcriptional changes (matched to figure 1).**

G. Heatmap visualizing cluster identifying genes comparing human monocytes and macrophages from infected (4, 14 or 28dpi) or uninfected lungs. Pooled duplicates. Uninfected: 438 cells, 4dpi: 336 cells, 14dpi: 793 cells, 28dpi: 1368 cells were analyzed. Marker genes for each cluster of cells were identified using the Wilcoxon test (two-tailed) with Seurat.

H. Temporal distribution of transcriptional changes associated with monocytes and macrophages in infected (4, 14 or 28dpi) or uninfected lungs. Pooled duplicates analyzed. Uninfected: 438 cells, 4dpi: 336 cells, 14dpi: 793 cells, 28dpi: 1368 cells included in analysis.

Figure S1

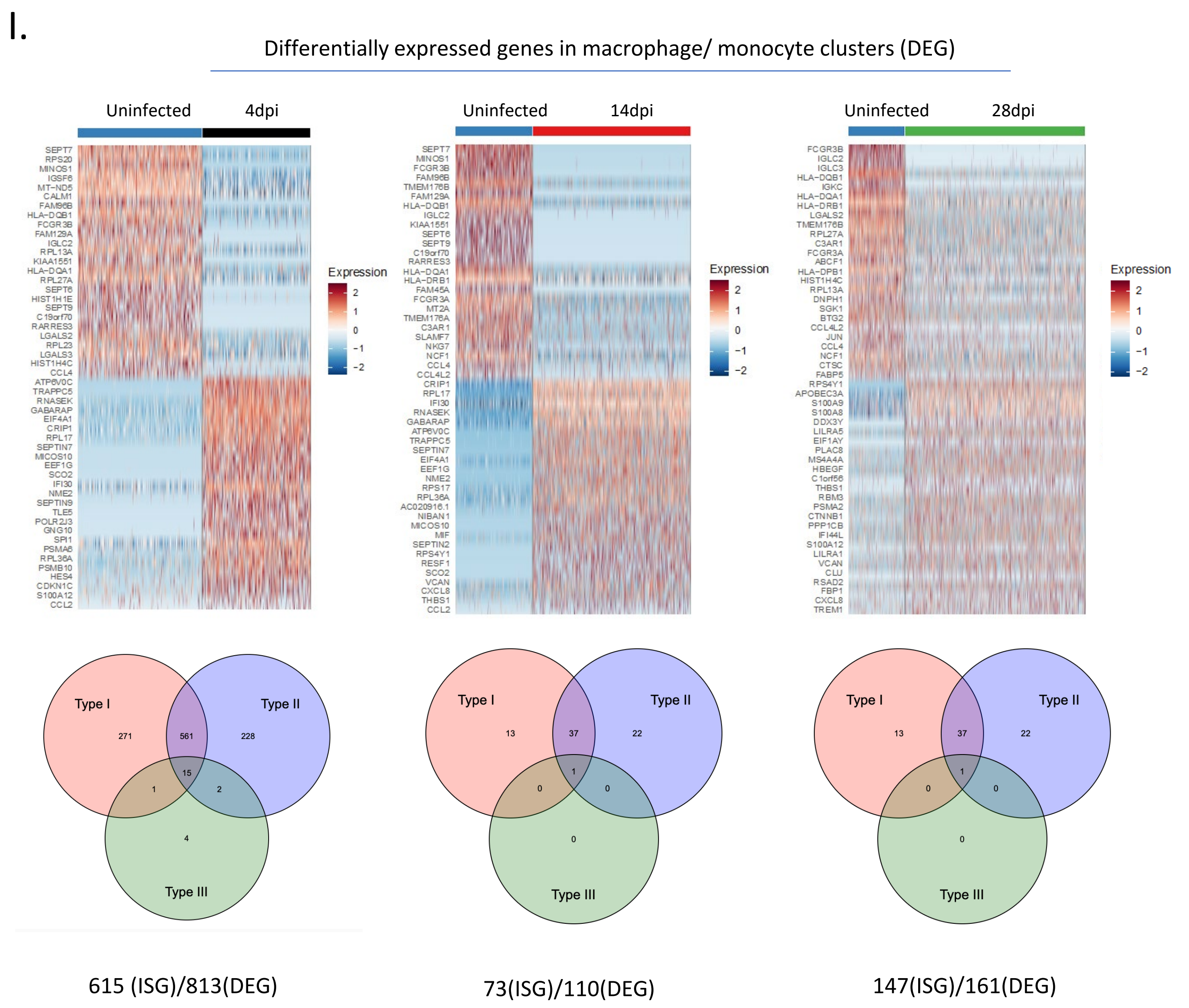

**Figure S1: Anti-IFNAR2 and Remdesivir therapy reverses infection induced transcriptional changes (matched to figure 1).**

I. Top: Heatmap of representative genes that are differentially regulated (DEGs) in human macrophages from 4, 14, 28dpi lungs compared with uninfected lungs. Uninfected: 438 cells, 4dpi: 336 cells, 14dpi: 793 cells, 28dpi: 1368 cells included in analysis. Bottom: Distribution of interferon stimulated genes within these DEGs. Pooled duplicates analyzed .

### Figure S1

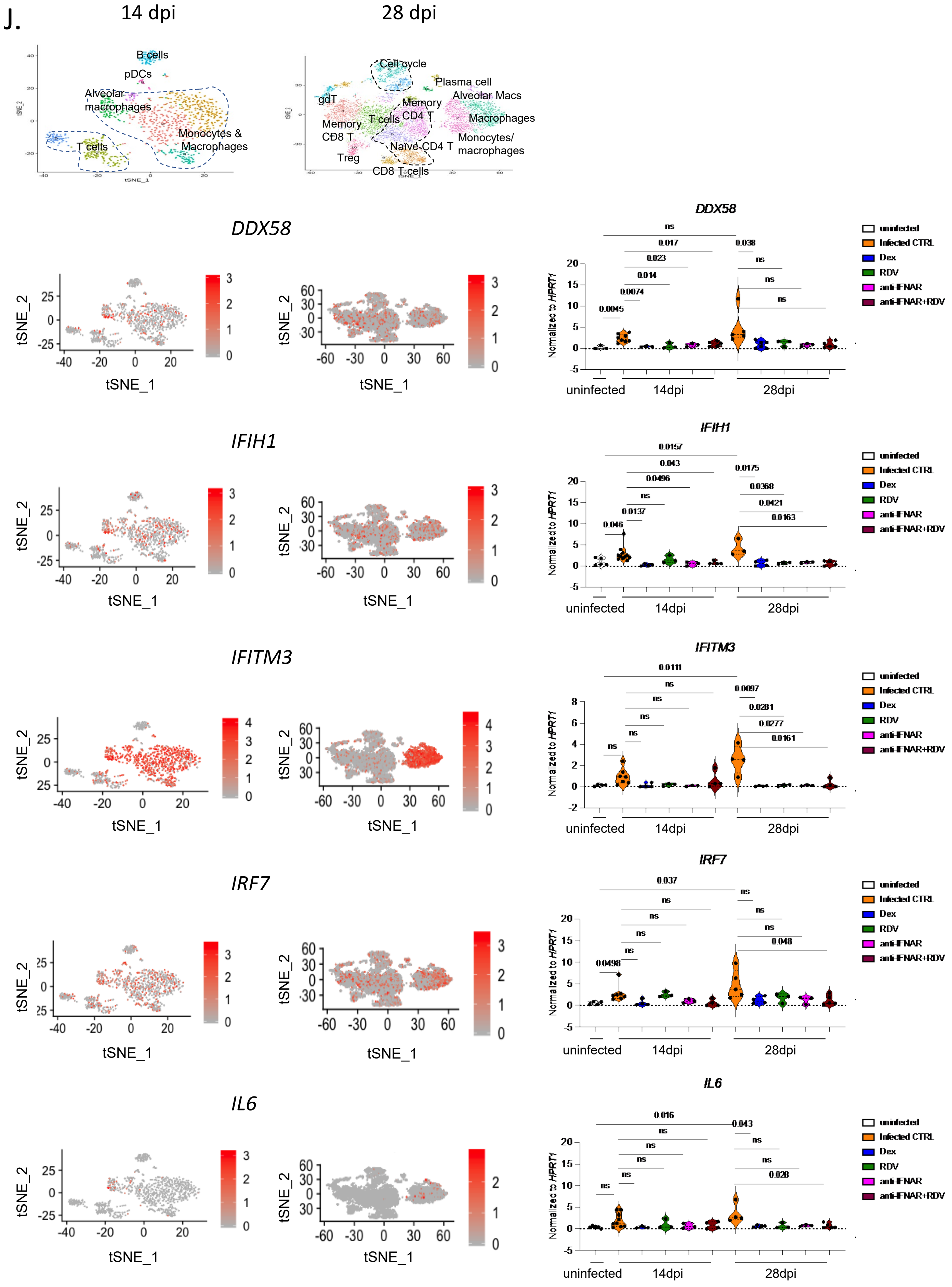
J.
14 dpi
28 dpi
B cells
pDCs
Alveolar macrophages
T cells
Monocytes & Macrophages
Cell cycle
Plasma cell
Alveolar Macs
gdT
Memory CD4 T
T cells
Memory CD8 T
Macrophages
Monocytes/ macrophages
Naïve CD4 T
Treg
CD8 T cells
DDX58
IFIH1
IFITM3
IRF7
IL6
tSNE_1
tSNE_2
Normalized to HPRT1
uninfected
14dpi
28dpi
Infected CTRL
Dex
RDV
anti-IFNAR
anti-IFNAR+RDV
ns
0.0045
0.0074
0.014
0.023
0.017
0.038
0.046
0.0137
0.0496
0.043
0.0157
0.0175
0.0368
0.0421
0.0163
0.0111
0.0097
0.0281
0.0277
0.0161
0.0498
0.037
0.048
0.016
0.043
0.028

**Figure S1: Anti-IFNAR2 and Remdesivir therapy reverses infection induced transcriptional changes (matched to figure 1).**
J. Relative expression of interferon inducible or inflammatory genes in treated or untreated MISTRG6-hACE2 mice infected with SARS-CoV-2 mice at 14dpi or 28dpi. Uninfected baseline expression values are presented as reference. The distribution of cells that preferentially express these genes is overlayed on the tSNE plots showing 14dpi and 28dpi human immune cells. *IFITM3* and *IL6* were particularly enriched in macrophage/monocyte clusters, while *IRF7*, *DDX58* and *IFIH1* were enriched in multiple immune cells such as T cells, B cells, and myeloid cells. Normalized to *HPRT1*. *DDX58*: uninfected n=3; CTRL-infected: 14dpi n=8, 28dpi n=: 5; Dex 14dpi n=3, 28dpi n= 6; RDV 14 and 28dpi n=3; anti-IFNAR2 14 and 28 dpi n=3, anti-IFNAR2+ Remdesivir 14 and 28dpi n=5 biologically independent mice examined over at least 2 independent experiments. *IFIH1*: uninfected n=5; CTRL-infected: 14dpi n =11, 28dpi n=: 3; Dex 14 and 28dpi n=4; RDV 14 and 28dpi n=3; anti-IFNAR2 14 and 28 dpi n=3, anti-IFNAR2+ Remdesivir 14 dpi n=4 and 28dpi n=5 biologically independent mice examined over at least 2 independent experiments. *IFITM3*: uninfected n=4; CTRL-infected: 14dpi n =7, 28dpi n=: 4; Dex 14 and 28dpi n=4; RDV 14 and 28dpi n=3; anti-IFNAR2 14 and 28 dpi n=3, anti-IFNAR2+ Remdesivir 14 and 28dpi n=4 biologically independent mice examined over at least 2 independent experiments. *IRF7*: uninfected n=4; CTRL-infected: 14dpi n =7, 28dpi n=: 5; Dex 14 and 28dpi n=4; RDV 14 and 28dpi n=3; anti-IFNAR2 14 and 28 dpi n=3, anti-IFNAR2+ Remdesivir 14dpi n=4, 28dpi n=5 biologically independent mice examined over at least 2 independent experiments. *IL6*: uninfected n=5; CTRL-infected: 14dpi n =9, 28dpi n=: 4; Dex 14dpi n=3, 28dpi n=4; RDV 14 and 28dpi n=3; anti-IFNAR2 14 and 28 dpi n=3, anti-IFNAR2+ Remdesivir 14dpi n=4, 28dpi n=5 biologically independent mice examined over at least 2 independent experiments. Unpaired, two-tailed t-test.

Figure S1

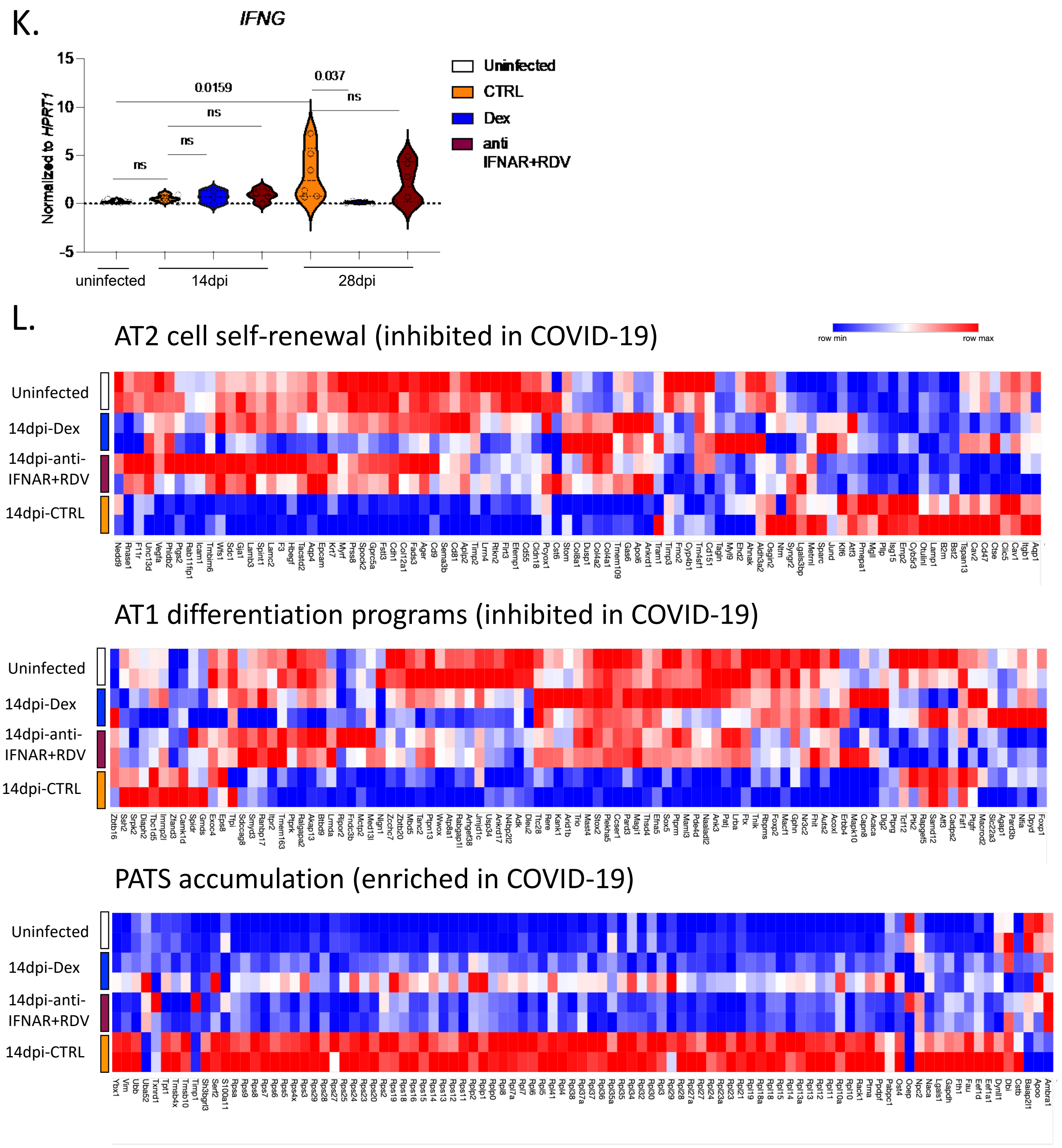

**Figure S1: Anti-IFNAR2 and Remdesivir therapy reverses infection induced transcriptional changes (matched to figure 1).**

K. Relative expression of *IFNG* in treated or untreated MISTRG6-hACE2 mice infected with SARS-CoV-2 mice at 14dpi or 28dpi. Uninfected baseline expression values are presented as reference. Normalized to *HPRT1*. Uninfected n=7; CTRL infected: 14dpi n =7, 28dpi n=: 6; Dex 14dpi=3, 28dpi= 5; anti-IFNAR2+ Remdesivir 14dpi n=4, 28dpi n=6. over at least 2 independent experiments. Unpaired, two-tailed t-test.

L. Heatmap of AT2 cell self-renewal and AT1 differentiation and pre-alveolar type 1 transitional cell state (PATS) associated genes at in uninfected or infected (14dpi) lungs in response to therapeutics. AT2 cell self-renewal and AT1 differentiation gene signature was inhibited while PATS gene signature was enriched in autopsy lungs of patients with severe COVID-19[7]. Top differentially expressed genes in epithelial cluster 7 of autopsy lungs[7] were used in the analysis. Duplicates were analyzed for each condition. Normalized counts of duplicates visualized as min-max transformed values, calculated by subtracting row mean and diving by SD for each gene. Rows (genes) clustered by hierarchical clustering (one-minus Pearson).

A.

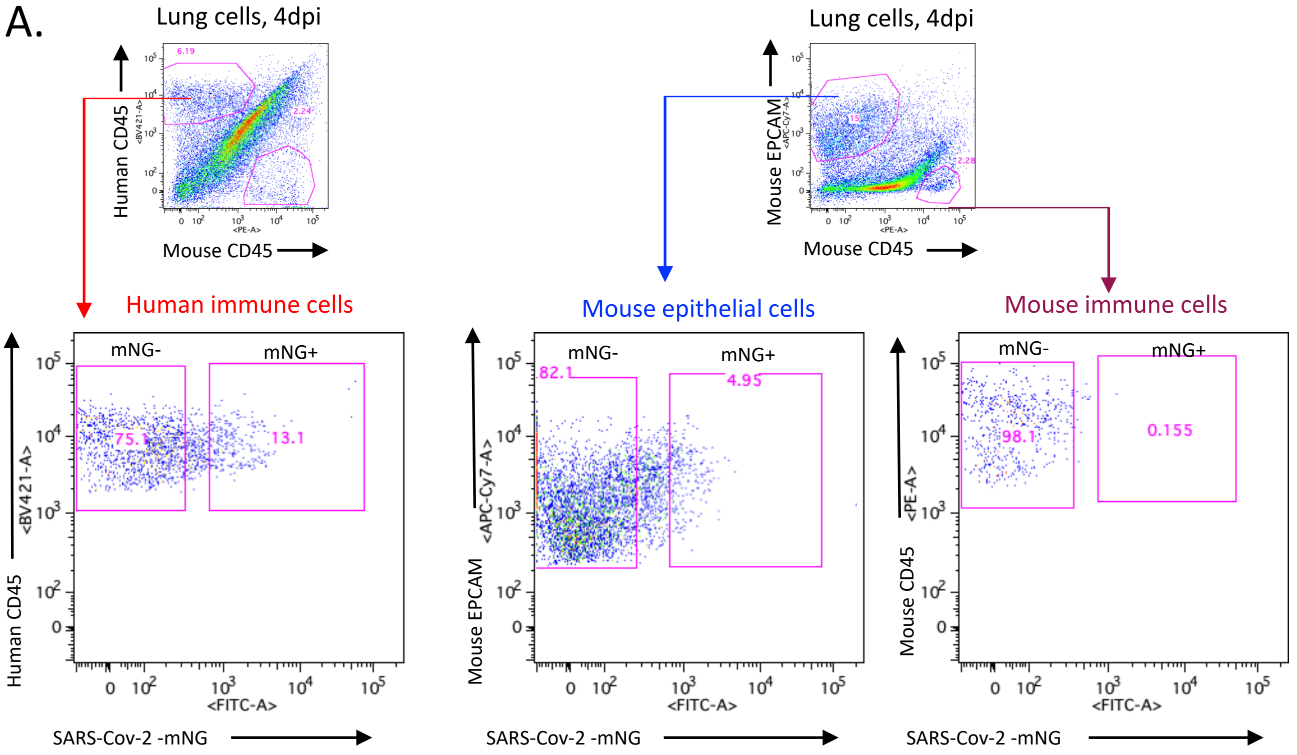

B.

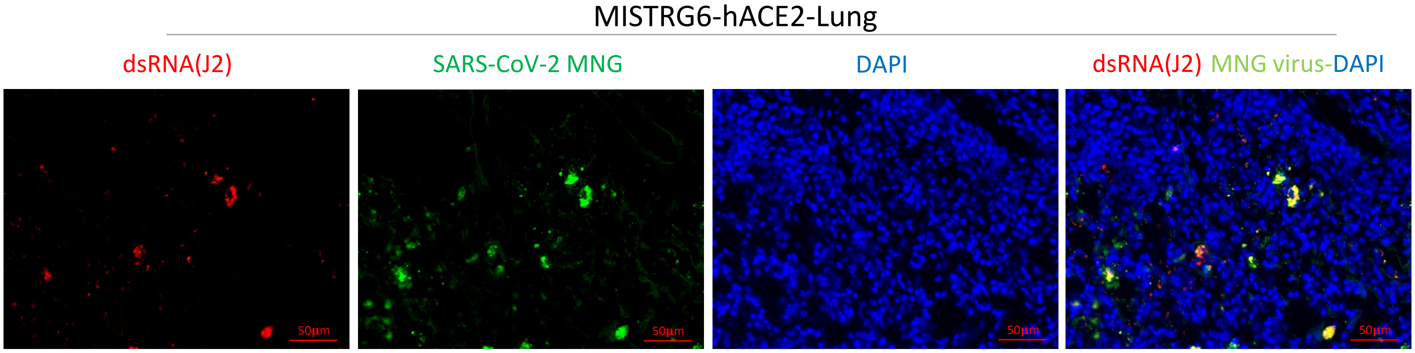

**Figure S2. Viral replication products are detected in human macrophages (matched to figure 2).**
A. Representative gating strategy for sorting mNG+ and mNG- human immune cells, mNG+ and mNG- mouse epithelial cells and mouse immune cells. Lung cells from SARS-CoV2-mNG infected MISTRG6-hACE2 mice were stained with antibodies against human CD45, mouse CD45, and mouse EPCAM. Sorted cells were used for viral quantification (Fig. 2) and characterization of the inflammasome pathway (Fig. 3).
B. Representative fluorescent microscopy images showing colocalization of double stranded RNA (clone rJ2) staining, mNG signal and DAPI staining in fixed lung tissue at 4dpi. Representative of n=4.

Figure S2

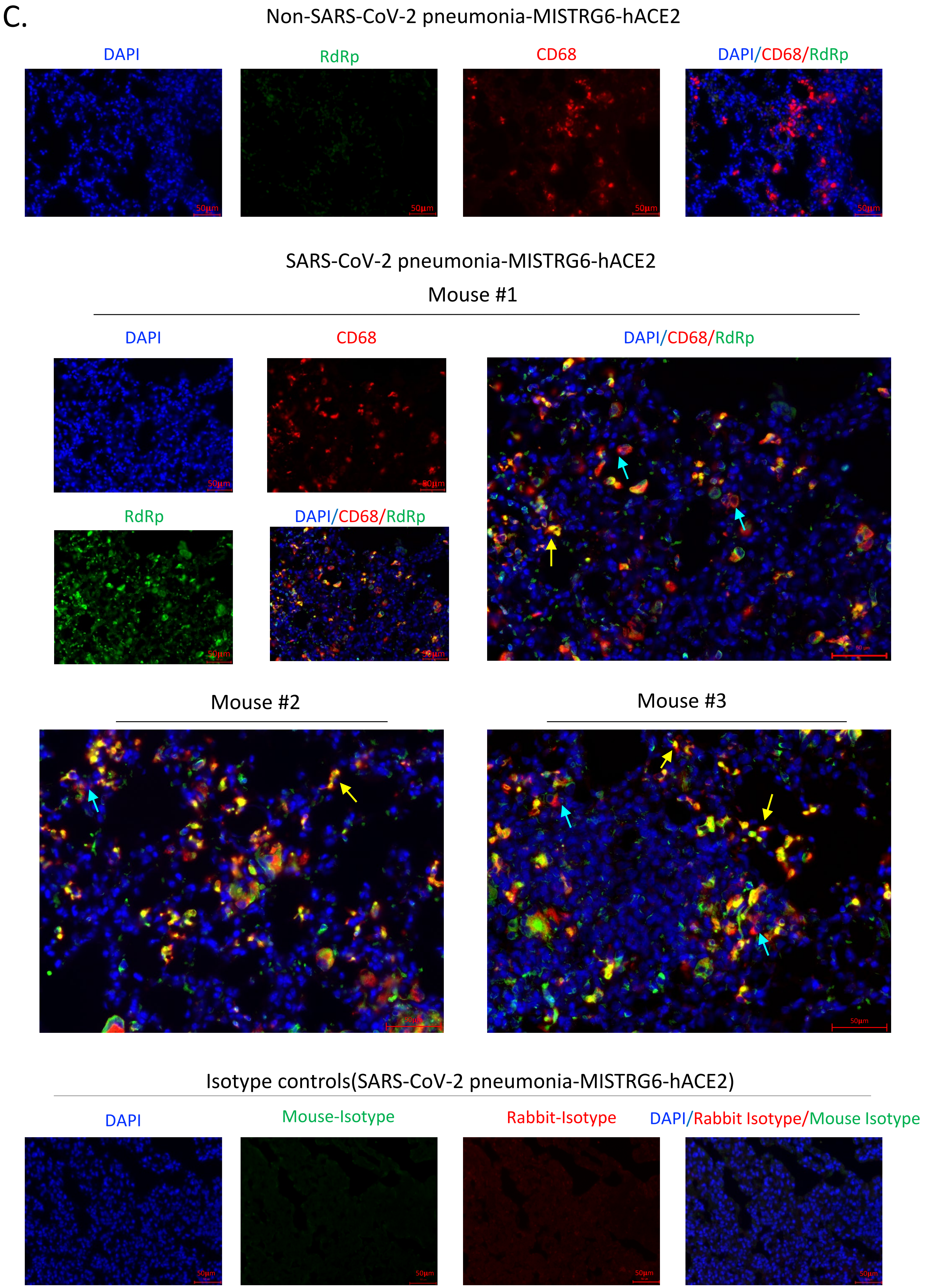
C.
Non-SARS-CoV-2 pneumonia-MISTRG6-hACE2
DAPI
RdRp
CD68
DAPI/CD68/RdRp
50µm
SARS-CoV-2 pneumonia-MISTRG6-hACE2
Mouse #1
DAPI
CD68
DAPI/CD68/RdRp
RdRp
DAPI/CD68/RdRp
Mouse #2
Mouse #3
Isotype controls(SARS-CoV-2 pneumonia-MISTRG6-hACE2)
DAPI
Mouse-Isotype
Rabbit-Isotype
DAPI/Rabbit Isotype/Mouse Isotype

**Figure S2. Viral replication products are detected in human macrophages (matched to figure 2).**
C. Representative fluorescent microscopy images of RNA dependent RNA polymerase (RdRp), anti-human CD68 and DAPI staining in fixed lung tissue from SARS-CoV-2 infected or control MISTRG6-hACE2 mice (Non-SARS-CoV-2 pneumonia). Representative of n=7 biologically independent SARS-CoV-2 infected mice examined over 3 independent experiments. Yellow arrows mark RdRp+ human macrophages. Blue arrows mark RdRp- human macrophages. Isotype controls (bottom panels) and non-COVID pneumonia lungs (bacterial infection, top panels) n=3 biologically independent mice are presented as controls. Pseudo-colors were assigned for visualization.

D.

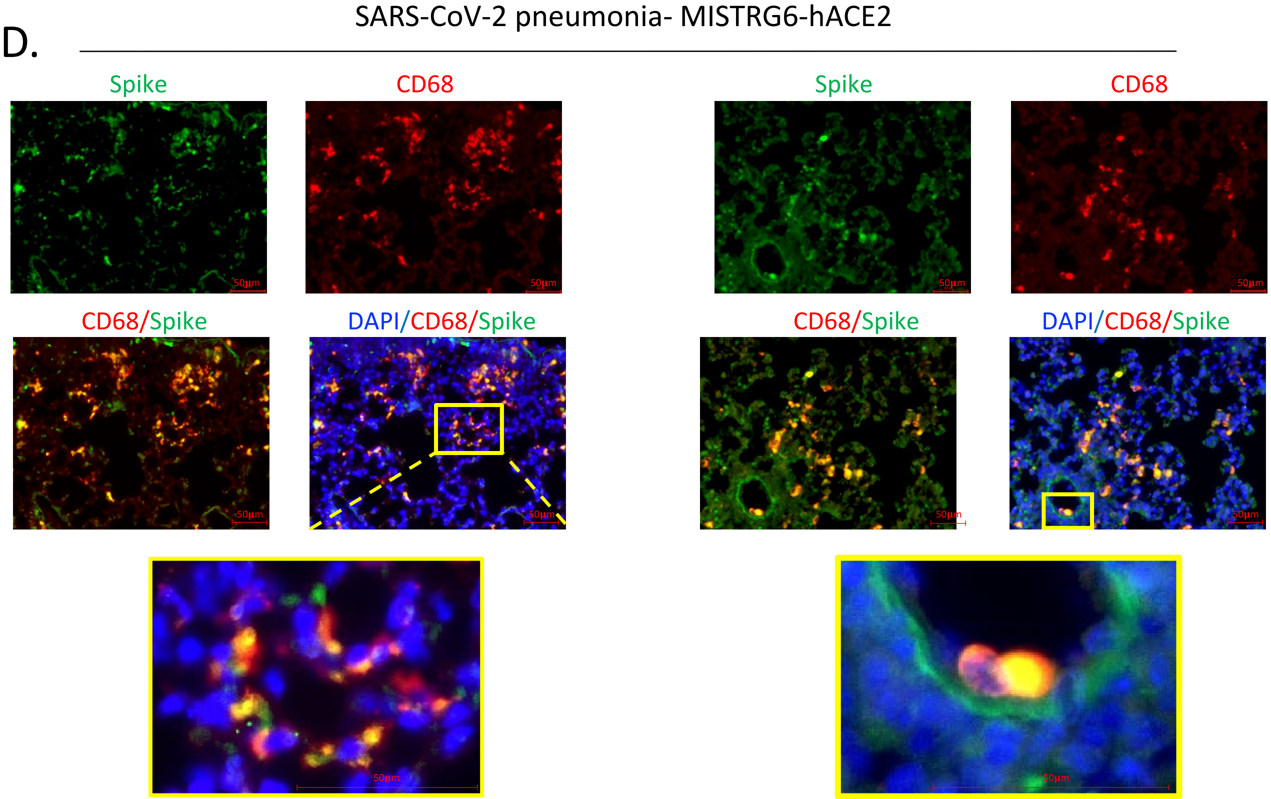

E.

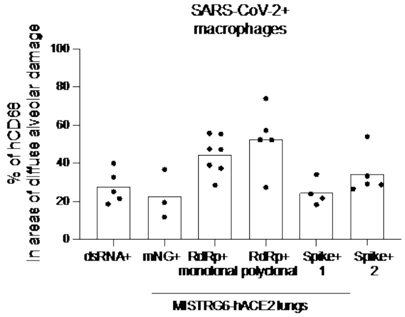

**Figure S2. Viral replication products are detected in human macrophages (matched to figure 2).**
D. Representative fluorescent microscopy images of Spike (S), human CD68, and DAPI staining in fixed lungs of SARS-CoV-2- infected MISTRG6-hACE2 mice. Yellow rectangle provides a higher magnification view of the selected area. Pseudo-colors were assigned for visualization. Representative of n=5 biologically independent mice examined over 3 independent experiments.
E. Quantification of viral replication products or machinery in human lung macrophages from SARS-CoV-2 infected MISTRG6-hACE2 mice measured by immunofluorescent staining. Quantification was performed based on representative high-power images (40x) in areas showing diffuse alveolar damage. Frequencies of dsRNA, mNG, RdRp, and Spike positive human macrophages out of hCD68+DAPI+ cells are plotted. dsRNA: 20, 81, 133, 135, 52 human macrophages were counted. mNG: 30, 103, 110 human macrophages were counted. RdRp monoclonal: 187, 59, 85, 106, 142, 63, 59 human macrophages were counted. RdRp polyclonal: 134, 21, 22, 218, 44 human macrophages were counted. Spike (antibody 1): 21, 22, 218, 44 total human macrophages were counted. Spike (antibody 2): 63, 83, 163, 101, 57 human macrophages were counted. N=5 (dsRNA+), N=3 (mNG), N=7 (RdRp+ monoclonal), N=5 (RdRp+ polyclonal), N=4 (Spike-1), N=5 (Spike-2) biologically independent mice representative of at least 2 independent experiment. Means with all datapoints are shown. See supplementary methods for details of antibodies used.

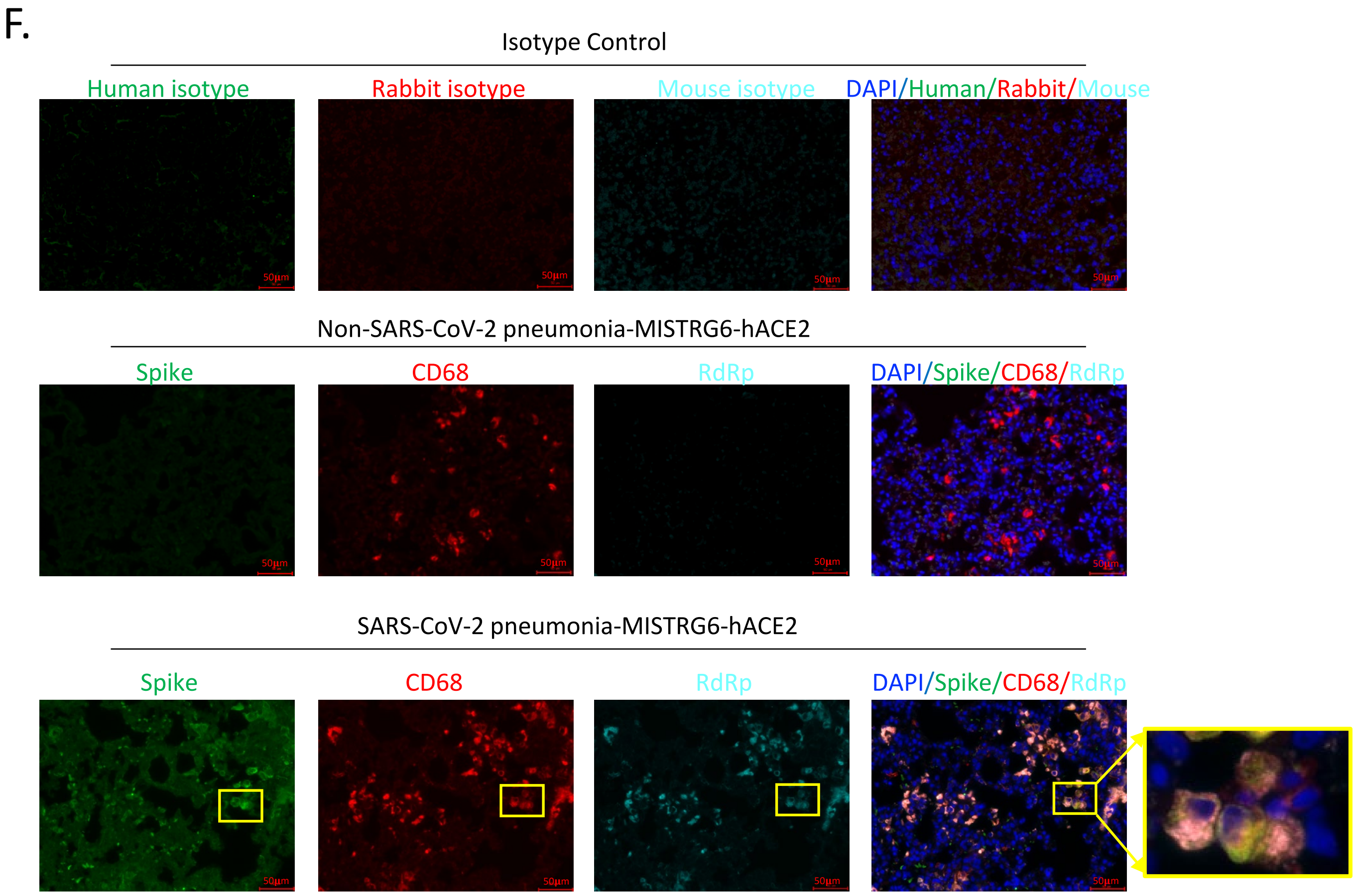

**Figure S2. Viral replication products are detected in human macrophages (matched to figure 2).** F. Representative fluorescent microscopy images and quantification of colocalization of Spike (S), RNA dependent RNA polymerase (RdRp), human CD68, and DAPI staining in fixed lungs of SARS-CoV-2-infected MISTRG6-hACE2 mice. Top panel: isotype control staining of SARS-CoV-2- infected lungs. Middle panel: control lungs with Non-SARS-CoV-2, bacterial pneumonia. Bottom panels: SARS-CoV-2-infected MISTRG6-hACE2 mice. Yellow rectangle provides a higher magnification view of the selected area. Pseudo-colors are assigned for visualization. Representative of n=5 biologically independent mice over 3 independent experiments.

Figure S3

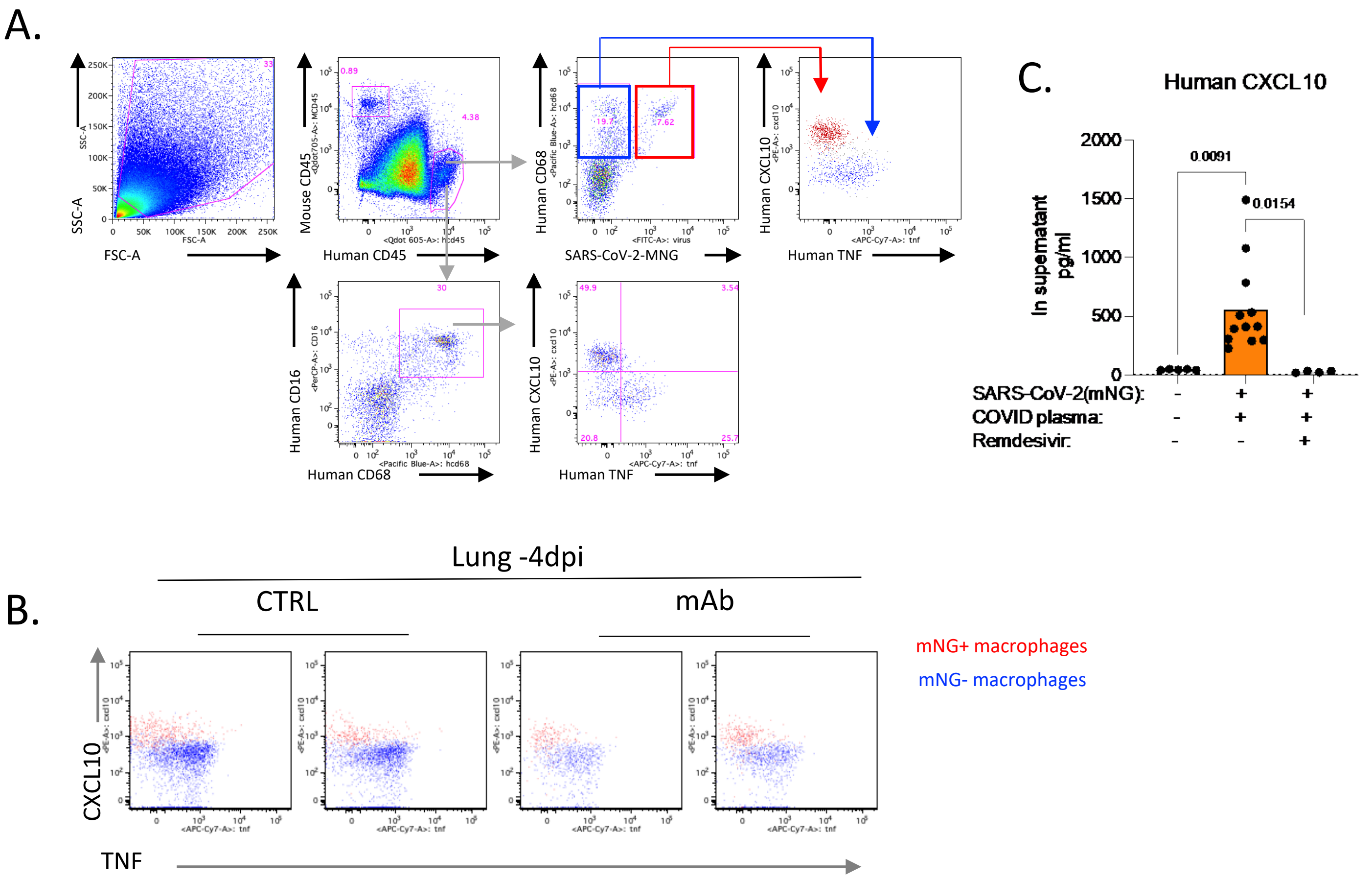

**Figure S3. SARS-CoV-2 infection of human macrophages activates inflammasomes and leads to a unique inflammatory transcriptional signature in vivo (matched to figure 3).**
A. Representative gating strategy of CXCL10 or TNF producing human macrophages in MISTRG6-hACE2 mice infected with SARS-CoV-2-mNG.
B. Representative flow cytometry plots of CXCL10 and TNF staining in mice therapeutically treated with mAb or control untreated mice. Representative of n=4 biologically independent mice.
C. CXCL10 production measured by ELISA in supernatants of BM-macrophages infected with SARS-CoV-2 in vitro. Infected BM-macrophage cultures were supplemented with pooled plasma from COVID-19 and were treated with Remdesivir or not. Uninfected n=5, CTRL infected n=12, RDV n=4 over 3 independent experiments. Means with individual values are plotted. Unpaired, two-tailed t-test.

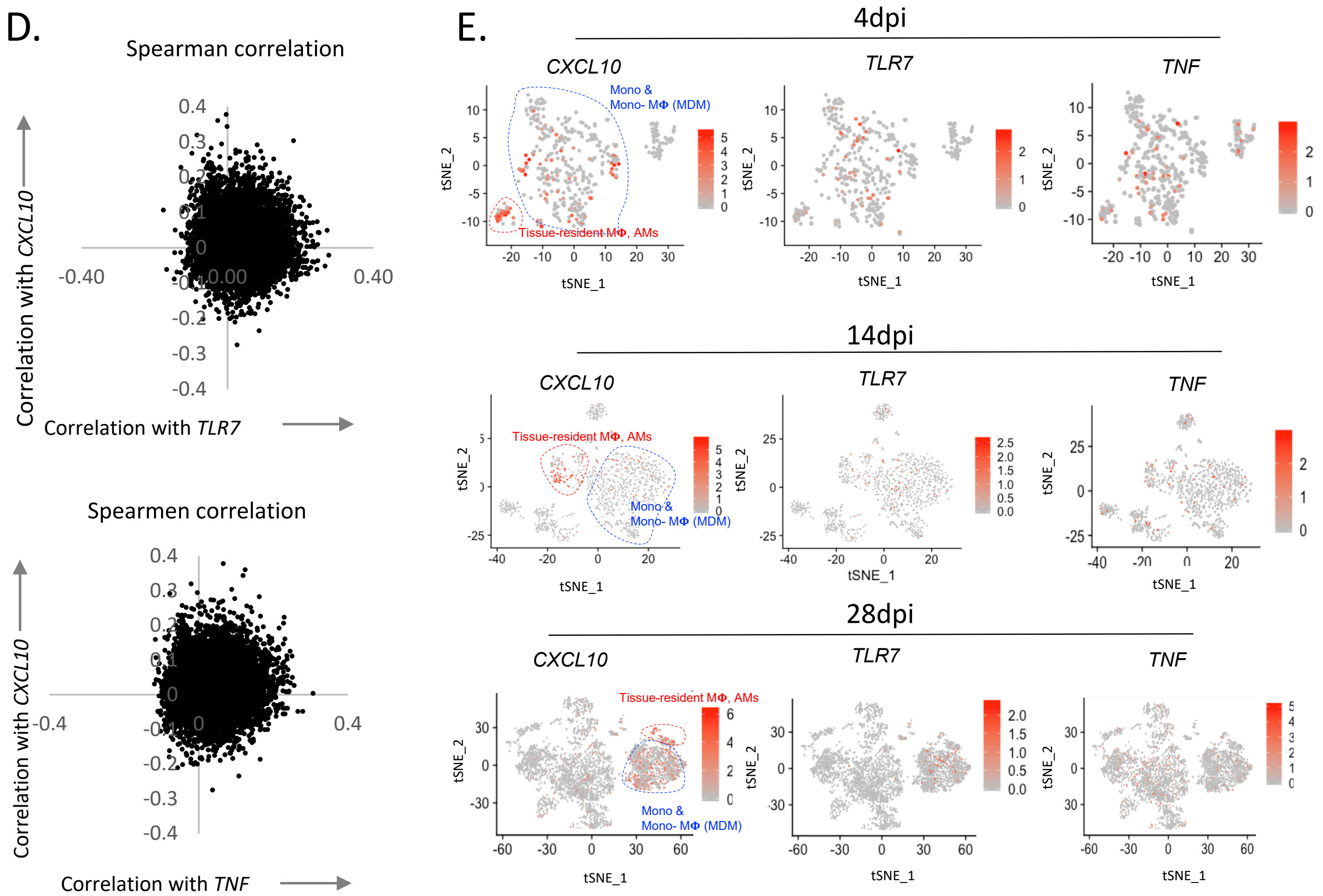

**Figure S3. SARS-CoV-2 infection of human macrophages activates inflammasomes and leads to a unique inflammatory transcriptional signature in vivo (matched to figure 3).**

D. Spearman correlation values of each gene based on its correlation with *CXCL10* or *TNF* or *TLR7*.

E. Expression and distribution of *CXCL10, TNF* and *TLR7* in human immune cells from infected (4, 14 and 28dpi) MISTRG6-hACE2 mice.

**F.** Top genes that correlate with *CXCL10* (4dpi, Pearson and Spearmen correlation combined) are enriched for distinct inflammatory molecules.

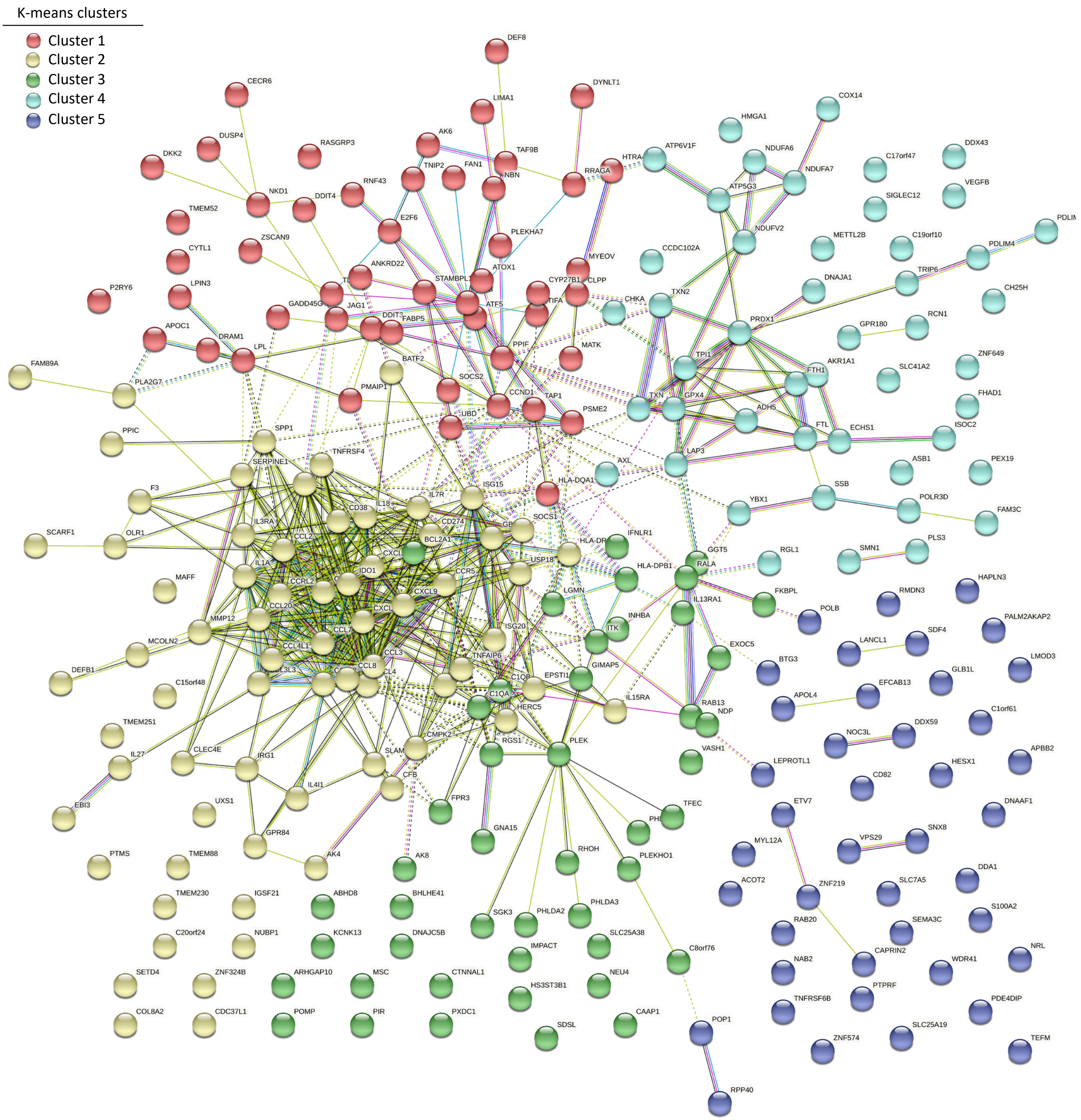

**Figure S3. SARS-CoV-2 infection of human macrophages activates inflammasomes and leads to a unique inflammatory transcriptional signature in vivo (matched to figure 3).**

F. Network (STRING v11.0) analysis of top *CXCL10*-associated genes. K-means clustering. Clusters and their corresponding pathway analysis are available as source files.

G. Top genes that correlate with *CXCL10* but not with *TLR7* and *TNF* are enriched for the same distinct inflammatory molecules and alveolar macrophages.

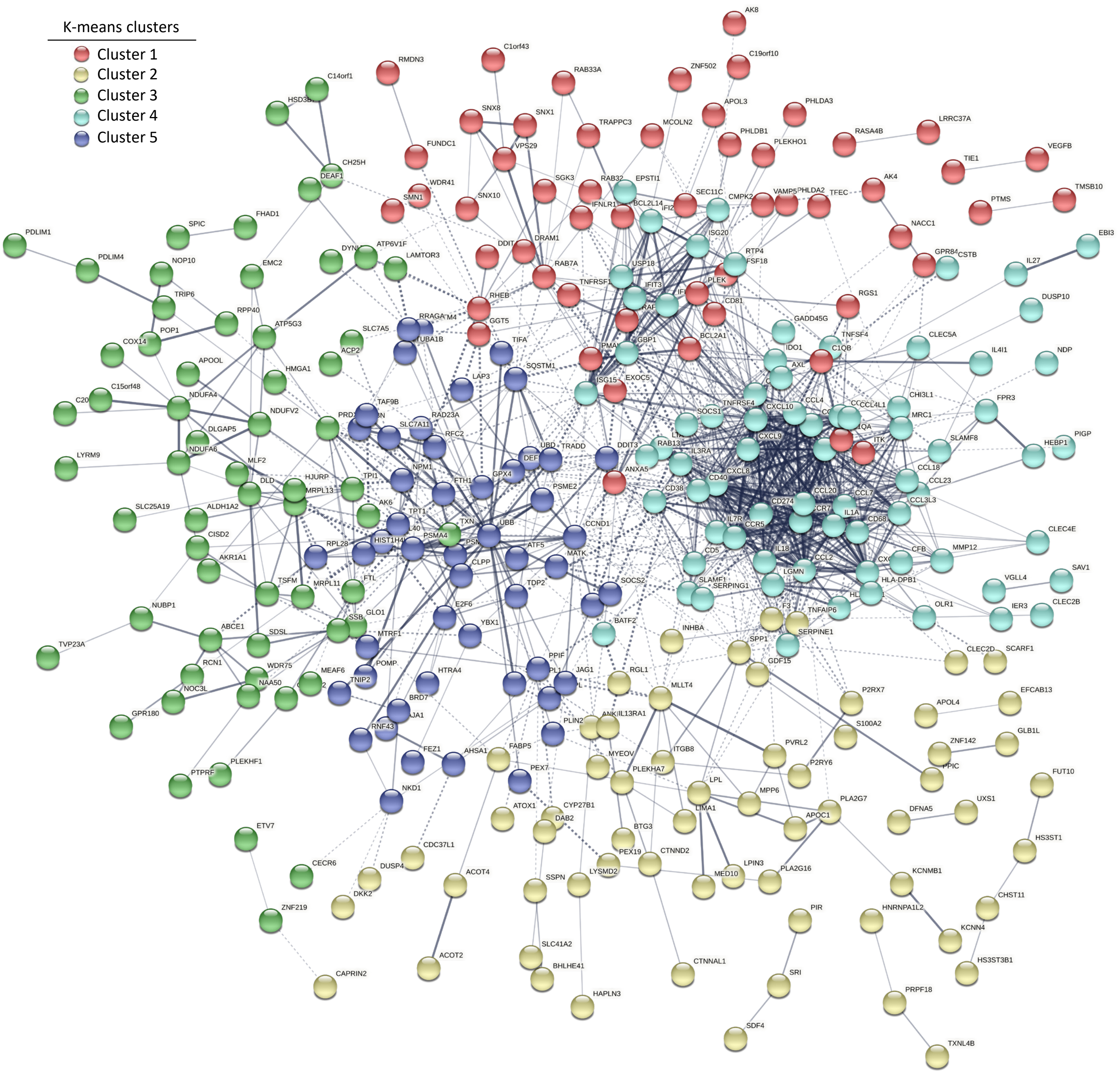

**Figure S3. SARS-CoV-2 infection of human macrophages activates inflammasomes and leads to a unique inflammatory transcriptional signature in vivo (matched to figure 3).**
G. Network (STRING) analysis of genes that are preferentially associated with CXCL10 but not with TLR7 or TNF. Disconnected nodes in the network are not displayed. K-means clustering. Clusters and their corresponding pathway analysis are presented as source files.

### Figure S3

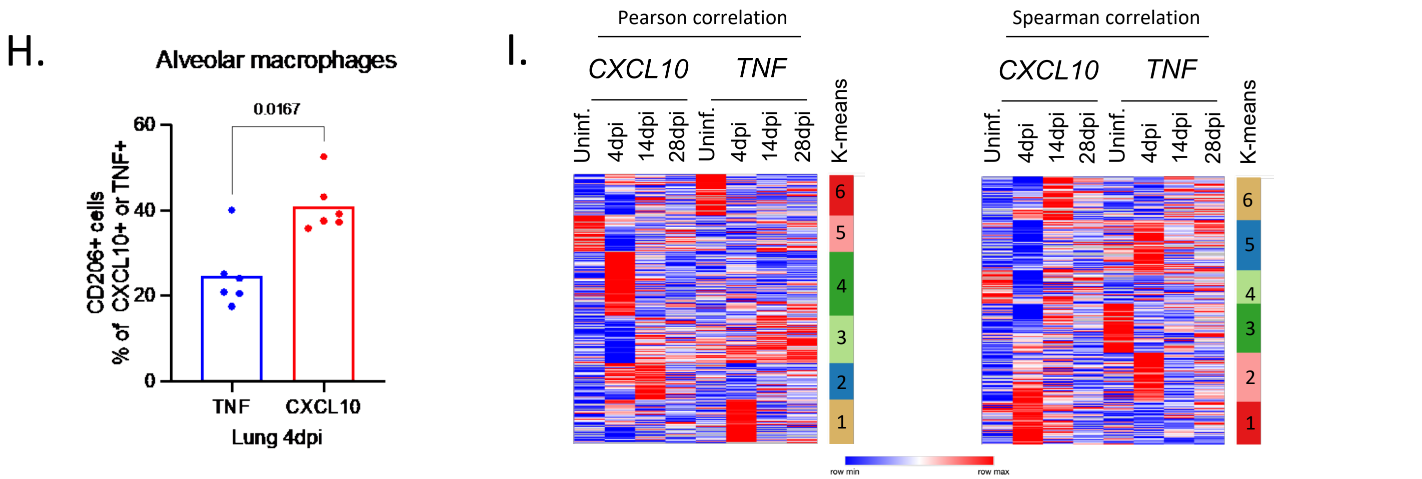

**Figure S3. SARS-CoV-2 infection of human macrophages activates inflammasomes and leads to a unique inflammatory transcriptional signature in vivo (matched to figure 3).**

H. Proportions of TNF or CXCL10 producing macrophages among alveolar ($CD206^{hi}CD68^{+}$) macrophages. Unpaired, two-tailed t-test. N=6 biologically independent mice examined over 3 independent experiments. MISTRG6-hACE2 mice were infected with SARS-CoV-2-mNG and lungs were analyzed at 4dpi.

I. Distribution of *CXCL10* or *TNF* associated genes at 4, 14, 28 dpi in lungs infected with SARS-CoV-2 or not. Analysis performed on macrophages of 4dpi lungs in Fig 4D was extended to more timepoints. Pearson (right) and Spearman (left) correlation values were calculated for each gene for its correlation with *CXCL10* or *TNF* in human monocytes and macrophages isolated from uninfected and infected (4, 14 and 28 dpi) lungs. K-means clustering analysis.

Figure S4

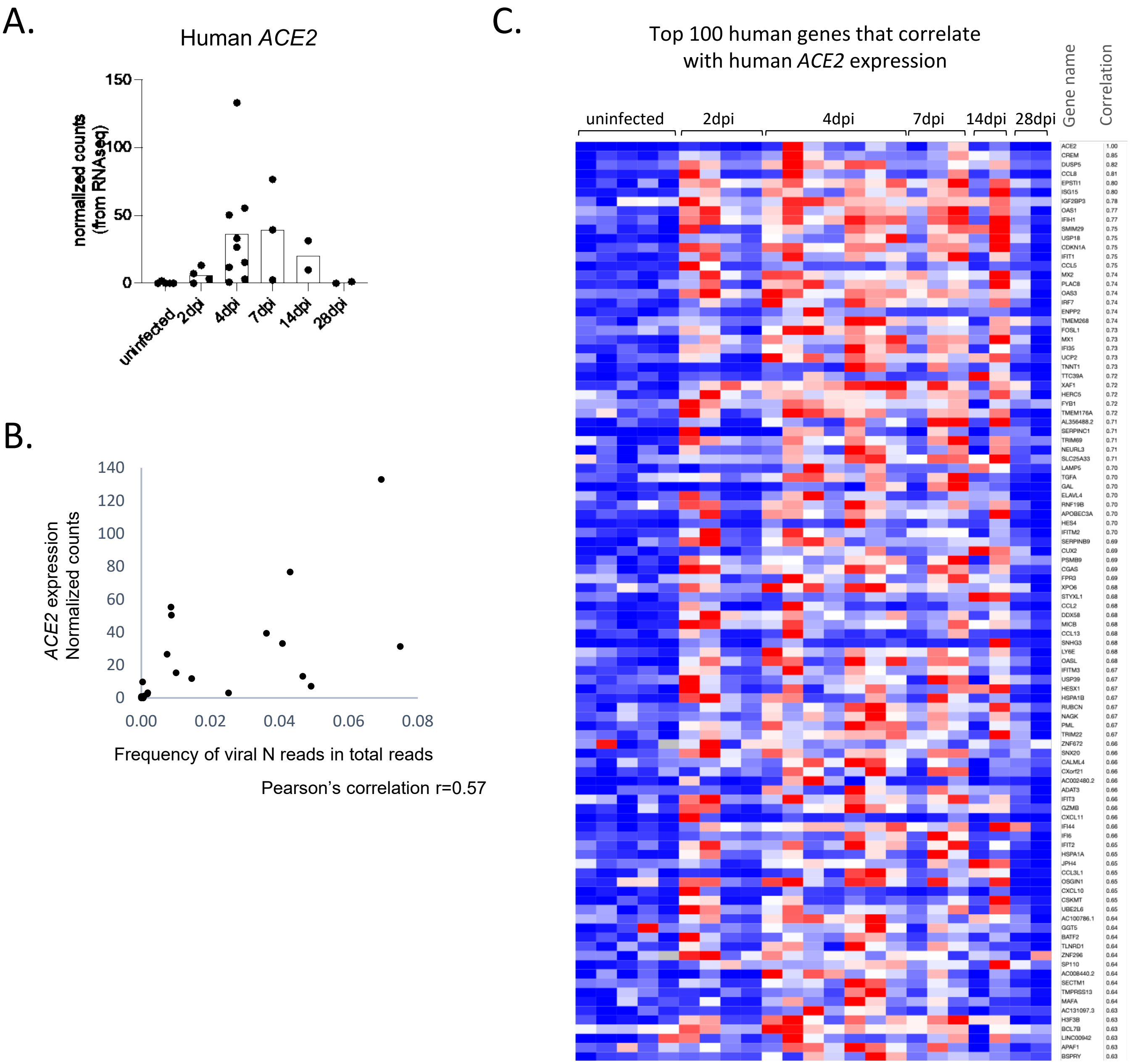

**Figure S4. *ACE2* expression is inducible and highly correlates with genes that are associated with type I interferon signaling.**

A. Normalized count for *ACE2* expression during the course of infection in whole homogenized lungs of MISTRG6-hACE2 infected with SARS-CoV-2 (GSE186794). We confirmed that the transcript measured here is the full length *ACE2*, not the non-functional truncated version named MIRb-ACE247. Uninfected n=5, 2dpi n=4, 4dpi =9, 7dpi n=3, 14dpi n=2, 28dpi n=2 biologically independent mice examined over at least 2 experiments.

B. Two-way plot showing normalized count for *ACE2* expression and the corresponding percent SARS-Cov-2 transcripts within total transcripts (murine, human, and viral) in lungs of SARS-CoV-2 infected MISTRG6-hACE2 mice. Pearson correlation value between *ACE2* and percent viral transcripts was calculated as 0.57. N=20 biologically independent mice examined over at least 2 independent experiments. 2,4,7,14,28 dpi pooled.

C. Top 100 human genes that correlate with *ACE2* expression in uninfected and infected lungs. Genes ranked based on correlation value that is displayed as a numerical column. Uninfected n=5, 2dpi n=4, 4dpi n=9, 7dpi n=3, 14dpi n=2, 28dpi n=2 over at least 2 experiments (GSE186794).

Figure S4

D.

| Gene Set Name [# Genes (K)] | Description | # Genes in Overlap (k) | k/K | p-value | FDR q-value |
|---|---|---|---|---|---|
| REACTOME_INTERFERON_SIGNALING [203] | Interferon Signaling | 21 | | $9.58\ e^{-29}$ | $2.75\ e^{-25}$ |
| REACTOME_INTERFERON_ALPHA_BETA_SIGNALIALING [71] | Interferon alpha/beta signaling | 16 | | $7.58\ e^{-28}$ | $1.09\ e^{-24}$ |
| REACTOME_CYTOKINE_SIGNALING_IN_IMMUNE_NE_SYSTEM [884] | Cytokine Signaling in Immune system | 29 | | $3.17\ e^{-25}$ | $3.03\ e^{-22}$ |
| REACTOME_ANTIVIRAL_MECHANISM_BY_IFN_ST_STIMULATED_GENES [82] | Antiviral mechanism by IFN-stimulated genes | 11 | | $8.91\ e^{-17}$ | $6.39\ e^{-14}$ |
| WP_TYPE_II_INTERFERON_SIGNALING_IFNG [37] | Type II interferon signaling (IFNG) | 6 | | $3.31\ e^{-10}$ | $1.9\ e^{-7}$ |
| WP_THE_HUMAN_IMMUNE_RESPONSE_TO_TUBERCERCULOSIS [23] | The human immune response to tuberculosis | 5 | | $2.19\ e^{-9}$ | $1.05\ e^{-6}$ |
| REACTOME_OAS_ANTIVIRAL_RESPONSE [9] | OAS antiviral response | 4 | | $3.7\ e^{-9}$ | $1.52\ e^{-6}$ |
| REACTOME_CHEMOKINE_RECEPTORS_BIND_CHEMHEMOKINES [58] | Chemokine receptors bind chemokines | 6 | | $5.54\ e^{-9}$ | $1.99\ e^{-6}$ |
| REACTOME_NEGATIVE_REGULATORS_OF_DDX58_58_IFIH1_SIGNALING [34] | Negative regulators of DDX58/IFIH1 signaling | 5 | | $1.77\ e^{-8}$ | $5.65\ e^{-6}$ |
| WP_CYTOSOLIC_DNASENSING_PATHWAY [74] | Cytosolic DNA-sensing pathway | 6 | | $2.46\ e^{-8}$ | $6.51\ e^{-6}$ |

**Figure S4. *ACE2* expression is inducible and highly correlates with genes that are associated with type I interferon signaling.**
D. Pathway enrichment analysis for genes highly correlating with *ACE2* expression (as shown in Fig S5C)

#### Supplementary Tables

**Table S1: Human genes that are differentially regulated in lungs of infected MISTRG6-hACE2 in response to therapeutics.**

Genes that are upregulated in response to infection and downregulated in response to therapeutics (dexamethasone, anti-IFNAR+ Remdesivir) in these infected mice at 14dpi were included in the analysis (matched to Fig 1d). Normalized expression of duplicates. N=2 biologically independent mice examined over 2 -independent experiments. Differential expression analysis was performed with DESeq2 and statistical significance was deemed using Wald test.

**Table S2: Cluster identifying markers and markers that identify temporal transcriptional changes associated with monocytes and macrophages in infected (4, 14 or 28dpi) or uninfected lungs of MISTRG6-hACE2 mice (matched to Fig 1g).**

N=2 biologically independent mice for each condition was pooled. Marker genes for each cluster of cells were identified using the Wilcoxon test with Seurat. For the adjusted P-values the Bonferroni correction was used.

**Table S3: Expression of human genes that are enriched in macrophages (clusters identified as part of Fig. 1g) during SARS-Cov-2 infection and their response to anti-IFNAR2 and Remdesivir therapy (matched to Fig. 1h). Normalized expression of duplicates analyzed.**

N=2 biologically independent mice examined over 2 independent experiments. Differential expression analysis was performed with DESeq2 and statistical significance was deemed using Wald test.

**Table S4: Pearson and spearman correlation values calculated for each gene for its correlation with CXCL10, TNF or TLR7 in human monocytes and macrophages at 4dpi (based on Fig. 1g, matched to Fig. 3d).**

For Pearson's test, significance was based on the t-test with statistic based on Pearson's product-moment correlation coefficient cor(x, y) and following a t distribution with length(x)-2 degrees of freedom. For Spearman's test, p-values are computed using algorithm AS 89 with exact = TRUE. Correlation values, p-values (two-tailed) and FDR-adjusted p-value are presented.

**Table S5: Patient specimens used for immunofluorescent (IF) staining.**

Details of patient demographics for specimens use in IF staining: Age, gender, medication, time of death post-symptom onset (dps), co-morbidities, cause of death and histopathological findings. This table is presented as part of supplementary methods.
